# Amplicon/Protein Bead Display enables quantitative *in vitro* biochemistry at scale

**DOI:** 10.64898/2026.05.28.728566

**Authors:** Daria R. Passow, Anvita Gupta, Samuel Thompson, Anshul Kundaje, Polly M. Fordyce

## Abstract

Scalable methods for producing and characterizing protein libraries are essential for generating standardized datasets needed to train models linking sequence to function. Here, we present Amplicon/Protein Bead Display (APB-Display), which expresses and purifies >100,000 protein variants *in vitro* in 1 day. APB-Display uses particle-templated emulsification to generate libraries of hydrogel beads that covalently display many copies of a given protein variant and its encoding DNA. By incubating beads with fluorescently-labeled ligands, sorting, and sequencing to generate titration curves (APB-TiteSeq), we simultaneously quantified expression levels and binding affinities (*K_d_*s) for >18,000 FLAG epitope variants interacting with M2 anti-FLAG antibody in 3 days, revealing chemistry-dependent epistasis between positions 4 and 5. Single-concentration binding measurements (APB-SortSeq) paired with neural network denoising further returned quantitative affinities for >88,000 variants. APB-Display requires only standard laboratory equipment and access to a FACS sorter, providing an accessible platform for quantitative *in vitro* biochemistry at scale.

## Introduction

Recent advances in deep learning have transformed our ability to predict protein structure from sequence [1–3] and design sequences that adopt a given structure [4, 5]. These advances were made possible by decades of accumulated sequence [6] and structural [7] data in standardized formats. Learning to predict interaction affinities and efficiently design selective protein binders will similarly require large, standardized datasets linking sequence to quantitative measurements of binding affinity and selectivity [8–11]. Ideally, these measurements would report absolute thermodynamic constants (*K_d_*s), allowing standardization of data across molecules and laboratories and prediction of molecular behavior under conditions never experimentally probed [12]. Moreover, measured sequences should be informed by model predictions to enable active learning cycles [13], which require fast experimental turnaround to be practical. Acquiring *K_d_* measurements at this scale and speed thus demands methods that combine the quantitative rigor of *in vitro* biochemistry with high throughput and broad accessibility.

Cell-based assays can survey large sequence spaces but introduce biological complexity that fundamentally limits their scope and ability to return thermodynamic constants. Cell-based multiplexed assays of variant effects (MAVEs [14]) link binding to fitness [15–21], fluorescence [22], or luminescence [23], yielding outputs that can correlate with *K_d_* and allow inference of free energy changes. However, these output parameters depend on both interaction affinities and protein abundance, complicating their interpretation [24–26]. While yeast surface display paired with FACS-based titration curves (Tite-Seq [27–30] and related methods [31, 32]) control for protein abundance and return *K_d_*s by directly quantifying binding of fluorescently-labeled ligands to surface-displayed variant libraries, these approaches bias libraries toward variants that are efficiently displayed on the cell surface and can be influenced by other cell-surface proteins [33]. More generally, all cell-based methods are restricted to sequences that can be efficiently translated, escape proteolysis, and are non-toxic. As a result, cellular fitness landscapes convolve molecular fitness properties with cell type-specific properties (*e.g.* expression of endogenous proteins that modulate variant abundance [34, 35], install post-translational modifications [36], or possess competing activities [37]).

*In vitro* assays can investigate binding of purified proteins under precisely-controlled buffer conditions to yield true interaction affinities (*K_d_*s). However, traditional biophysical approaches individually clone, express, purify, quantify, and assess binding of each variant to systematically increasing ligand concentrations, limiting practical throughput to tens (*e.g.* isothermal calorimetry, surface plasmon resonance, biolayer interferometry, and gel shift assays [12]) or hundreds (*e.g.* holdup assays [38, 39] and microfluidic approaches [40–42]) of protein variants per experiment. To increase throughput, *in vitro* display-based methods leverage advances in oligonucleotide and cell-free protein synthesis to enable pooled expression, purification, and binding characterization for larger protein libraries. Monovalent *in vitro* display formats (*e.g.* mRNA display [33, 43–46], cDNA display [47], ribosome display [48], and Cas9-display [49]) pan variant libraries against an immobilized ligand followed by sequencing bound and unbound fractions to yield enrichment scores that reflect relative binding preferences. While enrichment scores can in principle correlate with *K_d_* [43, 44], most applications report these scores without affinity calibration. Furthermore, these approaches typically assess binding at a single, unquantified ligand concentration and lose interactions with fast dissociation rates during required wash steps, limiting both assay dynamic range and the ability to detect transient interactions [12, 33]. Alternate approaches tether ribosome-displayed [50, 51] or Cas9-displayed [52] proteins to DNA microarray surfaces for fluorescence-based binding measurements but require specialized imaging infrastructure and have not returned absolute affinities.

Multivalent bead display offers a compelling but underexploited architecture for scalable *in vitro* affinity measurements. In a canonical bead display workflow [53], emulsion PCR of a pooled DNA library generates beads bearing clonal DNA populations; *in vitro* transcription/translation (IVTT) in a second set of emulsions expresses and captures the encoded proteins on those same beads, thereby producing libraries where each bead displays a single protein variant at high copy number. Subsequent incubation with fluorescently-labeled ligand makes it possible to directly estimate per-bead ligand occupancies via fluorescence-based sorting and sequencing without a need for individual molecule counting [54]. While prior platforms have used this workflow to screen large libraries of protein binders [54–61], characterization was limited to selecting top hits for downstream validation. Previous bead display approaches also tethered molecules to beads via non-covalent linkages that led to DNA loss and cross-talk throughout the workflow [60] and used rigid polystyrene beads with high levels of non-specific protein adsorption, limiting dynamic range [54]. Spectrally-encoded hydrogel beads demonstrated that the low nonspecific background of hydrogel matrices enables quantitative binding measurements (ΔΔGs) [62, 63], but these assays were limited to hundreds of variants.

Here, we present Amplicon/Protein Bead Display (APB-Display), a fully *in vitro*, hydrogel bead-based platform for pooled expression and purification of large protein libraries in less than 8 hours using only standard laboratory equipment. To validate the ability to quantify expression and interaction affinities using Amplicon/Protein Beads (APBs), we profiled interactions between variants of the 8-residue FLAG epitope and the M2 anti-FLAG antibody, a well-characterized system with a co-crystal structure [46, 49, 50, 52, 64–66]. We systematically assessed how sequence variation impacts expression levels, revealing substantial position- and residue-specific effects for even a simple linear peptide. By coupling APB-Display with multi-bin FACS sorting and deep sequencing across a ligand concentration series (APB-TiteSeq), we demonstrated parallelized measurements of absolute *K_d_* for >18,000 FLAG epitope variants in a single three-day experiment at low cost ($0.15 per variant) and with an energetic resolution of <1.4-fold (0.3 kT). By fitting models of increasing complexity to these data, we dissected the structure of the resulting sequence-affinity landscape to reveal chemistry-dependent epistasis between residues 4 and 5. Finally, we demonstrated that single-concentration measurements (APB-SortSeq) paired with neural network denoising extend quantitative affinity recovery to >88,000 variants. Together, APB-Display and its extensions provide a platform for quantitative *in vitro* biochemistry at the scale and speed needed to train predictive models linking protein sequence to function.

## Results

### APB-Display pipeline enables high-throughput protein expression, purification, and characterization

Amplicon/Protein Bead Display (APB-Display) recombinantly expresses and purifies large protein libraries in less than 8 hours by leveraging particle-templated emulsification, emulsion PCR, and emulsion IVTT (Fig. 1a). Downstream characterization assays paired with sequencing-based readouts enable diverse applications; here, we demonstrate direct quantification of expression levels and absolute binding affinities by incubating APB libraries with fluorescently-labeled targets, FACS sorting, and deep sequencing (APB-TiteSeq) (Fig. 1b).

**Figure 1:**
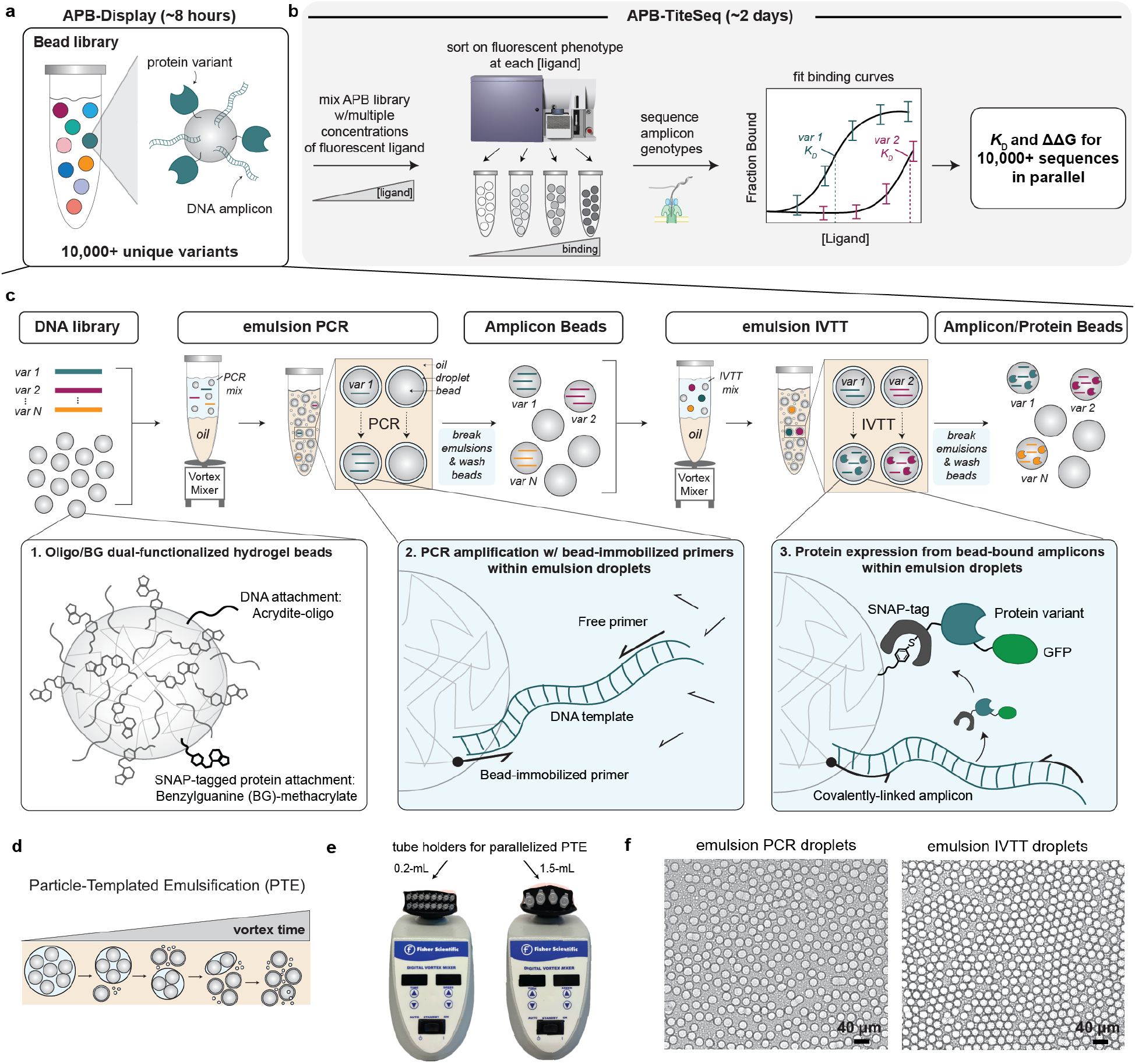
Overview of the APB-Display workflow. **a.** Schematic illustrating that APB-Display generates libraries of bead particles displaying many copies of a DNA amplicon and the protein it encodes. **b.** APB-TiteSeq workflow for quantifying expression and binding affinities at scale. **c.** Detailed schematic of workflow used to generate APBs. Dual-functionalized polyacrylamide hydrogel beads bearing covalently-attached reverse primers and benzylguanine (BG) groups are encapsulated with reagents for emulsion PCR (generating Amplicon Beads) and emulsion IVTT (generating Amplicon/Protein Beads) via particle-templated emulsification (PTE). Breaking emulsions and washing between steps and after emulsion IVTT facilitates reagent exchange and purification, respectively. **d.** Schematic illustrating the principle of water-in-oil droplet formation via vortexing during PTE. **e.** Images showing foam tube holders mounted on standard vortex mixers for parallelized PTE. **f.** Representative brightfield images of APBs in particle-templated emulsions after emulsion PCR (left) and emulsion IVTT (right).

The APB-Display workflow starts with a pooled DNA library encoding SNAP- and GFP-tagged protein variants and polyacrylamide hydrogel beads bearing covalently-coupled oligonucleotides and benzylguanine (BG) groups for DNA and protein attachment, respectively (Fig. 1c) [67]. Here, we generated dual-functionalized, ∼ 18-micron diameter hydrogel beads using a simple flow-focusing microfluidic device (Supplementary Figures 1 and 2). Once produced in bulk, bead stocks were stored and used across multiple APB-Display experiments.

To generate libraries, we encapsulate beads with PCR reagents and DNA templates in uniform aqueous-in-oil droplets via particle-templated emulsification (PTE) [68] using foam tube holders affixed to a standard vortex mixer (Fig. 1d-f). The use of a dilute DNA concentration (Poisson λ ∼ 0.1) ensures that most droplets contain zero or one template molecule, such that PCR amplification yields beads displaying either no DNA or many copies of a single variant (Amplicon Beads; Fig. 1c). Next, we break emulsions, wash, and re-encapsulate Amplicon Beads with IVTT reagents via a second round of PTE. During the IVTT reaction, the N-terminal SNAP-tag on expressed proteins drives covalent coupling to bead-attached BG groups, while the C-terminal GFP tag makes it possible to quantify expression. Breaking emulsions and washing yields Amplicon/Protein Bead (APB) libraries displaying clonal populations of purified proteins and the DNA amplicons that encoded them.

### APB-Display generates bead libraries with 1:1 genotype-phenotype linkages

Accurately quantifying sequence/function relationships via APB-Display requires that particles reliably link genotypes and phenotypes by displaying clonal DNA amplicons and clonal populations of the corresponding functional, full-length proteins without substantial mixing or cross-talk. To test for this, we generated a 25-variant library of SNAP- and eGFP-tagged FLAG epitopes (wildtype DYKDDDDK plus A, L, and E substitutions at each position) using APB-Display (Fig. 2a) and quantified both expression and binding to a 1:1 mixture of the M2 anti-FLAG antibody and a Cy5-labeled secondary antibody (Fig. 2b).

**Figure 2:**
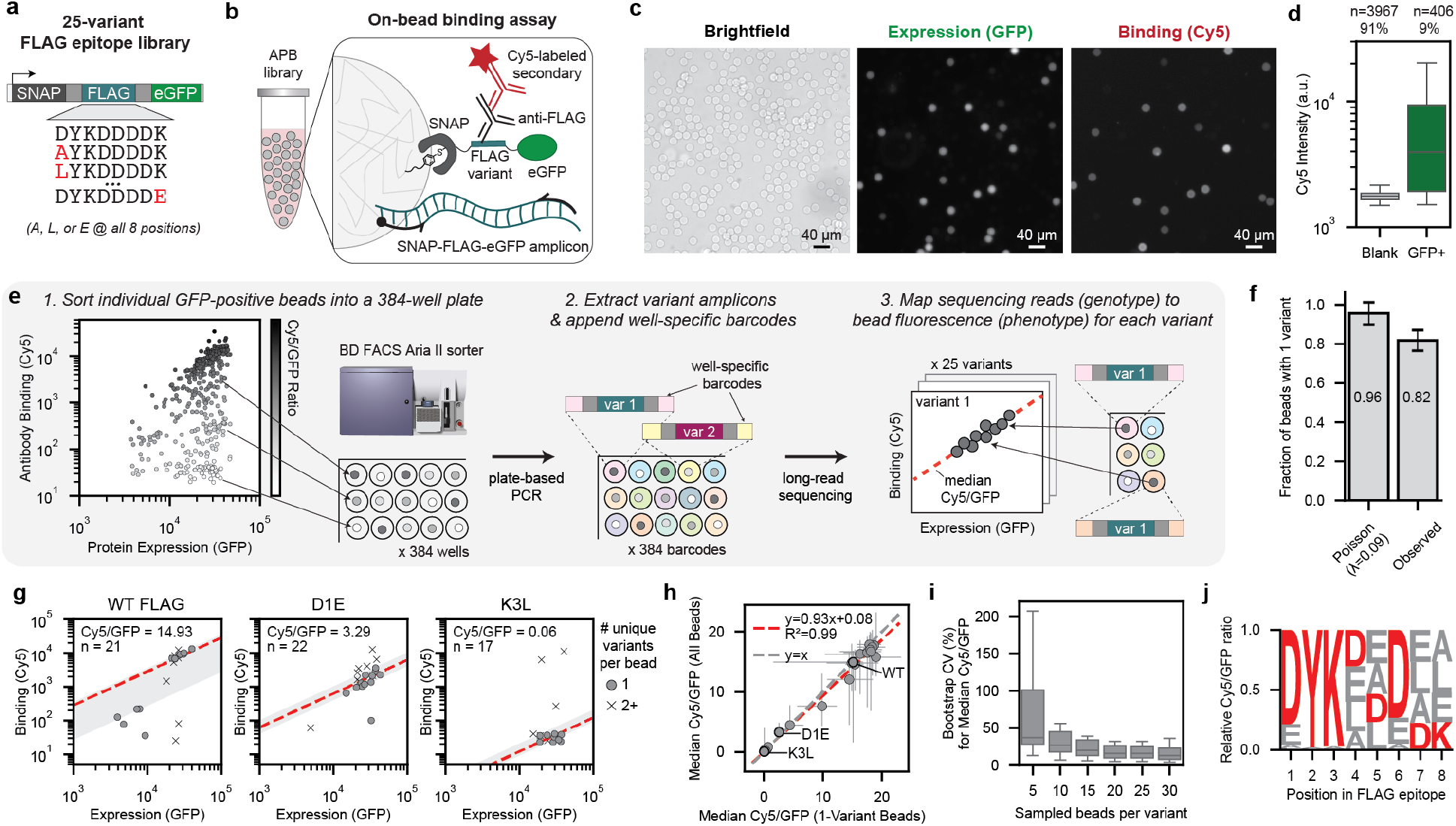
APB-Display generates protein libraries with expected genotype-phenotype linkages. **a.** Schematic of DNA templates encoding expression of a 25-variant FLAG library expressed as SNAP and eGFP fusion proteins on APBs. **b.** Schematic of experiments to detect binding between bead-displayed FLAG epitopes and M2 anti-FLAG antibodies using a Cy5-labeled secondary antibody. **c.** Representative brightfield and fluorescence images of APBs after incubation with M2 and Cy5-labeled secondary antibodies reporting on per-bead expression levels (GFP intensities), and antibody binding (Cy5 intensities). **d.** Box plots of per-bead median Cy5 pixel intensities (arbitrary units) for blank and GFP-positive beads. **e.** Schematic of single-bead sorting and sequencing workflow for validating genotype-phenotype linkages. FACS plot (far left) displays measured GFP (x-axis) and Cy5 (y-axis) intensities for the 384 GFP-positive beads sorted into a multi-well plate. **f.** Bar plots of expected (based on Poisson loading at λ = 0.1) and observed fractions of single-variant beads in the GFP-positive library. Error bars represent the Poisson-derived standard deviation. **g.** Measured Cy5 versus eGFP fluorescence intensities for all beads displaying three representative FLAG variants. Circles indicate single-variant beads; X’s indicate multi-variant beads. Red dashed lines indicate median Cy5/GFP ratio; gray shading shows interquartile range [0.25,0.75]. **h.** Median Cy5/GFP ratios calculated from all beads bearing a given variant versus single-variant beads only. Error bars denote interquartile range [0.25, 0.75]. **i.** Coefficient of variation (CV%) of median Cy5/GFP ratios across 1,000 bootstrap samples as a function of beads sampled per variant. **j.** Sequence logo showing relative position-normalized Cy5/GFP ratios for single amino acid substitutions (A, L, or E) at each position in the FLAG epitope. Red letters indicate wildtype residues; grey letters indicate substitutions.

#### Amplicon Beads show clonal DNA amplification

Under Poisson loading statistics, we expect that a DNA concentration corresponding to λ=0.1 should yield a droplet population in which 90% of droplets contain no DNA; of the 10% of droplets loaded with DNA, most (95%) should contain a single molecule (Supplementary Figure 3). To empirically identify the DNA concentration corresponding to this regime, we performed digital droplet PCR in vortexed emulsions under conditions that yielded droplets of similar size as our PTE-generated emulsions (median diameter = 11.6 μm, Supplementary Figure 4). Successful amplification of DNA within each droplet generated a fluorescence signal, making it possible to quantify the fraction of fluorescent droplets via bright field and fluorescence microscopy.

As expected, the fraction of fluorescence-positive droplets increased linearly with increasing DNA concentration; a linear fit to this calibration curve predicted that a DNA concentration of ∼ 500 fM should yield λ=0.1 (Supplementary Figure 4).

#### Amplicon Beads drive functional protein expression

Next, we used ∼ 500 fM of the 25-variant DNA library to produce APBs and test if full-length and properly folded proteins could be produced from bead-attached templates via emulsion IVTT. To simultaneously assess binding specificity, we added 6 nM each of M2 anti-FLAG and Cy5-labeled secondary antibody to the APB library, incubated overnight to ensure equilibrium within the hydrogel bead matrix (Supplementary Figure 5)[63], then imaged in GFP and Cy5 fluorescence channels (Fig. 2c). As expected for Poisson loading at λ=0.1, 9.3% of beads displayed strong GFP fluorescence after washing (Fig. 2d, Supplementary Figure 6). The presence of bead-associated GFP intensities established that expressed proteins were properly folded and full-length, as this requires a functional SNAP tag (to drive attachment to the bead) and GFP tag (to emit fluorescence). SNAP-FLAG-eGFP proteins incubated with beads lacking BG showed no detectable fluorescence (Supplementary Figure 7), confirming that SNAP-BG interactions are required for immobilization. To estimate the number of immobilized protein molecules per GFP-positive bead, we compared observed intensities with a droplet-based calibration curve (Supplementary Figures 8 and 9), yielding an estimate of ∼800 thousand immobilized protein molecules per GFP-positive bead (Supplementary Figure 10). This estimate guides experimental conditions (e.g. bead counts and antibody concentrations per reaction volume) that avoid ligand depletion when quantifying affinities (Supplementary Figure 11).

#### Amplicon/Protein Beads maintain genotype-phenotype linkages

Microscopy images confirmed that only GFP-positive APBs bound Cy5-labeled antibody (Fig. 2c-d), in line with previous reports that hydrogel beads minimize nonspecific interactions [62, 63, 67, 69]; additional controls assessing antibody binding to beads bearing SNAP-eGFP (with no FLAG epitope) showed minimal Cy5 fluorescence (Supplementary Figure 7), confirming that the antibodies do not bind SNAP or eGFP tags. To assess the fidelity of genotype/phenotype linkages, we used FACS to deposit 384 individual GFP-positive beads into a multi-well plate, recording both GFP intensity (expressed protein) and Cy5 intensity (bound antibody) intensities for each bead (Fig. 2e, Supplementary Figure 12). We then amplified DNA from each bead using with well-specific barcodes, pooled these barcoded amplicons, and identified variant sequences within each well via long-read sequencing (Supplementary Table 1). Of the 384 total wells, 296 (77%) yielded ≥ 25 reads corresponding to FLAG library variants (Supplementary Figure 13), covering all 25 library members with 8-23 beads each (Supplementary Figure 14). Read counts ranged from 25-1635 per bead (median = 602 counts, Supplementary Figure 15) and correlated only weakly with GFP fluorescence (*R*^2^=0.16; Supplementary Figure 16), suggesting that factors besides DNA amplicon concentration determine protein expression levels. Most (82%) of these beads contained a single FLAG variant, consistent with the 95% Poisson expectation at λ=0.1 (Fig. 2f).

APBs displaying a given FLAG variant bound the M2 antibody at similar strengths, visible as tight clusters along a given Cy5/GFP (binding/expression) ratio axis (Fig. 2g, Supplementary Figures 17 and 18). WT FLAG beads showed tighter binding (higher median Cy5/GFP ratios) than variants with core motif substitutions, confirming robust genotype-phenotype coupling. Median Cy5/GFP ratios for single-variant beads versus all (single-variant + multi-variant) beads were highly correlated (*R*^2^=0.99; Fig. 2h), indicating that, given sufficient oversampling, a low percentage of multi-variant beads does not substantially bias affinity estimates despite increasing variance (Supplementary Figure 19). Bootstrap analysis of median Cy5/GFP ratios for down-sampled APB populations established that 10 beads per variant yields a median coefficient of variation (CV) of 26%, with diminishing returns above 20 beads (Fig. 2i, Supplementary Figures 20 to 22).

### APB-derived binding strengths correlate with literature measurements

Computing per-position normalized median Cy5/GFP ratios across substitution variants (A, L, or E at each position) and visualizing this matrix as a logo [70] recovered the characteristic DYKxxDxx binding motif for the M2 antibody (Fig. 2j) [46, 49, 50, 52, 64, 65]. Median Cy5/GFP ratios from APB-Display correlated strongly with binding scores from two Cas9-display methods (*R*^2^=0.89 and 0.88; Supplementary Figure 23) and showed substantially improved dynamic range relative to a ribosome-display method (Supplementary Figure 24), establishing that 1:1 genotype-phenotype linkages generated by APB-Display enable accurate measurement of binding strengths.

### APB-TiteSeq quantifies absolute binding affinities at scale

To measure absolute binding affinities at scale, we extended APB-Display with a multi-bin sorting and sequencing strategy for generating parallelized *in vitro* titration curves (“APB-TiteSeq”, analogous to cell-based Tite-Seq methods [27–30, 71]). To demonstrate APB-TiteSeq, we generated a larger “single-variant” FLAG variant library that included WT and all 152 single amino acid substitutions flanked by SNAP and eGFP tags (Fig. 3a). Next, we split this library into 7 fractions and incubated them overnight with increasing concentrations (0.6-400 nM) of a 1:1 mixture of M2 primary antibody and Cy5-labeled secondary antibody. We then used FACS to sort all GFP-positive beads (25% of the total bead population) at each concentration into four bins based on their Cy5/GFP ratio (∼30,000 sorted beads over ∼3 minutes per concentration, ∼500 beads/variant, ∼200 beads/second; Supplementary Figures 25 and 26). We adjusted log-spaced bins to maintain roughly equal numbers of beads within each bin and set the value associated with each bin to its median Cy5/GFP ratio (dashed lines, Fig. 3b). Finally, we quantified variant frequencies within each bin by amplifying variant sequences from beads via PCR with with bin- and concentration-specific barcodes, pooling proportionally to bead counts, and counting amplicons via long-read sequencing (Supplementary Figure 27 and Supplementary Table 1). Input and sorted output variant frequencies were strongly correlated for 151/152 substitution variants (Fig. 3c), confirming minimal selection bias during library processing. The remaining variant (D4G) was underrepresented 60-fold in the input library and excluded from further analysis.

**Figure 3:**
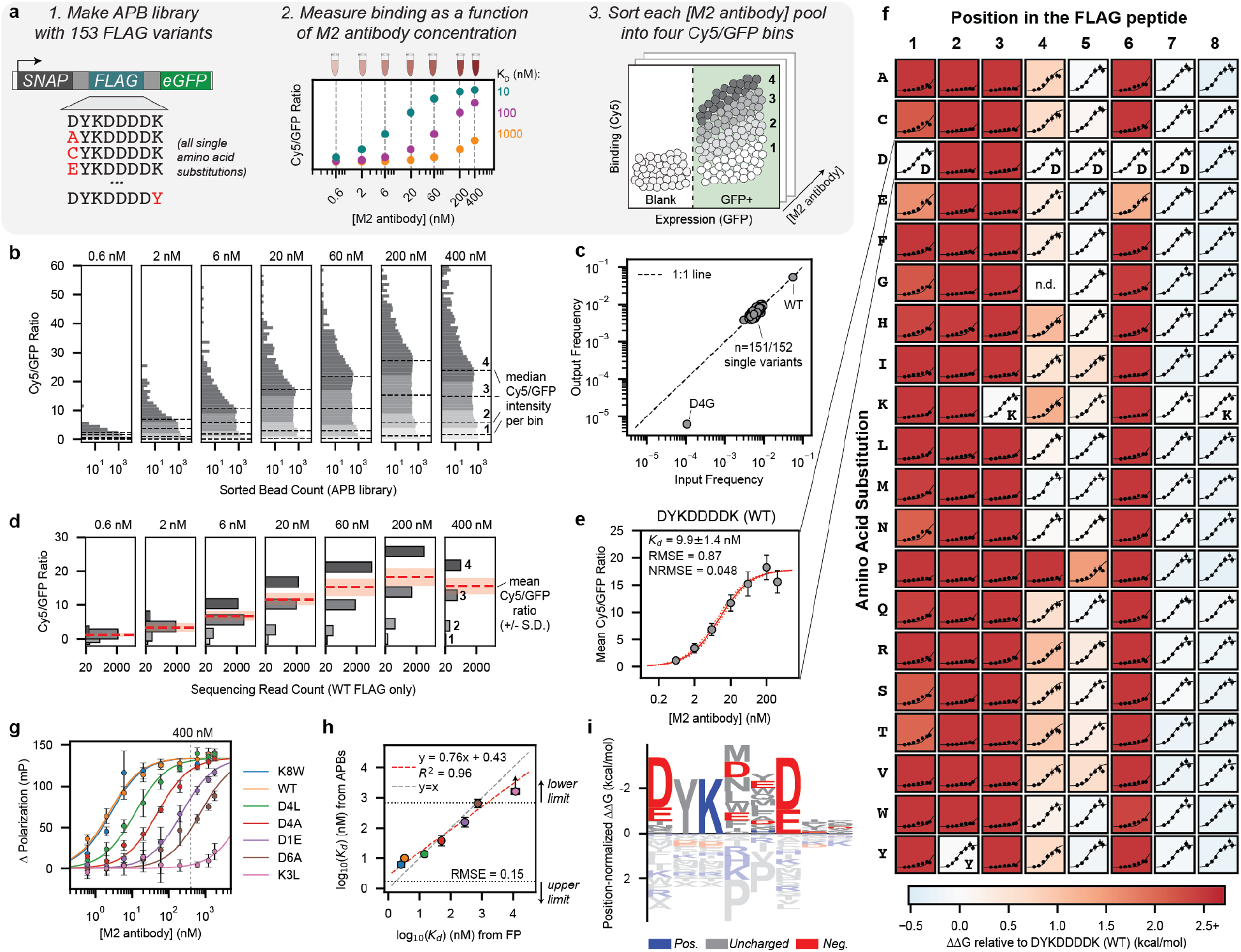
APB-TiteSeq quantifies equilibrium dissociation constants at scale with parallelized titration curves. **a.** (1) Single-variant FLAG library (WT plus all single amino acid substitutions) for APB generation. (2) The APB library is incubated with a series of M2 antibody concentrations, with each concentration pool sorted separately. (3) Representative FACS plots at each concentration with four sort gates along the Cy5/GFP axis. **b.** Histograms of Cy5/GFP ratios for sorted beads at each concentration. Colors denote sort bins; dashed lines indicate median Cy5/GFP values per bin. **c.** Variant frequencies in input DNA library versus output sorted bead library. **d.** Sequencing read counts for WT FLAG across bins and concentrations. Mean Cy5/GFP values (red dashed line ± SD) are computed as read-count-weighted averages of median bin Cy5/GFP values at each concentration. **e.** Binding curve for WT FLAG from mean Cy5/GFP values (± SD). Apparent *K_d_*extracted from Langmuir isotherm fit (solid red line). Dotted red lines: binding curves corresponding to fitted *K_d_*± SD. **f.** Heatmap of ΔΔ*G* values for all single-amino acid variants relative to WT FLAG (blue: more favorable binding; red: less favorable). Overlaid binding curves show fits to mean Cy5/GFP ratios (± SD) versus antibody concentration. ΔΔ*G* values are bounded within simulated measurement limits. **g.** Binding curves for seven FLAG variants measured via fluorescence polarization (mean ± SD of 3 technical replicates). Black dotted line: highest staining concentration used for APB-TiteSeq experiments (400 nM). **h.** Comparison of log_10_(*K_d_*) values from fluorescence polarization (± SD of technical replicates) and APB-TiteSeq (± fit SD). Marker colors match panel g. Black dotted lines and corresponding arrows denote upper and lower limits. Gray dashed line indicates 1:1 line; red dashed line indicates unweighted linear regression to points not indicated as limits. **i.** Sequence logo of ΔΔ*G* values for single amino acid substitutions normalized to the mean ΔΔ*G* at each position. Letter heights represent position-normalized ΔΔ*G*, with taller letters indicating larger effect sizes. Color indicates charge property under physiological conditions.

#### Binding isotherms can be constructed from binned sequencing data

To estimate binding affinities, we first calculated per-variant binding strengths (Cy5/GFP ratios) at each concentration by mapping sequencing read counts to fluorescence intensities (Fig. 3d). The mean Cy5/GFP ratio at each concentration was calculated as a weighted average of sort bins:

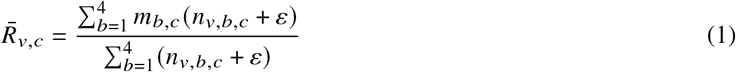

where 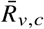 is the mean Cy5/GFP ratio for variant *v* at concentration *c, m_b,c_* is the median Cy5/GFP value of sort bin *b* at concentration *c, n_v,b,c_* is the normalized read count for variant *v* in bin *b* at concentration *c*, and ε = 0.1 is a pseudocount added to stabilize estimates for low-count variants. To estimate the strongest and weakest affinities that could be reliably detected under these experimental conditions, we simulated the full APB-TiteSeq experiment and data analysis pipeline and assessed agreement between returned fit parameters and input ‘ground truth’ values (Supplementary Figure 28 and Supplementary Tables 2 and 3). For log-uniformly-distributed affinities between 0.5-50,000 nM and experimentally-defined parameters (*e.g.* antibody concentrations spanning 0.6-400 nM, measured DNA loading rates, sort bin boundaries, bead counts, and estimated noise; Supplementary Figure 29), returned parameters began to deviate from ‘true’ values (>25% outside thermal noise threshold of 1 kT) for *K_d_*s tighter than 1.7 nM or weaker than 650 nM (Supplementary Figure 30); we used these values to set lower and upper limits on resolvable affinities from this particular experiment.

To estimate an apparent *K_d_* for each variant, we globally fit 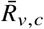 as a function of antibody concentration to a Langmuir isotherm, assuming 1:1 stoichiometry and a shared maximum signal (*R*_max_):

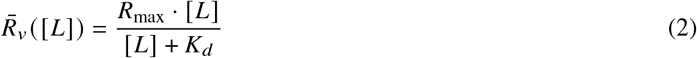

where 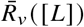 is the mean Cy5/GFP ratio for variant *v* at ligand concentration *L* and *K_d_* is the dissociation constant (ligand concentration at half-maximal response). As expected for a Langmuir isotherm, mean Cy5/GFP ratios for the WT FLAG peptide and many non-deleterious variants increased with antibody concentration until saturation (Fig. 3d-f, Supplementary Figures 31 to 38). Binding data were well-described by the Langmuir isotherm across all variants: the median normalized RMSE (NRMSE, defined as RMSE/*R*_max_) was 6.4% (10th percentile: 4.5%, 90th percentile: 9.8%; Supplementary Figure 39). Measured *K_d_*s followed a bimodal distribution (Supplementary Figure 40), with peaks near 10 nM for non-deleterious substitutions and >650 nM for core-motif substitutions.

#### APB-TiteSeq affinities agree with measurements via orthogonal solution-phase techniques

An assay designed to quantify affinities in high throughput must return accurate measurements. To experimentally assess APB-TiteSeq accuracy, we used an orthogonal solution-phase method (fluorescence polarization, FP) to quantify concentration-dependent binding for 7 peptides spanning the fitted affinity range of the single-variant library (WT and D1E, K3L, D4A, D4L, D6A, K8W variants; Fig. 3g). Log-transformed affinities (*K_d_*s) from globally fitting FP binding curves were well-correlated with APB-TiteSeq values (*R*^2^=0.96, RMSE=0.15; Fig. 3h). FP returned a quantitative estimate of 11.8 µM for the K3L peptide while APB-TiteSeq returned a lower limit estimate of 650 nM under the concentrations used here; our simulations accurately predicted this effect (Supplementary Figure 41).

#### Measured impacts of single-residue substitutions at each position yield a simple model of the affinity landscape

Position-specific affinity matrices (PSAMs) provide a simple model that predicts the expected energetic impact of substitutions at each residue on binding affinity, assuming that each residue contributes independently and additively to binding [72]. Here, we constructed a PSAM by computing the relative energetic impacts (ΔΔ*G*s) of each single substitution relative to WT FLAG (Fig. 3f). We also generated a logo visualization from position-normalized ΔΔ*G*s that reveals the characteristic DYKxxDxx binding motif [46, 49, 50, 52, 64, 65] while also detailing the charge properties of amino acids favored and disfavored at each position (Fig. 3i). Consistent with prior preference maps [46, 49, 50, 52], mutations to positions 1, 2, 3, and 6 have the largest impacts on affinity. At positions 1 and 6, charge-conservative substitutions (D1E and D6E) cause modest (∼1.5 kcal/mol) reductions in affinity, with non-conservative substitutions causing greater reductions. At positions 2 and 3, all substitutions disrupt binding by at least 2.5 kcal/mol, consistent with their key antibody contacts [64]. At position 4, positively charged residues (R, K, H) are disfavored by ∼ 1 kcal/mol, consistent with disruption of the D4-R32 salt bridge [64]. While positions 4 and 5 tolerate most substitutions, prolines are disfavored by >2.5 and 1.5 kcal/mol, respectively, revealing sensitivity to conformational change in the middle of the epitope. Consistent with their lack of antibody contacts, non-motif positions 5, 7, and 8 are highly tolerant to mutations (±0.5 kcal/mol), with the exception of proline at position 5.

### APB-TiteSeq scales to quantify 18,000 binding affinities within a week

Single amino acid substitutions cannot reveal epistatic interactions [73, 74] or provide the combinatorial sequence diversity needed to train predictive models [10]. Both tasks require multi-site mutant libraries; however, library size grows exponentially with mutation number, such that quantifying impacts of multi-site mutations requires scalable strategies for high-throughput experimental characterization. To demonstrate that APB-Display can generate, purify, and characterize large multi-site libraries, we designed a 23,283-member FLAG variant library consisting of two sub-libraries: all pairwise substitutions (including substitutions that generate premature TAG stop codons) and quadruple substitutions at positions 1, 4, 5, and 7 (Fig. 4a). After assembling this “four-position” library via cell-free Golden Gate assembly and PCR purification (2 days) and generating APBs (9% GFP-positive, 1 day), we again performed APB-TiteSeq at six M2 antibody concentrations (0.6-400 nM) to quantify protein expression and binding affinities. Overall, we sorted ∼ 300,000 beads per concentration in 9 hours (∼20 beads/variant, 45 minutes per sort; Fig. 4b, Supplementary Figures 42 to 45). After sorting, we barcoded each sorted bead population for pooled long-read sequencing (0.5 days; Supplementary Figures 46 and 47 and Supplementary Table 1).

**Figure 4:**
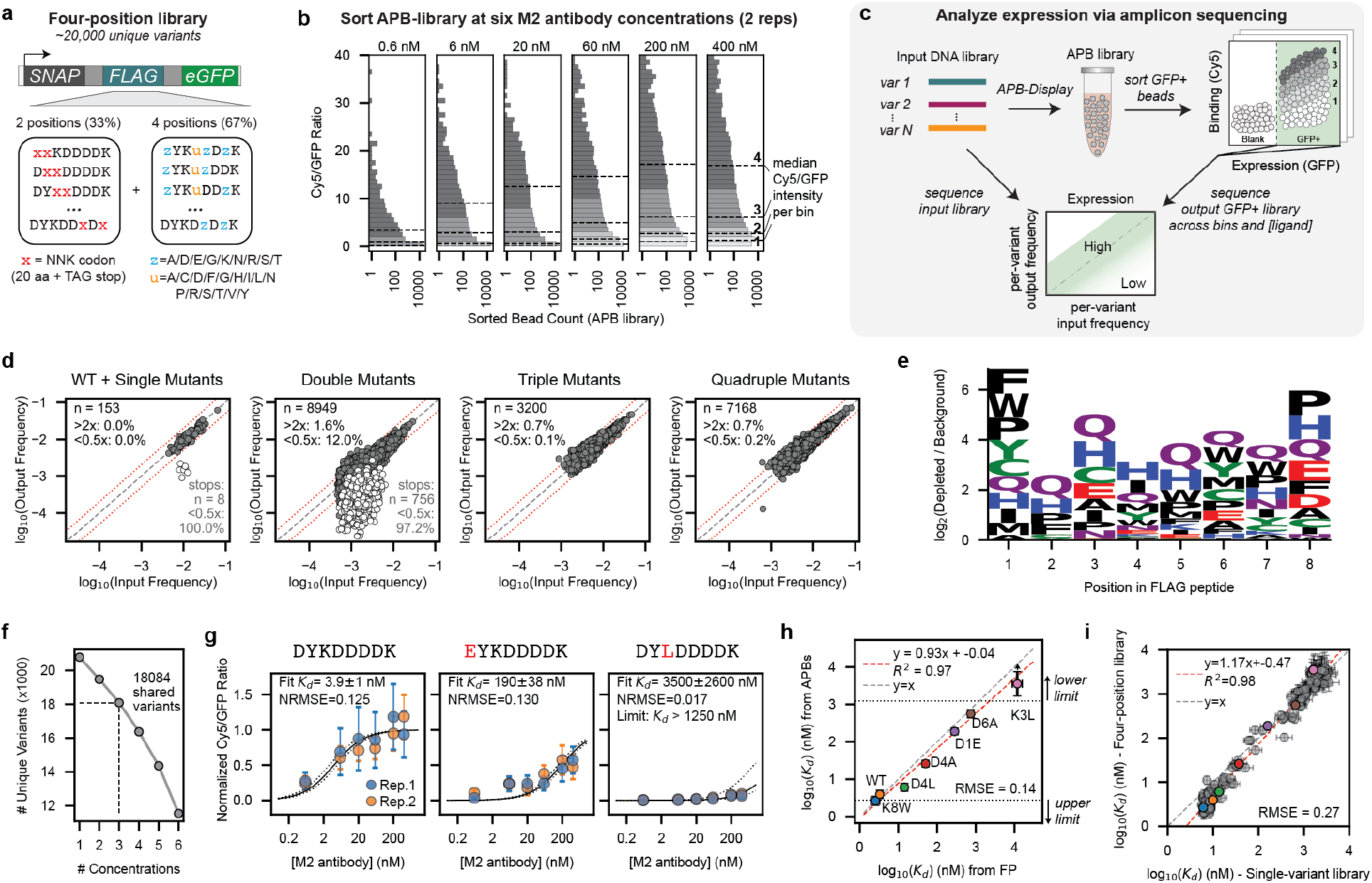
APB-TiteSeq scales to 18,000-variant libraries. **a.** Schematic of four-position library with all pairwise substitutions (including stop codons) and quadruple substitutions at positions 1, 4, 5, 7. **b.** Measured Cy5/GFP distributions at each antibody concentration. Bars are colored by sort bin; dashed lines indicate median Cy5/GFP ratios for each bin. **c.** Schematic of expression screening experiment in which comparing output/input variant frequency ratios identifies variants with reduced expression or truncated products. **d.** Log_10_-transformed output versus input variant frequencies. Each marker denotes average frequency across two replicate experiments. White and grey markers indicate variants with and without stop codons, respectively. Black dashed line indicates the 1:1 line; red dotted lines indicate 2-fold enrichment (>2x, above) or depletion (<0.5x, below) the 1:1 line. **e.** Amino acid enrichment among expression-depleted double mutants (output/input <0.5x). Letter heights reflect log_2_ enrichment in depleted fraction relative to all double mutants; tall letters indicate residues associated with reduced expression; color indicates chemical property. **f.** Number of variants present within two experimental replicates as a function of minimum concentration count threshold. Variants with fewer than 3 concentration points per replicate were excluded. **g.** Representative binding curves for three variants. Black solid line denotes jointly-fitted *K_d_*, while black dotted lines indicate *K_d_*± SD. Points represent normalized mean Cy5/GFP ratios (± SD), point colors denote replicate. **h.** Comparison of log_10_(*K_d_*) values from fluorescence polarization (± SD of technical replicates) and APB-TiteSeq (± joint fit SD). Marker colors match Fig. 3g. Black dotted lines and corresponding arrows denote upper and lower limits. Gray dashed line indicates 1:1 line; red dashed line indicates unweighted linear regression to points not indicated as limits. **i.** Scatter plot comparing log_10_(*K_d_*) for all single variants measured via APB-TiteSeq within the four position library versus the single-position library. Each marker represents fitted log_10_(*K_d_*); error bars indicate ±SD from the fit. Colored points indicate 7 ‘calibration’ peptides with associated FP measurements (colors match panel h). Gray dashed line indicates 1:1 line; red dashed line indicates unweighted linear regression to all points.

#### Comparing variant frequencies between input and GFP-positive beads quantifies expression levels

Producing soluble protein at sufficient levels remains a substantial challenge for protein functional characterization, and differences in expression level often confound attempts to quantify biophysical constants within cell-based assays [26, 75]. By comparing variant frequencies in sorted GFP-positive fractions versus the input library (Fig. 4c), APB-TiteSeq can directly quantify sequence-dependent differences in expression within the same experiment used to measure binding affinities.

We used this approach to quantify expression of all four-position library variants shared across two experimental replicates (20324/23283 variants, 87% coverage). Input and output frequencies were highly correlated between replicates (*R*^2^=0.92 and 0.85, respectively; Supplementary Figure 48), so we used replicate-averaged frequencies for all expression analyses. For WT FLAG and all single amino acid substitutions, output frequencies for the 153 variants without stop codons were within 2-fold of the input frequencies, while all 8 stop codon-containing variants were >2-fold depleted with a significantly shifted population mean relative to non-stop variants (permutation test *p* < 10^−4^) (Fig. 4d, Supplementary Figure 49). For the triple and quadruple mutants, which all lack stop codons, less than 0.2% were >2-fold depleted (Fig. 4d), suggesting relatively even expression levels for linear FLAG epitopes containing primarily polar or subtractive substitutions.

The double mutant library, generated via NNK codons, revealed a more complex expression landscape. As expected, >97% of the 764 double mutant variants containing premature stop codons were >2-fold depleted from the GFP-positive fraction (Fig. 4d) with a significantly shifted population mean relative to non-stop double mutant variants (permutation test *p* < 10^−4^; Supplementary Figure 49). However, 12% of non-stop double mutants were >2-fold depleted from GFP-positive fractions (Fig. 4d). This depletion could reflect sequence patterns that systematically reduce expression of full-length protein, though we cannot exclude increased sampling noise for low-abundance variants. To test whether sequence-dependent effects contribute to reduced expression, we compared the frequency of each amino acid in the >2-fold depleted double mutant pool to its frequency among all double mutants. Glutamine (Q), histidine (H), and proline (P) were significantly enriched within the >2-fold depleted pool (Fisher’s exact test, Bonferroni-corrected *p* < 0.05; Supplementary Figure 50) with position-specific enrichments that suggested distinct mechanisms (Fig. 4e). Prolines were preferentially depleted at terminal positions, possibly consistent with conformational strain in the nascent polypeptide chain driving premature termination. By contrast, Q and H were equally depleted across positions. As the codons for Q (CAA, CAG) differ from stop codons UAA and UAG by a single nucleotide at the first codon position, Q codons may be misread by release factors under cell-free expression conditions; the basis for H depletion is less clear. Regardless of mechanism, the amino acid- and position-specificity of these enrichments argues against sampling noise as the primary explanation for reduced expression and illustrates how APB-TiteSeq provides a simultaneous, integrated screen for expression efficiency alongside affinity measurements.

#### APB-TiteSeq affinities remain accurate for large libraries

Next, we quantified the accuracy of APB-TiteSeq affinity measurements as library size increases. We first established limits on resolvable affinities by simulating the APB-TiteSeq experimental pipeline for 20,000 variants with log-uniformly-distributed affinities between 0.5-50,000 nM using experimentally-derived parameters from each of two experimental replicates (Supplementary Figure 51 and Supplementary Tables 2 and 3). Returned affinities began to deviate from ‘true’ values (>25% outside the thermal noise threshold of 1 kT) for *K_d_*s tighter than 2.8 nM [2.6, 2.9] or weaker than 1.3 µM [1.2, 1.4] (mean [replicate 1, replicate 2]; Supplementary Figure 52), establishing upper and lower limits, respectively, for the 0.6-400 nM concentration range used here.

To estimate an apparent *K_d_* for each variant, we fit concentration-dependent binding curves for all variants from the four-position library that had >1 sorted bead in ≥ 3 concentrations in each of two experimental replicates (18,084 variants, 78% of the theoretical diversity; Fig. 4f). For each replicate, we globally fit mean Cy5/GFP ratios (Equation 1) at each antibody concentration to a Langmuir isotherm (Equation 2), again assuming a 1:1 binding stoichiometry and an experiment-specific saturation value (*R*_max_). Returned affinities agreed well between individual replicates (*R*^2^=0.84; Supplementary Figure 53). To aggregate information across replicate experiments and increase measurement precision, we re-calculated affinities using a joint fitting procedure that returned a single *K_d_* per variant and two experiment-specific *R*_max_ values. Returned aggregate affinities showed improved agreement across replicates (*R*^2^=0.97 and 0.92 against replicates 1 and 2 respectively; Supplementary Figure 53), and we used these jointly-fit affinities for all future analyses. Binding data were well-described by the Langmuir isotherm: the median NRMSE was 4.9% (10th percentile: 2.6%, 90th percentile: 19.6%; Supplementary Figures 54 to 58). For this four-position library, many variants (66%) bound the M2 antibody with fitted *K_d_*s at or above our simulation-estimated upper limit of detection (1.3 µM), as would be expected for heavily mutated epitopes (Supplementary Figure 59). A small subset (4%) bound with affinities within 2-fold of the WT epitope (3.9 nM). Given that our tight-binder limit (2.8 nM) is within 2-fold of the WT affinity, we did not see evidence of substantially higher-affinity variants within these 18,084 variants.

Log-transformed affinities from APB-TiteSeq still agreed strongly with FP measurements for calibration variants within our measurement limits (*R*^2^=0.97, RMSE=0.14; Fig. 4h). The only measurement with >2-fold deviations in *K_d_* was for the K3L variant, consistent with simulations that predicted this variant should be beyond our upper limit of detection (Supplementary Figure 60). Finally, APB-TiteSeq measurements of single-variant substitutions within the four-position library were highly correlated with measurements of the single-variant library alone (*R*^2^=0.98; Fig. 4i), further establishing reproducibility across libraries and experiments and that APB-Display can scale to return accurate affinities for >18,000 variants in parallel.

### LARGE-scale APB-TiteSeq datasets enable neural network denoising and detection of epistasis

To quantitatively understand how each position within the FLAG epitope contributes to M2 antibody binding affinity, we systematically assessed the degree to which models of increasing complexity could explain the observed binding affinity landscape, similar to previous approaches [76, 77]. At the simplest level, an additive PSAM model derived from single-mutant affinities explained a substantial fraction of variance in multi-mutant affinities (*R*^2^=0.84; Fig. 5a), confirming that independent positional contributions dominate the sequence-affinity landscape. To ask if non-additive interactions contribute beyond this additive baseline, we trained a convolutional neural network (CNN; Fig. 5b) on all measured affinities, allowing it to learn higher-order interactions from the data without imposing assumptions about their form. The CNN achieved strong predictive accuracy on held-out variants (average five-fold cross validation *R*^2^=0.94; Supplementary Table 4, Supplementary Figures 61 to 65). Across all variants, the RMSE between measured and CNN-predicted log-transformed *K_d_* values was 0.15 (within 1.4-fold or 0.3 kT; Fig. 5b), establishing that the model successfully captured the sequence-affinity landscape. This strong performance suggests the CNN can accurately predict affinities for mutant combinations not directly measured or with noisy measurements, providing a smoothed and complete map of the pairwise interaction landscape (Fig. 5c).

**Figure 5:**
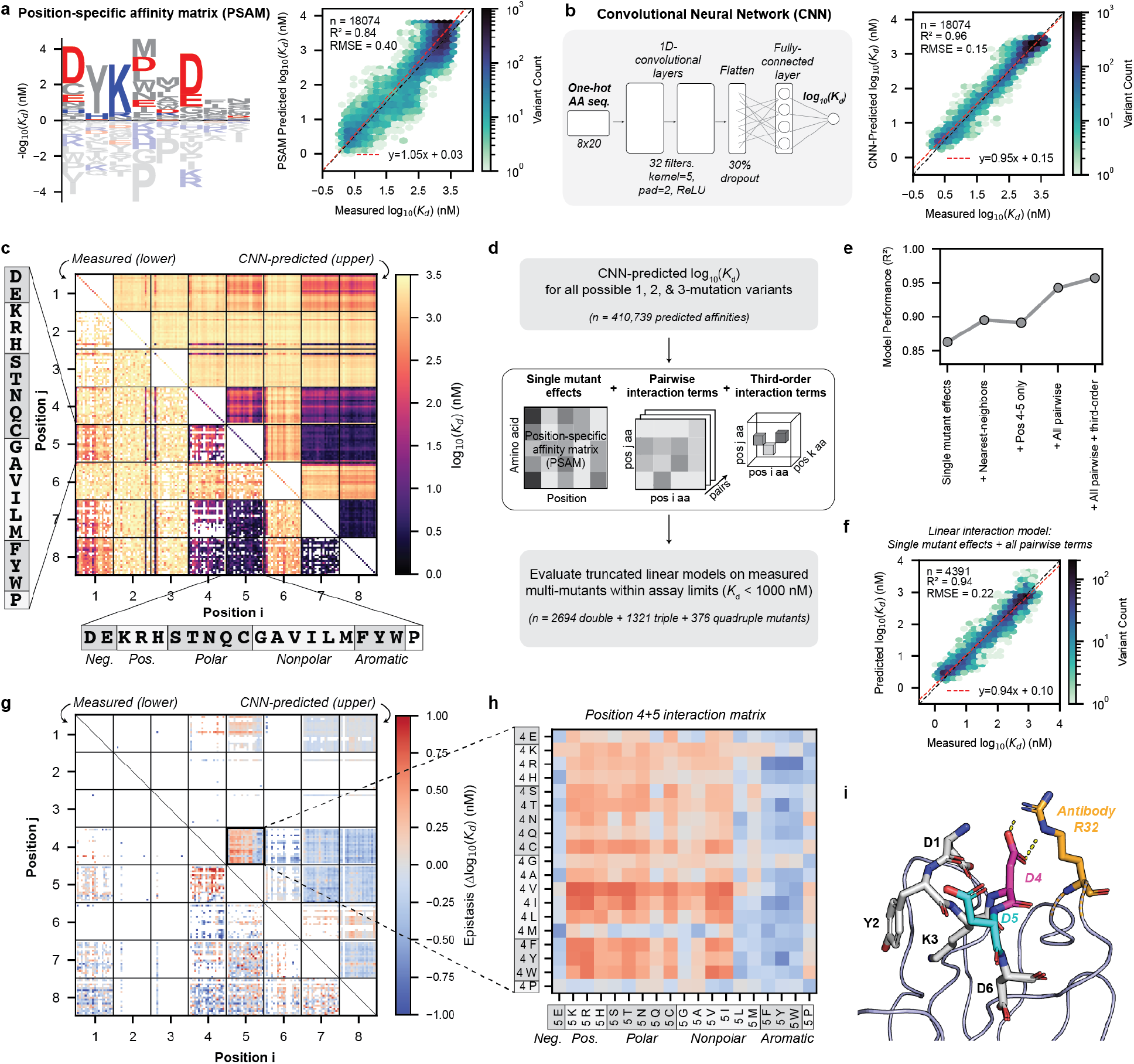
Machine learning analysis reveals epistasis in the FLAG/anti-FLAG landscape. **a.** Position-specific affinity matrix normalized to per-position mean log_10_(*K_d_*) (left) and single-site PSAM model predictions versus measured log_10_(*K_d_*) for the four-position library (right). PSAM predictions are bounded to the highest measured affinity (log_10_(*K_d_*)=3.8). **b.** CNN architecture schematic (left) and predictions versus measured log_10_(*K_d_*) (right). **c.** Measured (lower triangle) and CNN-predicted (upper triangle) affinities for double mutant variants. Diagonal: measured single-mutant affinities. Colors: log_10_(*K_d_*) in nM. **d.** Workflow for using CNN predictions to populate parameters of a linear interaction model. **e.** Performance (*R*^2^) of truncated linear models versus measured affinities (*K_d_*< 1000 nM). **f.** CNN-parameterized linear model predictions (all first- and second-order terms) versus measured affinities (*K_d_*< 1000 nM). **g.** Measured (lower triangle) and CNN-predicted (upper triangle) pairwise epistasis terms for double mutants (*K_d_*< 1000 nM). Colors: Δlog_10_ (*K_d_*) deviation from additive prediction. **h.** CNN-derived epistasis values for all pairwise substitutions at positions 4 and 5. **i.** Crystal structure of M2 Fab bound to FLAG (PDB: 8RMO). Antibody backbone: ribbon; R32: orange sticks; FLAG peptide: white sticks with D4 (magenta) and D5 (cyan); yellow dotted lines: D4/R32 salt bridge.

#### Linear interaction models reveal underlying structure of the sequence-affinity landscape

To decompose the CNN’s predictions into interpretable components, we used it as an oracle to populate the coefficients of a series of linear models incorporating an increasing number of between-residue interaction terms (Fig. 5d). Specifically, we performed *in silico* mutagenesis to derive coefficients quantifying single-mutant effects, pairwise interaction terms, and third-order interaction terms from CNN-predicted affinities. As mutant combinations with predicted affinities weaker than lower limits (∼ 1000 nM) could not be accurately measured, we excluded them from this analysis. We then compared the predictions of each model against experimental measurements (Supplementary Figure 66). Single-mutant effects alone explained substantial variance (*R*^2^=0.86; Fig. 5a,e), as expected for interactions between a folded antibody and a short linear motif (SLiM) like the FLAG epitope [18, 50, 62]. Incorporating nearest-neighbor coupling terms slightly improved model performance, with most of this gain due to interactions between positions 4 and 5 (*R*^2^=0.86 to *R*^2^=0.89; Supplementary Figure 66), in line with previous reports proposing non-additivity between these two sites [50]. Including all remaining pairwise interactions drove additional small improvements (Fig. 5f), possibly due to weak, longer-range pairwise interactions mediated through the peptide conformational ensemble [78]. Incorporating third-order interaction terms did not drive additional improvements, indicating that high-order epistasis is minimal in this system.

Since pairwise interactions dominate the non-additive structure of the landscape, we visualized all quantifiable pairwise interaction terms from measured and CNN-predicted datasets (Fig. 5g). These heatmaps reveal position-specific patterns of non-additivity: simultaneous mutations at positions 1+4, 1+5, 6+7, and 6+8 generally lead to positive epistasis (weaker-than-additive binding), while simultaneous mutations at positions 4+6, 4+7, 4+8, 5+7, and 5+8 generally lead to negative epistasis (tighter-than-additive binding). Simultaneous mutations at positions 4+5 lead to positive or negative epistasis depending on the chemical identity of the amino acid at position 5 (Fig. 5h).

#### Proposed mechanism for non-additivity between positions 4 and 5

The crystal structure of the bound FLAG/anti-FLAG complex provides a potential mechanistic explanation for this observed epistasis between positions D4 and D5. FLAG residue D4 forms a salt bridge with antibody residue R32 while neighboring D5 is solvent exposed (Fig. 5i). Thus, when D4 is mutated, the neighboring D5 residue may form an alternative salt bridge with R32 that rescues the interaction. Consistent with this model, position 5 mutations that facilitate this interaction (*e.g.* glutamate (E)) generate negative (tighter-than-additive) epistasis, while mutations that disrupt the interaction (*e.g.* positive, polar, and small nonpolar residues) generate positive (weaker-than-additive) epistasis (Fig. 5h). Large nonpolar and aromatic residues at position 5 are an exception to this rule, suggesting that Van der Waals interactions or aromatic cation-pi interactions may offer alternative modes of productive interaction with R32 (Fig. 5i).

### APB-SortSeq scales to libraries of >**100,000 variants via single-concentration affinity estimation**

FACS sorting time scales linearly with library size and number of ligand concentrations for APB-TiteSeq, creating a fundamental tradeoff between throughput and precision (Fig. 6a). While multi-concentration APB-TiteSeq can precisely quantify affinities for ∼20,000 variants, four-bin sorting at a single ligand concentration (“APB-SortSeq”, analogous to cell-based sort-seq methods [79]) offers a practical way to profile much larger libraries with reduced measurement precision.

**Figure 6:**
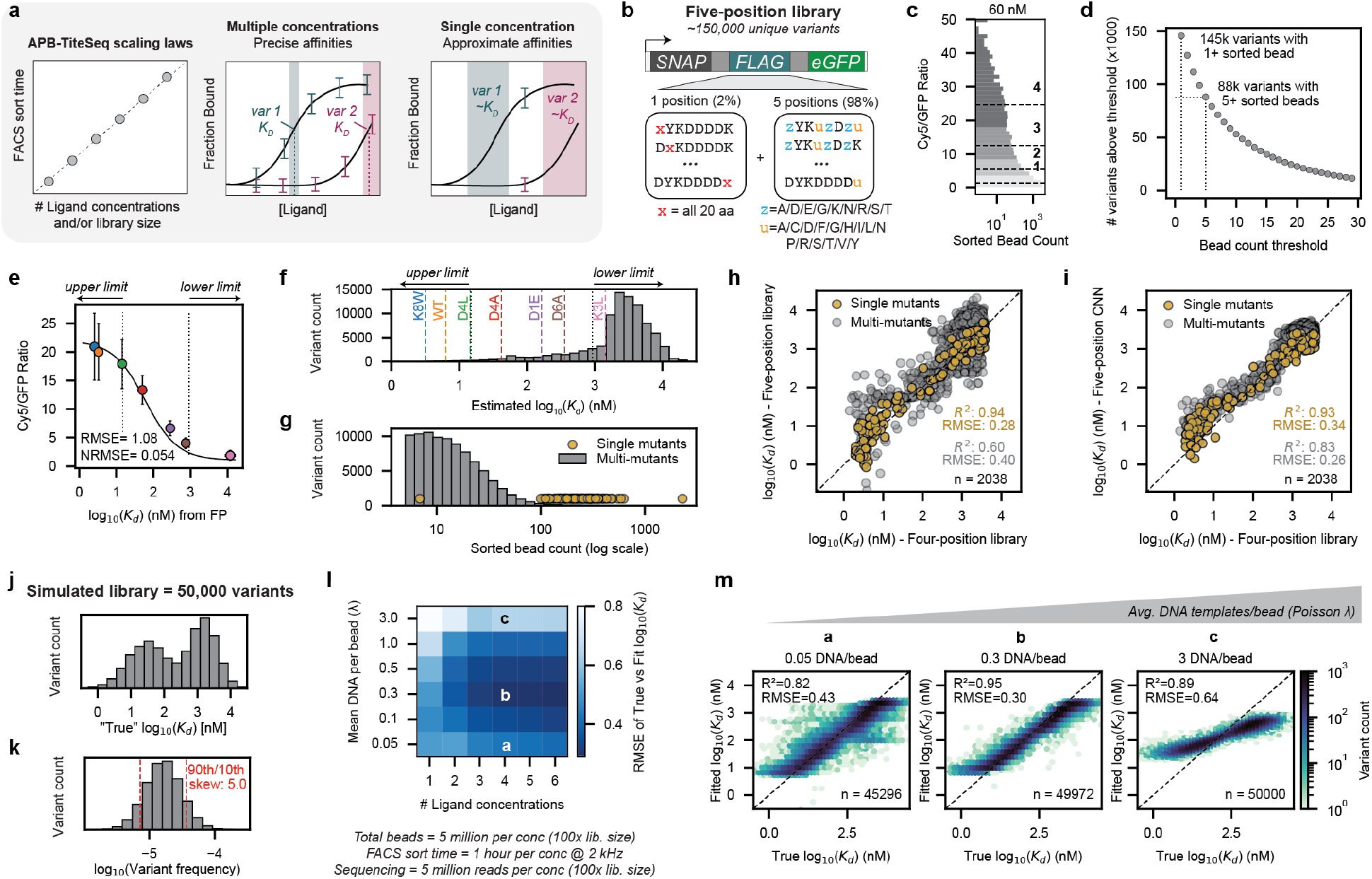
APB-SortSeq enables quantitative affinity estimation across library sizes via tunable experimental design. **a.** FACS sort time scales linearly with library size and number of ligand concentrations (left). Multiple concentrations enable precise *K_d_*measurements (center); a single concentration yields approximate estimates with higher throughput (right). **b.** Five-position library: 152 single-substitution variants (2%) plus combinatorial variants at positions 1, 4, 5, 7, and 8 (98%). **c.** Cy5/GFP distributions for beads sorted at 60 nM M2 antibody. Colors: sort bins; dashed lines: median bin values. **d.** Unique variants detected as a function of sorted bead count threshold. **e.** Calibration curve of mean Cy5/GFP ratios versus FP-measured log_10_(*K_d_*) (± SD). Black line: Langmuir isotherm fit determining *R*_max_ and *R*_min_; black dotted lines and arrows: simulation-defined upper and lower limit thresholds. **f.** Estimated log_10_(*K_d_*) distribution across the five-position library. Colored dashed lines: calibration variants; black dotted lines and arrows: simulation-defined upper and lower limit thresholds. **g.** Sorted bead counts for single-mutant (yellow) and multi-mutant (gray) variants (log scale). **h.** Measured log_10_(*K_d_*) for 2,038 variants shared between the four- and five-position libraries. Yellow: single-mutant variants; gray: multi-mutant variants. **i.** CNN-denoised log_10_(*K_d_*) values for five-position library variants versus four-position library measurements. Yellow: single-mutants; gray: multi-mutant variants. **j.** Simulated bimodal affinity distribution for a representative 50,000-variant library used to explore design parameters. **k.** Simulated library frequency distribution with 5-fold skew between 10th and 90th most abundant variants. Red dotted lines indicate log_10_ frequencies for the 10th and 90th most abundant variants. **l.** Measurement accuracy (RMSE of fitted vs true log_10_(*K_d_*) in nM) as a function of DNA loading rate (Poisson λ) and number of ligand concentrations. Letters (a,b,c) correspond to conditions in panel m. **m.** Hexbin plots of simulated fitted vs true log_10_(*K_d_*) at 4 log-spaced ligand concentrations (1, 10, 100, 1000 nM) across 3 DNA loading rates (0.05, 0.3, and 3 DNA/bead), illustrating tradeoffs between noise, dynamic range, and variant recovery.

To experimentally demonstrate this increased scale, we generated a 145,000-variant FLAG library consisting of two sub-libraries: a set of 152 single-substitution variants (2% of sequences) and a much larger combinatorial set of randomized codons at positions 1, 4, 5, 7, and 8 (up to 5 combinatorial mutations, 98% of sequences; Fig. 6b). Using cell-free Golden Gate assembly, we cloned this “five-position” library in 2 days; expressing and purifying all variants via APB-Display took an additional day and yielded 3% GFP-positive beads. We then incubated APBs overnight with a single antibody concentration (60 nM, selected to be near the midpoint of the previous concentration ranges) and sorted 1.6 million GFP-positive beads into four Cy5/GFP bins over 8 hours, equivalent to ∼10-fold library coverage (Fig. 6c, Supplementary Figures 67 and 68). Downstream sequencing returned over 145,000 and 88,000 variants with ≥1 and ≥5 sorted bead(s), respectively (Fig. 6d, Supplementary Figure 69).

#### Calibrated single-concentration measurements enable high-throughput affinity estimation

Mean Cy5/GFP fluorescence ratios (Equation 1) across the 88,083 variants with ≥ 5 sorted beads spanned a 22-fold range, from 1.1 to 24.1. To derive estimated affinities from these intensity ratios, we compared mean Cy5/GFP ratios to log_10_(*K_d_*) values for the 7 peptides previously characterized via FP (Fig. 6e). These ratios scaled log-linearly with *K_d_* and saturated at both extremes, consistent with Langmuir isotherm-type behavior and resolution limits predicted by experimentally-parameterized simulation (Fig. 6e, Supplementary Figures 70 and 71 and Supplementary Tables 2 and 3). Using the Langmuir binding equation (Equation 2) to calibrate Cy5/GFP ratios into affinities yielded estimated affinities between 0.01 nM and 32 μM; simulation established an assay dynamic range between 15 and 950 nM (Fig. 6f, Supplementary Figure 71). As expected for a library including extensive combinatorial substitutions, most variants bound weakly or not at all (81% of variants with *K_d_* >950 nM).

#### Ligand concentrations and bead sampling depth influence measurement precision

To understand how APB-SortSeq measurement precision depends on the number of concentrations sampled per variant, we compared fitted *K_d_* values for the 2,038 variants shared between the five-position library (measured at 1 concentration; Fig. 6c) and the four-position library (measured at 6 concentrations; Fig. 4b). We simultaneously investigated the impact of the number of beads sampled per variant by taking advantage of the fact that the 152 single-mutant variants were 10-fold more abundant than multi-mutant variants within the five-position library (100-1000 beads/variant; Fig. 6g). Affinities for the highly sampled single-mutant variants showed strong agreement across both libraries (*R*^2^=0.94, RMSE=0.28; Fig. 6h); the more sparsely sampled multi-mutant variants remained correlated but with substantially higher residuals (*R*^2^=0.60, RMSE=0.40; Fig. 6h). This degradation was most pronounced for variants with affinities far from the 60 nM staining concentration, where a single-point measurement provides limited discrimination. These results demonstrate that single-concentration sorting can approach the accuracy of multi-concentration experiments when variants with affinities near the staining concentration are sorted at sufficient depth.

#### CNN denoising recovers accurate affinities for sparsely sampled variants

To denoise the multi-mutant measurements by leveraging mutational patterns across sequence contexts, we trained a CNN on the five-position library. The average 5-fold cross-validation score for the five-position CNN (*R*^2^=0.70; Supplementary Figures 72 to 76 and Supplementary Table 5) was lower than for the four-position CNN (*R*^2^=0.94; Supplementary Table 4), consistent with higher per-variant noise from single-concentration measurements. Nevertheless, CNN-denoised affinities for shared multi-mutant variants agreed substantially better with the four-position library measurements never seen by the model (*R*^2^=0.83, RMSE=0.26; Fig. 6i), establishing that combining large, single-concentration datasets with simple CNN training can recover accurate affinities.

#### Parameter optimization for APB experiments

The increased noise observed for sparsely-sampled combinatorial variants raises a practical question: given a fixed sorting budget, what DNA loading rate (Poisson λ) and number of ligand concentrations maximize measurement quality? To address this, we used our APB-TiteSeq simulation pipeline to explore how these parameters impact measurement accuracies for a representative deep mutational scanning library of 50,000 variants with a bimodal affinity distribution (Fig. 6j) and 5-fold frequency skew between the 10th and 90th percentile variants (Fig. 6k), roughly based by the properties of our single-variant FLAG library. Measurement accuracy (RMSE of fitted versus true log_10_(*K_d_*)) improved with increasing numbers of log-spaced ligand concentrations before leveling off after 4 concentrations (Fig. 6l, Supplementary Figure 77). At a fixed number of concentrations, the mean DNA loading rate (Poisson λ) presented a clear tradeoff. Lower λ values favor clonality, yielding a wider dynamic range but increased per-variant noise and reduced recovery of rare variants (Fig. 6m). Higher λ values improve variant throughput [80] and reduce noise at the cost of reduced dynamic range. Mean DNA loading rates between 0.1 and 0.3 molecules per bead balanced these competing demands and enabled the highest measurement accuracy across all concentration sets.

## Discussion

Generating the large, standardized datasets needed to train predictive models linking protein sequence to function requires scalable, accessible, and precisely-controllable methods for quantifying functional parameters using thermodynamic constants. To meet these requirements, we developed three related platforms: APB-Display, APB-TiteSeq, and APB-SortSeq. Each builds on a shared set of core innovations: (1) dual-functionalized hydrogel beads for covalent DNA and protein capture, (2) particle-templated emulsification (PTE) that eliminates a need for microfluidics during expression and characterization, and (3) FACS-compatible bead sizes (∼ 18 μm) that enable rapid multi-bin sorting. APB-Display provides a foundation for quantifying expression and binding by co-localizing a protein variant with its encoding DNA on a single bead. APB-TiteSeq extends APB-Display by sorting beads across multiple ligand concentrations, returning absolute equilibrium dissociation constants for >10,000 variants in a single experiment. Where rough *K_d_* estimates are sufficient, APB-SortSeq uses a single ligand concentration combined with neural network denoising to extend throughput to >100,000 variants. Together, these platforms offer, to our knowledge, the first fully *in vitro* suite capable of measuring absolute *K_d_*s at this scale.

### Quantitative landscapes enable mechanistic insight and model training

We demonstrated all three APB platforms by quantifying interactions between the M2 antibody and libraries of FLAG epitope variants. The FLAG epitope is one of the most widely used commercial affinity tags [66], and several platforms have measured FLAG/anti-FLAG binding across single-variant (n=153 [49, 52]) and combinatorial-variant (n=12,739 [50]) libraries. However, none have reported absolute *K_d_* values. Using APB-TiteSeq and APB-SortSeq affinities to place these published datasets on an absolute *K_d_* scale reveals that array-based formats are limited by high background that results in small dynamic range (1–2 orders of magnitude), false positives, and poor predictions for *K_d_*s weaker than ∼100 nM (Supplementary Figures 78 and 79), likely due to nonspecific binding to array surfaces and large display proteins (*i.e.* polymerase, ribosome, dCas9). Each of these methods also predict tight affinity interactions that were not supported by either APB-TiteSeq or orthogonal FP measurements (DYKDDDDW; Supplementary Figures 79 and 80) or on-bead antibody titration (“superFLAG” DYKDEDLL; Supplementary Figures 81 and 82). Having established absolute *K_d_* values across >18,000 combinatorial variants spanning up to 4 simultaneous substitutions, APB-TiteSeq provides a reference landscape that exposes these limitations while extending dynamic range to ∼2.7 orders of magnitude (up to 1000 nM; Supplementary Figure 52). The scale and accuracy of these measurements enabled new mechanistic insights: we identified chemistry-dependent epistasis at positions 4 and 5, which we propose arises from salt bridge compensation between residue pairs. This demonstrates how standardized thermodynamic measurements at scale can not only detect epistasis but generate mechanistic hypotheses to explain it.

### Scalability and accessibility

All APB-TiteSeq and APB-SortSeq experiments reported here required approximately three days of hands-on time after DNA library preparation (1 day for APB generation, 1 day for FACS sorting, and 0.5 days for sequencing preparation), with total time and cost scaling with the number of variants and ligand concentrations (Supplementary Table 6). The 153-member single-variant library cost $13 per variant, while the ∼ 20,000 variant four-position library and the ∼ 100,000 variant five-position library cost $0.15 and $0.05 per variant, respectively. DNA synthesis costs were negligible for our large degenerate codon libraries (<$0.01 per variant) but represented a significant per-variant cost for our small user-defined libraries (∼ $5 per variant). For larger user-defined libraries, recent advances in oligo pool-based assembly methods [81, 82] offer promising ways to reduce synthesis costs while increasing diversity. Beyond time and cost, the only remaining barrier to adoption is the availability of dual-functionalized hydrogel beads; we are working to scale up bead production and distribution.

### Future directions and extensions

APB platforms are extensible along three axes: protein system, throughput, and dynamic range. Generalizing to new protein systems may require IVTT optimization. However, the successful folding of SNAP and eGFP tags within emulsion droplets, and prior IVTT production of diverse functional proteins including enzymes [83–85], transcription factors [11, 40, 41, 76, 86], and monobodies [42], suggest broad compatibility. Conveniently, the fluorescence-based expression readout on APBs is well-suited to guide expression optimization, as evidenced by our ability to identify residue-specific expression patterns across FLAG variants (Fig. 4c,d); this same readout can be used to screen IVTT conditions (*e.g.* temperature, additives, expression system) for other protein systems. To increase throughput, operating APB-SortSeq at high DNA loading rates (analogous to a recent bead-based screening approach [80]) could enrich very large libraries (> 10^6^ variants) before applying APB-TiteSeq for rigorous quantification (Fig. 6m). Finally, the dynamic range of APB-TiteSeq (∼ 1–1000 nM here) is adjustable: lower ligand concentrations will resolve tighter binders, while higher concentrations will extend measurements to weaker binders, as demonstrated previously with spectrally-encoded hydrogel beads [62, 63]. To facilitate these design decisions, our simulation pipeline identifies optimal experimental parameters (*e.g.* DNA loading rates, ligand concentrations, and bead counts).

Historically, the speed of computation has greatly outpaced the speed at which designed proteins can be experimentally characterized, and the lack of standardized, quantitative experimental datasets has hampered the development of accurate computational models [10]. APB-TiteSeq addresses both challenges: with a three-day turnaround time and per-variant costs below $0.15, it has the potential to enable active learning loops [13] that have previously been limited by experimental throughput and cycle iteration time. By returning standardized thermodynamic constants rather than relative rankings, each cycle of APB-TiteSeq produces data that is simultaneously useful for immediate design tasks and can be aggregated across molecules, laboratories, and iterations. We envision these cumulative datasets as the foundation for physically-grounded models that are accurate enough to reliably link protein sequence to function.

## Methods

### Plasmid construction

Gibson assembly (New England Biolabs) was used to generate a Golden Gate destination plasmid with an N-terminal SNAP-tag and C-terminal eGFP tag (SMT213). This plasmid also contains a chloramphenicol resistance gene, T7 promoter, E.coli translation initiation site, and hairpin terminator for compatibility with cell-free transcription and translation. FLAG variant DNA sequences with flanking BsaI recognition sequences were ordered as gBlocks or oligo pools (Integrated DNA Technologies) (Supplementary Table 7), then cloned into the SMT213 vector using Golden Gate assembly (New England Biolabs), transformed into NEB5-alpha competent cells (New England Biolabs), and recovered via miniprep (Qiagen). Whole-plasmid sequencing confirmed proper assembly of all variants.

To generate FLAG variant libraries with more than 10,000 variants, degenerate codon libraries with flanking BsaI recognition sequences were ordered as ssDNA oligos (Integrated DNA Technologies) (Supplementary Table 7), converted to dsDNA via Klenow extension (New England Biolabs), and assembled into the SMT213 construct via Golden Gate assembly (New England Biolabs). Assembled products were PCR amplified directly from the Golden Gate reaction using universal primers flanking the T7 promoter and terminator regions (Supplementary Table 8) to ensure compatibility with downstream assays (Primestar GXL, Takara Bio). Linear amplicons of the expected length (Supplementary Table 9) were gel-extracted and purified using the ZymoClean Gel DNA recovery kit (Zymo Research). Long-read sequencing (Plasmidsaurus) confirmed successful assembly of each amplicon library.

### Fluorescence microscopy

A Nikon Ti2 scope with SOLA light engine and Zymo cMOS camera was used to collect all microscopy images. Droplets were imaged using Countess cell counting slides, while beads were imaged within silicone gaskets (CultureWell, Grace Bio-Labs) adhered to glass slides. All images were collected with a 10x objective and 2×2 binning. Brightfield images were collected using overhead LED illumination passing through a Cy5 filter cube. Fluorescence images were collected using GFP and Cy5 filter cubes then darkfield- and flatfield-corrected with a custom python script. Circle finding (beads or droplets), median fluorescence intensity within each circle, and local background intensity around each circle were computed using the magnify image analysis package (https://github.com/FordyceLab/magnify) and custom python scripts.

### Chemical synthesis of benzylguanine methacrylate

Benzylguanine methacrylate (BG-MA) was synthesized as previously described [67]. Briefly, equal volumes of 40mM of BG-PEG-NH2 (New England Biolabs), 40mM methacrylic acid NHS ester (Sigma Aldrich), and 60mM triethylamine (Sigma Aldrich) were mixed together. All stock solutions were prepared fresh in DMSO (Sigma Aldrich). The reaction was incubated for 16 hours at 30°C with shaking at 400 rpm. The reaction was quenched with the addition of one volume of 100mM Tris-HCl (pH 7.5), followed by 3 hours of rotation at room temperature. The resulting 5mM BG-MA stock solution (concentration estimate based on 75 percent completion) was stored at -20°C for further use.

### Microfluidic device fabrication

The flow-focusing droplet generation device was slightly modified from [67] in AutoCAD2023 (Autodesk, Inc.) and ordered as a film photomask at 32,000 dpi (Fineline Imaging). Silicon master molds with 10 µm relief heights were constructed using SU-8-2010 photoresist and standard photolithography techniques. Poly(dimethylsiloxane) (PDMS) microfluidic devices were fabricated from the master molds using soft-lithography at a 1:5 elastomer base:crosslinker ratio (Momentive RTV 615). After a 15 min bake at 80°C, droplet generation devices were hole punched using a 1 mm biopsy punch (PicoPunch), monolithically bonded to a blank 1:10 PDMS slab, and baked at 80°C for 48 hours to bond the slabs together.

### Microfluidic generation of functionalized hydrogel beads

To generate hydrogel droplets functionalized with DNA oligos and benzylguanine, unpolymerized hydrogel mix was prepared with 10mM Tris-HCl pH 7.5 (Invitrogen), 15mM NaCl (Sigma Aldrich), 1mM EDTA (Invitrogen), 6% v/v mono:bis-acrylamide (37.5:1 ratio, BioRad), 0.3% ammonium persulfate (Sigma Aldrich), 30µM acrydite-modified oligonucleotide (IDT, Supplementary Table 8), and 50µM benzyl-guanine methacrylate (BG-MA). An oil mix was prepared with 0.4% N,N,N’,N’-Tetramethylethylenediamine (TEMED; Sigma Aldrich) in EvaGreen oil (BioRad). The unpolymerized hydrogel mix was filtered through a 0.22µm filter, then each of the hydrogel and oil mixes were loaded into syringes and connected to syringe pumps.

Hydrogel-in-oil droplets were formed by flowing the oil mix at 530 uL/hr and the unpolymerized hydrogel mix at 200uL/hr into the flow-focusing device. Given the flow rates and droplet diameter (∼14 µm = 1.43 pL Supplementary Figure 2), we estimate a ∼38 kHz droplet generation rate. Droplets were collected in 0.5-mL eppendorf tubes and incubated at room temperature to facilitate hydrogel polymerization. Polymerized hydrogel beads were recovered from the emulsions after 48 hours. After excess oil solution was drained from each tube by pipetting, 1H,1H,2H,2H-Perfluoro-1-octanol (PFO, Sigma Aldrich) was added at 50% of the volume of the emulsion pellet, and the tube was inverted 10 times to destabilize the emulsion. Tris-Tween buffer (10mM Tris HCl pH 7.5, 0.1% Tween-20) was added in 5-10x excess of the PFO-emulsion volume to extract the hydrogel beads into aqueous buffer solution. After vigorous tube inversion, a clear separation between aqueous and oil phases was observed, and the upper aqueous/hydrogel bead layer was transferred to fresh tubes. Further extractions with Tris-Tween buffer were performed until the translucent aqueous phase turned clear. Recovered hydrogel beads were washed 3x with Tris-Tween buffer by pelleting the beads via centrifugation (2000xg for 2 min), removing supernatant, and replacing with fresh buffer. Brightfield imaging using a calibration slide and image processing with custom python scripts were used to quantify the diameters of recovered hydrogel beads. Functionalized beads were stored at 4°C for further use.

### DNA template optimization for emulsion PCR

Digital droplet PCR was used to identify optimal DNA template concentrations for bead-based emulsion PCR. Based on molar concentration calculations for a given DNA template molecule, dilutions of a DNA stock solution (initial concentration verified by Qubit HS dsDNA assay, Invitrogen) were added to 25 µL PCR reactions containing PrimeSTAR GXL buffer (1x), PrimeStarGXL DNA polymerase (0.025 units/uL), dNTPs (200 µM), forward primer (0.2 µM), reverse primer (0.2 µM), and SYBR Green dsDNA dye (1x, Thermo Fisher). Assembled PCR reactions were emulsified with the addition of 60 µL EvaGreen oil (BioRad) and subsequent vortexing to generate polydisperse emulsion droplets with median diameters similar to that of the functionalized hydrogel beads (∼18 microns, typically 1 min of horizontal vortexing at 3000 rpm). The resulting emulsion was placed in a thermal cycler for 20 cycles of amplification (2 min 96° C, 20 cycles of [10 sec 96° C, 15 sec 60° C, 1 min/kb 68° C], 5 min 68° C, 12° C hold). After amplification, droplets were loaded into cell-counting slides (Countess, Invitrogen) for microscopy imaging in brightfield and GFP channels. The fraction of fluorescent droplets in each image were quantified using magnify (https://github.com/FordyceLab/magnify) and custom python scripts, then compared to Poisson statistics to estimate the average number of initial DNA molecules per droplet. Optimal concentrations were selected as those that closely aligned with the expected fraction of fluorescent droplets for the target Poisson distribution (typically 10% fluorescent droplets for a *λ* = 0.1 Poisson distribution).

### Emulsion PCR

A minimum volume of 20 uL of pelleted hydrogel beads in Tris-Tween buffer were used for each emulsion PCR reaction. PCR reagents were added to a final volume of 40uL (1x PrimeSTAR GXL buffer, PrimeStarGXL DNA polymerase (0.05 units/uL, Takara), 200µM dNTPs, 0.2µM forward primer, 0.002µM reverse primer) in a 0.2-mL tube. Plasmid DNA template was quantified using Qubit HS dsDNA assay kit and added to the reaction at the molar concentration required to achieve approximately 0.1 DNA molecules per 18-micron (3 pL) hydrogel bead. For larger libraries (*i.e.* the four and five position libraries), we linearly scaled up reaction volumes in 1.5-mL tubes.

Assembled PCR reactions were mixed by vortexing and incubated at room temperature for 3-5 min to allow PCR reagents to diffuse into the hydrogel beads. After incubation, beads were pelleted by centrifugation and 80% of the supernatant volume (typically 16uL for a 40uL PCR reaction) was removed by pipetting. The remaining bead pellet was vortexed briefly before 60uL of EvaGreen droplet oil (BioRad) was added to each reaction (2-3x the final reaction volume).

To generate particle-templated emulsions, sample tubes were vortexed for 5 minutes at maximum speed (3000 rpm) on a Fisher Scientific digital vortex mixer (120V, Cat No. 0215370). For best emulsion quality, tubes were placed horizontally on a foam tube holder (excised from Corning 480101) secured to the vortex mixer using lab tape, presumably to increase the force exerted on the hydrogel bead particles. The resulting emulsion was placed in a thermal cycler for 32 cycles of amplification (2 min 96° C, 32 cycles of [10 sec 96°C, 15 sec 60°C, 1 min/kb 68°C], 5 min 68°C, 12°C hold). For large libraries prepared in 1.5-mL tubes, we aliquoted emulsions into 0.2-mL tubes for thermal cycling.

After amplification, Amplicon Beads were recovered from the emulsions by draining excess oil and adding PFO at 50% of the volume of the emulsion pellet (typically 15-20uL). Tubes were inverted 10 times to promote emulsion destabilization. Next, 100uL of Tris-Tween buffer was added to extract the Amplicon Beads beads into aqueous buffer solution. After vigorous tube inversion, a clear separation between aqueous and oil phases was observed, and the upper aqueous/hydrogel bead layer was transferred to fresh 0.2mL PCR tubes. A second extraction with 100uL of Tris-Tween buffer was performed to increase bead recovery. Amplicon Beads were washed 3x with 150uL of Tris-Tween buffer, via cycles of buffer addition, vortexing, centrifugation, and supernatant removal. Amplicon Beads were stored at 4°C until further use. For large libraries, we transferred emulsions back to 1.5-mL tubes before adding PFO and linearly scaled volumes for Amplicon Bead extractions.

### Emulsion IVTT

Emulsion IVTT reactions (typically 30uL) were prepared by adding IVTT reagents to the Amplicon Bead pellets recovered from the emulsion PCR step. PURExpress reagents (12uL Part A, 9uL Part B, New England Biolabs) were allowed to defrost on ice, then combined together for 15 minutes on ice before adding 0.8 Units/uL of SUPERase-In RNase inhibitor (Invitrogen). The IVTT mixture was added directly onto Amplicon Bead pellets in 0.2-mL tubes and mixed by vortexing. The sample was incubated on ice for 5 minutes to allow the IVTT reagents to soak into the hydrogel beads. For larger libraries (*i.e.* the four and five position libraries), we linearly scaled up reaction volumes in 1.5-mL tubes.

Next, 60uL of EvaGreen oil (BioRad) was added to each tube, and particle-templated emulsions were generated by vortexing horizontally at 3000 rpm for 3-5 minutes. Emulsion IVTT samples were incubated at 37°C for 2 hours in a thermal cycler or 1.5-mL heat block to promote protein expression and SNAP-tag mediated bead capture. After incubation, Amplicon/Protein Beads were recovered from emulsions exactly as described in the emulsion PCR section, with the exception that PBS-T buffer (1X PBS, 0.1% Tween-20) was used for the bead extraction and wash steps. Amplicon/Protein Beads were stored at 4°C, protected from light, until further use.

### Protein concentration calibration

A fluorescence standard curve for estimating protein concentration on beads was generated using vortexed emulsion droplets containing known concentrations of recombinant eGFP protein (Origene). 40 µL of dSurf oil was added to 15 µL of serially-diluted eGFP protein in PBS-T buffer, then the samples were vortexed horizontally for 1 min to form polydisperse emulsion droplets. Emulsions were loaded into cell-counting slides (Countess, Invitrogen) for microscopy imaging in brightfield and GFP channels (300ms exposure times). Median GFP intensities for droplets with 16-24 µm diameters were quantified for each dilution sample using magnify (https://github.com/FordyceLab/magnify) and custom python scripts. Linear regression of eGFP concentration versus median fluorescence intensity in bead-sized droplets produced a standard curve for estimating protein concentrations from microscopy images of droplet-loaded Amplicon/Protein Beads in the GFP fluorescence channel (collecting using identical 300ms exposure times).

### FLAG/anti-FLAG binding assay

To measure binding of M2 anti-FLAG antibody (Sigma F1804) to variants of the FLAG epitope, Amplicon/Protein Beads displaying SNAP-FLAG-eGFP variants were blocked for 30 minutes in PBS-T + 1% BSA buffer, then mixed with varying concentrations of the M2 antibody in PBS-T + 1% BSA buffer. An equimolar amount of Cy5-labeled secondary antibody (goat anti-mouse IgG, Thermo Fisher A10524) was pre-complexed with the primary antibody for 30 minutes on a rotator at room temperature prior to bead addition. Beads and antibodies were incubated for at least 18 hours on a rotator at 4°C to allow the binding interactions to reach equilibrium.

To verify that protein attachment and antibody staining were specific, nonspecific binding controls were performed using beads added directly to IVTT reactions rather than Amplicon/Protein Beads prepared by emulsion PCR and emulsion IVTT. Three conditions were tested: (1) BG-functionalized beads added to an IVTT reaction expressing SNAP-FLAG-eGFP, (2) BG-functionalized beads added to an IVTT reaction expressing SNAP-eGFP (no FLAG tag), and (3) non-functionalized beads (lacking BG) added to an IVTT reaction expressing SNAP-FLAG-eGFP. All reactions were incubated for 2 hours at 37°C on a rotator to ensure uniform protein attachment across beads. After protein expression, beads were washed, blocked, and stained with pre-complexed M2 and Cy5 secondary antibody as described above, then imaged by fluorescence microscopy.

To identify equilibrium conditions, BG-functionalized beads displaying SNAP-FLAG-eGFP (prepared as above) were imaged at 0.5, 15, and 39 hour timepoints after mixing with the antibodies, and the Cy5/GFP intensity ratio was fit to an exponential binding approach curve to determine *t*_99_.

### Bead sorting with FACS

Immediately prior to bead sorting, beads were pelleted and washed 3x in 400uL of PBS-T buffer and filtered with a 35 µm strainer into 5-mL FACS tubes. Beads were sorted on a BD FACS Aria II sorter in the Stanford Shared FACS Facility equipped with a 100 µm nozzle. Drop delay calibration was performed with 6 µm Accudrop beads (BD) and fine-tuned with 20 µm AccuCount beads (Spherotech). Upon flowing beads labeled with eGFP-tagged FLAG variants and Cy5-labeled antibodies, singlet beads were identified using forward and side scatter parameters, while GFP-positive beads were identified relative to the large population of blank beads in each APB sample. For single-bead sorting, GFP-positive beads were sorted into individual wells of an empty 384-well plate using the “index sort” feature to record the fluorescence profiles of each sorted bead. For pooled sorting, GFP-positive beads were sorted into four empty 1.5-mL tubes in “4-way purity” mode according to roughly log-spaced Cy5/GFP ratiometric gates, adjusted to match the dynamic range of the sample while maintaining roughly equal populations in each bin.

### Amplicon sequencing from sorted single beads

DNA amplicons were recovered from Amplicon/Protein Beads deposited into a 384-well plate via multi-stage PCR. First, sorted beads were concentrated at the bottom of the 384 well plate by centrifugation. Next, PCR reagents were added to each well to a final volume of 8 µL (1x PrimeSTAR GXL buffer, PrimeStarGXL DNA polymerase (0.025 units/µL), 200µM dNTPs, 0.2 µM inner forward primer, 0.2 µM inner reverse primer). To extract only the FLAG variant region, universal inner primers recognizing the conserved regions flanking the variant region (Supplementary Table 8) were used for the first 10 rounds of amplification (1 min 96°C, 10 cycles of [10 sec 96°C, 15 sec 60°C, 15 sec 68°C], 1 min 68°C, 12°C hold). To append well-specific barcodes, 2 µL of 384 uniquely-barcoded outer primers (0.2 µM each) from the evSeq pipeline (https://github.com/fhalab/evSeq) were stamped onto the plate prior to 15 more rounds of amplification (1 min 96° C, 15 cycles of [10 sec 96° C, 15 sec 60° C, 20 sec 68° C], 1 min 68° C, 12° C hold). Plate-based PCR samples were pooled and gel-purified as described in the evSeq protocol with minor adjustments. Briefly, 5 µL of PCR supernatant from each well were pooled together in the presence of 4mM EDTA (pH 8.1). Amplicons of target length (262 bp; Supplementary Table 9) were extracted from a 1% agarose gel and purified with the ZymoClean Gel DNA recovery kit (Zymo Research D4001). Purified amplicon DNA was quantified using a Qubit dsDNA HS assay kit (Invitrogen) and submitted for custom long-read sequencing by Plasmidsaurus (Eugene, OR). Sequencing reads were filtered and demultiplexed based on exact matches to FLAG variant and barcode sequences using custom python scripts with Dask-based parallelization.

### Bootstrap analysis of down-sampled single-bead populations

To assess the minimum number of APBs required for reliable variant characterization, we performed bootstrap analysis on Cy5/GFP ratio measurements from individual sorted beads. For each variant, beads were sampled with replacement from the full population to generate 1,000 bootstrap replicates across subpopulation sizes of 5, 10, 15, 20, 25, and 30 beads. For each replicate, the median Cy5/GFP ratio was computed, and the coefficient of variation (CV%; sample standard deviation divided by mean, expressed as a percentage) was calculated across the 1,000 replicate medians to quantify measurement precision as a function of sample size. Only beads with at least 25 sequencing reads per variant were included. Bootstrap sampling was performed on the full bead population (including multi-variant beads), using a fixed random seed for reproducibility.

### Amplicon sequencing from four-way sorted beads

DNA amplicons were recovered from sorted Amplicon/Protein Beads in 1.5-mL eppendorf tubes via multi-stage PCR. Briefly, sorted beads were concentrated at the bottom of the sort tube by brief centrifugation, then 200uL of Tris-Tween buffer was added to dilute the FACS sheath buffer. Bead solutions were transferred to 0.2-mL PCR tubes and pelleted by centrifugation. The supernatent was removed to leave approximately 10 µL of bead pellet and buffer in each tube. PCR reagents were added to a final volume of 16uL (PrimeSTAR GXL buffer (1x), PrimeStarGXL DNA polymerase (0.025 units/uL), 200 µM dNTPs (200 µM), inner forward primer (0.2 µM), and inner reverse primer (0.2 µM)). To extract only the FLAG variant region, universal inner primers recognizing the conserved regions flanking the variant region (Supplementary Table 8) were used for the first 10 rounds of amplification (1 min 96°C, 10 cycles of [10 sec 96°C, 15 sec 60°C, 15 sec 68°C], 1 min 68°C, 12°C hold). To append sort bin-specific barcodes, 4 µL of uniquely-barcoded outer primer set (0.2 µM each) from the evSeq pipeline (https://github.com/fhalab/evSeq) was added to each PCR reaction prior to 15 more rounds of amplification (1 min 96°C, 15 cycles of [10 sec 96°C, 15 sec 60°C, 20 sec 68°C], 1 min 68°C, 12°C hold).

Barcoded amplicons from each sort bin were purified using KAPA purification beads (Roche) using a 0.9X bead-to-sample volumetric ratio to retain fragments ≥250 bp. Purified amplicons were eluted into nuclease free water and quantified using a Qubit dsDNA HS assay kit (Invitrogen). Barcoded amplicons from each sort bin were combined in mass ratios proportional to the number of beads in each sort bin and submitted as a single pool for custom long-read sequencing by Plasmidsaurus (Eugene, OR). Barcoded sequencing reads were demultiplexed by filtering for exact matches to 4 bp regions flanking the FLAG variant region (total length = 32, minimum Phred 20 quality score at each base (99% accuracy)), followed by filtering for exact matches to evSeq barcodes (minimum 1 barcode match, minimum Phred 20 quality score at each base) using custom python scripts with Dask-based parallelization.

### Library frequency and expression analysis

Sequencing reads from input (original) and output (post-sort) libraries were filtered to retain only variants matching expected sequence patterns (e.g., single variants or mutations only at positions 1, 4, 5, and 7). Variants were classified by edit distance (Hamming distance) from the wild-type FLAG sequence (DYKDDDDK). Variant frequency was calculated as the read count of a given variant divided by total library read count. For output libraries, all sorted reads were analyzed as a single pool without bead count normalization across samples. For the four-position library, per-variant frequencies were averaged across two biological replicates (DRP236 and DRP237) and variants with fewer than 10 reads were excluded from analysis. Replicate concordance was assessed by Pearson linear regression of log-transformed frequencies, with *R*^2^ computed as the square of the Pearson correlation coefficient. The log_2_ ratio of output to input frequency was used as the measure of relative expression; variants with log_2_(output/input frequency) < −1 (i.e., >2-fold depleted) were classified as depleted.

To compare the log_2_ frequency ratio distributions of stop codon–containing versus non-stop variants, we used a two-sided permutation test (10,000 permutations). In each permutation, the pooled log_2_ frequency ratios from both groups were randomly reassigned to groups of the same sizes as the observed groups, and the difference in group means was computed. The *p*-value was the fraction of permutations in which the absolute difference in means was at least as large as the observed value; when no permutation exceeded the observed statistic, *p* was reported as < 10−4.

To identify amino acids enriched among depleted double mutants, we extracted only the mutated positions from each double mutant sequence (positions differing from WT) and tallied the amino acid composition of the depleted pool (log_2_ ratio < −1, stop codons excluded) and the background pool (all double mutants, stop codons excluded). For each of the 20 canonical amino acids, a two-sided Fisher’s exact test was performed on the 2 ×2 contingency table of (amino acid present / absent) ×(depleted pool / background pool). *p*-values were Bonferroni-corrected across the 20 tests, and amino acids with corrected *p* < 0.05 were considered significantly enriched in the depleted pool. To visualize position-specific enrichment patterns, we computed per-position log_2_ odds ratios between the position frequency matrix (PFM) of the depleted pool and the PFM of the background pool (pseudocount 0.001 added to each frequency), retaining only positive (enriched) values.

### FACS analysis for determining representative Cy5/GFP values for each sort bin

FACS data were analyzed using an identical gating scheme to that applied during sorting. The median Cy5/GFP ratio within each gate served as the representative bin value. To account for nonspecific antibody binding at higher concentrations, background Cy5/GFP ratios were estimated via bootstrap sampling (10,000 iterations): each iteration sampled one Cy5 value from the blank bead population and one GFP value from the GFP-positive bead population (both with replacement), and the background ratio was taken as the median of the resulting distribution. Background-subtracted median Cy5/GFP ratios were used to represent the fluorescence intensity of a given bin for all mean Cy5/GFP calculations; background-subtracted 10th and 90th fluorescence values were used to estimate the fluorescence spread for uncertainty propagation.

### Estimating binding affinities from binned sequencing reads

Sequencing reads were filtered to retain only variants that matched expected sequence patterns (e.g., single variants or mutations at expected positions). Variants with premature stop codons were excluded from affinity analysis. Read counts for each variant–sort bin combination were normalized by the total number of sorted beads in each bin such that each normalized read count represents a single sorted bead. Depending on the library, variants with low read counts (e.g. <5 normalized reads for the five-position library) or insufficient read counts across concentrations (e.g. <3 concentrations with at least 1 normalized read for the four-position library) were filtered out.

For each variant *v* at antibody concentration *c*, we estimated the mean Cy5/GFP ratio *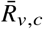* using a mean-bin inference approach adapted from [71]. Briefly, background-subtracted Cy5/GFP fluorescence ratios were measured for cells sorted into four bins. For each bin *b*, the median Cy5/GFP ratio *m_b,c_* was determined from the sorted population (Supplementary Table 3). Sequencing read counts were normalized to estimated bead counts per variant, and a pseudocount of 0.1 was added to each bin to stabilize estimates for low-abundance variants. The fractional bin probability for variant *v* in bin *b* was then computed as

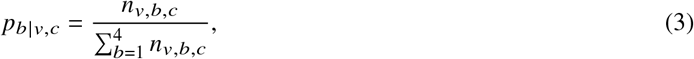

where *n_v,b,c_* is the pseudocount-adjusted normalized read count for variant *v* in bin *b* at concentration *c*, and the mean Cy5/GFP ratio was computed as the read-count-weighted average of bin medians:

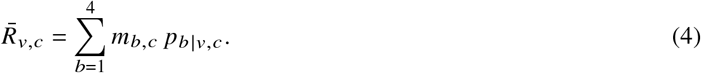

Uncertainty in 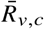 was propagated following Equation 6 in [71], adapted to linear fluorescence space. The within-bin fluorescence spread σ*_b,c_* was estimated from the 10th and 90th percentile Cy5/GFP values within each bin (*Q*1*_b,c_* and *Q*9*_b,c_*, respectively) using the normal-distribution quantile relation:

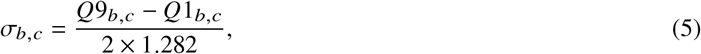

where 1.282 is the standard normal quantile at the 90th percentile. Error propagation through the weighted mean then yields

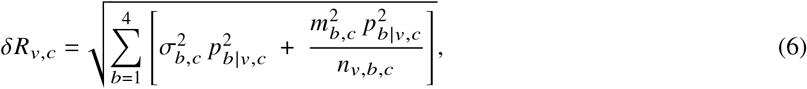

where the first term reflects within-bin fluorescence spread weighted by the squared bin probability, and the second term reflects Poisson sequencing noise, with higher-fluorescence bins contributing proportionally more uncertainty. Both terms are weighted by 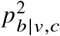 so the overall uncertainty is dominated by the bins in which the variant’s beads predominantly sorted.

For the single-variant library, a single global *R*_max_ was estimated by simultaneously fitting all variants’ concentration– response data to a Langmuir isotherm assuming 1:1 stoichiometry, 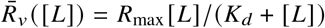 , using nonlinear least-squares optimization. Individual apparent *K_d_* values were then estimated for each variant by holding *R*_max_ fixed at this global value, and fit quality was assessed by RMSE and NRMSE (defined as RMSE / *R*_max_).

For the four-position library, apparent *K_d_* values were estimated by jointly fitting data from two replicate experiments simultaneously. Concentration–response data from each experiment were fit to the same Langmuir equation, sharing a single *K_d_* parameter per variant while allowing experiment-specific *R*_max_ values (one global *R*_max_ per experiment, shared across all variants); *K_d_* and the two *R*_max_ values were estimated by alternating least-squares, iterating between a per-variant *K_d_* update (nonlinear least squares minimizing pooled residuals across both experiments) and closed-form *R*_max_ updates until convergence.

For the five-position library, variants were measured at a single antibody concentration, precluding full isotherm fitting. Instead, the binding isotherm equation was inverted to extract *K_d_* from each variant’s mean Cy5/GFP ratio. Specifically, *R*_max_ was estimated by fitting 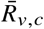 for the seven calibration variants as a function of their fluorescence polarization-derived *K_d_* values to the Langmuir equation (scipy `curve_fit`, bounds [0, 100]); *R*_min_ was set to the background-subtracted median Cy5/GFP ratio for the lowest sort bin. Mean Cy5/GFP ratios were then normalized to fractional occupancy as 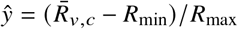, and *K_d_* was computed analytically as *K_d_* = [*L*](1 − ŷ)/ŷ, where [*L*] is the antibody concentration. Variants whose normalized ratio fell outside [0, 1] were excluded from further analysis.

### Simulation of APB-TiteSeq workflow

To characterize the accuracy and dynamic range of APB-TiteSeq as a function of experimental parameters, we implemented a Monte Carlo simulation recapitulating each physical step of the workflow. Briefly, a virtual library of *N* variants with assigned true *K_d_* values is loaded onto beads under Poisson statistics, amplified by emulsion PCR, and expressed by IVTT; stochastic noise is introduced at each stage via log-normal multiplicative factors modeling variability in PCR amplification, protein expression, and antibody labeling. Sources of cross-contamination (*e.g.* including rare DNA or protein transfer between beads during emulsion breaking and co-encapsulation of two beads in the same emulsion IVTT droplet) are also modeled. Beads are then virtually sorted by Cy5/GFP ratio across a titration series of ligand concentrations, and sequencing reads are drawn by multinomial sampling from the sorted amplicon pools. The resulting count data passes through the same analysis pipeline as experimental data to yield per-variant mean Cy5/GFP ratios, from which *K_d_* values are estimated by fitting a non-cooperative 1:1 binding isotherm.

To define the measurable affinity range, we compared fitted *K_d_* values to ground truth using a rolling-percentile analysis. Simulated variants were sorted by their true *K_d_*, and a sliding window (0.1 log_10_-unit wide) was advanced along the true-*K_d_* axis. Within each window, we computed the 25th, 50th, and 75th percentiles of the fitted log_1_0 (*K_d_*)values. We defined a thermal noise threshold of 0.44 log_10_ units, corresponding to 1 kT (≈ 0.6 kcal/mol ΔΔ*G*) at room temperature. The *weak-affinity bound* is the fitted *K_d_* at which the local 25th percentile of fitted values first exceeds the true *K_d_* by this threshold — *i.e.*, the point at which 25% of variants in a local window have their affinities underestimated by more than 1 kT. The *strong-affinity bound* is defined symmetrically as the fitted *K_d_* at which the local 75th percentile falls below the true *K_d_* by the same threshold, indicating that 25% of variants in that window are overestimated by more than 1 kT. Both bounds are expressed as fitted rather than true *K_d_* values so they can be applied directly to experimental measurements. The difference between the two bounds in log_10_ units defines the dynamic range of the assay. A full description of all simulation stages and parameters are provided in Supplementary Note 1 and Supplementary Table 2, respectively.

### Fluorescence polarization assay

N-terminal FITC-Ahx labeled peptides (95% purity, TFA removal, Genscript) diluted to 0.3 nM in PBS-T buffer were mixed with increasing concentrations of M2 anti-FLAG antibody (Sigma F1804) in a multi-well plate. Fluorescence polarization was measured using a Tecan Spark plate reader. Polarization values were normalized to no-antibody controls. Three technical replicates at each concentration were globally fit to a Langmuir isotherm to extract equilibrium dissociation constants (*K_d_* ± SD) using a custom python script.

### Position-specific affinity matrix (PSAM) model

To evaluate the degree to which multi-mutant affinities can be predicted from single-mutant effects alone, we constructed a position-specific affinity matrix (PSAM) from measured single-mutant *K_d_* values in the four-position library. For each position-amino acid combination, the single-mutant effect was defined as the difference between the jointly-fitted log_10_(*K_d_*) for that variant (*i*) and wildtype FLAG (*WT*). Multi-mutant affinities were predicted additively as:

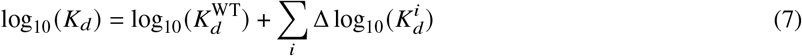

where the sum is taken over all mutated positions *i* and 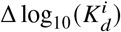 is the single-mutant effect at position *i*. Predicted values exceeding log_10_(*K_d_*) = 3.8 (the highest recorded affinity in the four-position library) were bounded to this value. PSAM predictions were evaluated against all jointly-fitted *K_d_* values in the four-position library, including lower-limit estimates, since correctly predicting that a variant binds weakly is meaningful even when neither the measured nor predicted *K_d_* can be precisely quantified.

### Neural network model for predicting FLAG binding

We trained a convolutional neural network (CNN) model to predict binding affinities (log_10_(*K_d_*)) from one-hot encoded FLAG epitope sequences. The CNN is composed of two 1D convolutional layers with kernel size of 5 and padding of 2, both with a filter size of 32, followed by a fully connected layer with dropout *p* = 0.3. To fully capture the context of the 8-residue peptide, we used a kernel size of 5 across stacked convolutional layers to achieve a cumulative receptive field of 9. This architecture ensures that the theoretical receptive field encompasses the entire peptide sequence, allowing every position within the input to influence the final prediction.

We trained a CNN on the four position library of 18,074 variants. We separately trained a CNN on the five-position library of 87,452 variants, with points with negative log_10_(*K_d_*) filtered out. Training was conducted for 100 epochs, with learning rate *lr* = 0.001 and the Adam optimizer. Both the four position and five position datasets were randomly split into five folds for training and validation, and the single-position mutants and wild type sequences were added separately to the training set to ensure their presence during model fitting. For each library, the five CNNs - each with a separate fold held-out - were ensembled together for inference on unseen data points.

### Epistasis analysis via CNN-derived linear interaction model

To decompose the four-position sequence-affinity landscape into interpretable additive and epistatic components, we used the four-position CNN (trained on all experimental data) as an oracle to derive the coefficients of a linear interaction model via *in silico* mutagenesis. For each single, double, triple, and quadruple mutant in the four-position sequence space, we queried the CNN directly to obtain predicted log_10_(*K_d_*) values. Single-mutant effects were defined as the difference between the CNN-predicted log_10_(*K_d_*) for each single mutant and wildtype. Pairwise coupling terms were then defined as the residual after subtracting wildtype (*WT*) and both single-mutant effects (*i, j*) from the CNN-predicted double-mutant affinity (*ij*):

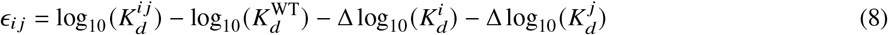

Third-order interaction terms were derived analogously by subtracting wildtype, all three single-mutant effects, and all three pairwise coupling terms from the CNN-predicted triple-mutant affinity. To evaluate the contribution of each interaction order, we constructed a series of truncated linear models of increasing complexity — single-mutant effects only, single-mutant effects plus nearest-neighbor pairwise terms (sequentially adjacent positions), single-mutant effects plus all pairwise terms, and single-mutant effects plus all pairwise and third-order terms — and evaluated each against experimental measurements. CNN-derived coefficients were used directly without refitting to experimental data. Variants with CNN-predicted log_10_(*K_d_*) > 3.0 were excluded from model evaluation to avoid artifacts resulting from lower-limit measurements; CNN-measured values were used to define this threshold to avoid circularity in the evaluation.

### Comparison to published FLAG binding datasets

Median Cy5/GFP ratios from the 25-variant FLAG library were compared to three published FLAG variant binding datasets: Prot-MaP [50], PICASSO [52], and CasPlay [49]. For each comparison, variants shared with the APB library were identified by exact amino acid sequence match, and median per-bead Cy5/GFP ratios were compared directly to the corresponding published metric. For PICASSO and CasPlay, linear regression was performed between APB median Cy5/GFP ratio and each assay’s binding score across shared variants. For Prot-MaP, which reports fluorescence intensities at six antibody concentrations (0, 1, 3, 10, 32, and 100 nM), linear regression was performed at each concentration separately; variants with APB median Cy5/GFP ratio ≤ 10 were excluded from regression because they show no measurable response in the Prot-MaP dataset at any reported concentration.

To characterize each assay’s dynamic range in affinity space, (*K_d_*)values from three larger libraries (single-variant and four-position APB-TiteSeq and five-position APB-SortSeq) were compared against published measurements for shared variants. For PICASSO and CasPlay, the detectable affinity interval was defined as the contiguous APB *K_d_* range over which the absolute rolling Spearman correlation between APB log_10_(*K_d_*) and binding score was≥ 0.5, computed in a centered rolling window of width max (10, 0.15· *N*) variants, where *N* is the number of shared variants. The reported dynamic range (in orders of magnitude) was calculated as the width of this interval on a log_10_(*K_d_*) scale, averaged across the three library comparisons. For Prot-MaP, rather than comparing to concentration-specific fluorescence intensities (as above), we used the reported Limit of Detection (LoD), a single summary metric derived across the full antibody concentration series, enabling direct comparison with APB-TiteSeq *K_d_* values. Variants annotated as “LoD >max” were assigned LoD = 100 nM. The upper *K_d_* boundary of Prot-MaP sensitivity was defined as the lowest APB-TiteSeq *K_d_* at which ≥ 50% of variants within a centered rolling window of width max 20, 0.05 ·*N* variants had LoD = 100 nM, indicating loss of affinity resolution above that value. The reported dynamic range was calculated as the difference between this upper boundary and the minimum APB-TiteSeq *K_d_* observed in the overlap population, on a log_10_(*K_d_*) scale, averaged across the three library comparisons.

### On-bead antibody titration of superFLAG and wildtype FLAG

SNAP-DYKDEDLL-eGFP (“superFLAG”), SNAP-DYKDDDDK-eGFP (“wildtype FLAG”), and SNAP-eGFP (negative control) fusion proteins were immobilized on BG-coated beads and incubated with M2 anti-FLAG primary antibody at seven concentrations (0.7, 2, 6, 13, 20, 60, and 100 nM) followed by a Cy5-conjugated secondary antibody for 21 hours at 4°C with rotation to ensure equilibrium binding. Beads were diluted 10-fold in buffer prior to fluorescence imaging. Per-bead median Cy5 and GFP intensities were background-subtracted using the local background median, and the background-subtracted Cy5/GFP ratio was computed for each bead; beads with non-positive background-subtracted GFP intensity were excluded. A Langmuir isotherm was fit to the per-bead Cy5/GFP ratios using alternating least squares, estimating a shared global *R*_max_ across the wildtype and superFLAG variants with an individual *K_d_* per variant. Per-concentration median Cy5/GFP values were used for fitting.

## Acknowledgments

D.R.P. was supported by a Stanford Graduate Fellowship and the Molecular Biophysics Training Program at Stanford (T32 GM136568). A.G. is supported by the Stanford Knight-Hennessey fellowship. S.T. is supported by the Arnold and Mabel Beckman Foundation through the Arnold O. Beckman Postdoctoral Fellowship in Chemical Sciences. This work was funded by NIH DP1CA290563, a Stanford Bio-X Interdisciplinary Initiatives Seed Grant (IIP11-68), and a Schmidt Sciences Polymath award to P.M.F. P.M.F. is also a Chan Zuckerberg Biohub San Francisco Investigator.

D.R.P. is grateful for support from Fordyce lab members past and present (particularly the Particle Party subgroup), thoughtful comments on the manuscript from Shawn Costello, Albert Lee, Matt DeJong, and Renee Hastings, and helpful suggestions regarding simulations and statistical analysis from Fletcher Passow. D.R.P. also thanks Iain Clark for particle-templated emulsification training and the Stanford Shared FACS Facility for FACS training and instrument access. Single-bead data was collected on an instrument in the Shared FACS Facility obtained using NIH S10 Shared Instrument Grants S10RR027431-01 and S10RR025518-01.

## Author Contributions

D.R.P., S.T., and P.M.F. conceptualized the project. D.R.P. performed all experiments, simulations, and related data analysis. A.G. trained the neural networks and performed related data analysis. D.R.P. and P.M.F. wrote the manuscript. S.T., A.K., and P.M.F. supervised the project.

## Conflict of Interest

Stanford University has filed a patent (US Patent application no. PCT/US63/707264) on APB-Display, and D.R.P and P.M.F. are named inventors. P.M.F. is a co-founder of Velocity Bio. The remaining authors declare no competing interests.

## Data Availability

All Autocad files, images, CSV data files, and data analysis notebooks will be made available at the OSF repository associated with this paper.

## Notes

### Summary of Updates

First introduction paragraph revised for clarity; author acknowledgements updated.

