## Supplementary Information for "Amplicon/Protein Bead Display enables quantitative *in vitro* biochemistry at scale"

Daria R. Passow[1], Anvita Gupta[2], Samuel Thompson[3], Anshul Kundaje[2,4], and Polly M. Fordyce[3,4,5,6,*]

[1]Biophysics Program, Stanford University
[2]Department of Computer Science, Stanford University
[3]Department of Bioengineering, Stanford University
[4]Department of Genetics, Stanford University
[5]ChEM-H Institute, Stanford University
[6]Chan Zuckerberg Biohub, San Francisco, CA

July 17, 2026

### List of Supplementary Figures


1 Microfluidic droplet generator. 4
2 Images and diameters of hydrogel droplets before and after polymerization into beads. 5
3 Poisson-predicted probabilities of a given number of DNA templates per droplet as a function of $\lambda$. 6
4 DNA template dilutions in vortexed emulsion droplets to identify optimal DNA concentrations for APB-Display. 7
5 Binding equilibration timecourse for SNAP-FLAG-eGFP and SNAP-eGFP constructs. 8
6 Microscopy-based quantification of GFP and Cy5 intensities for the 25-variant FLAG library. 8
7 Nonspecific binding controls for bead attachment and antibody staining. 9
8 Representative images of vortexed emulsions loaded with increasing concentrations of eGFP protein. 10
9 Calibration curve for converting GFP intensities into concentrations. 11
10 Distribution of estimated per-bead GFP concentration and molecule counts for the 25-variant library. 11
11 Calculations to test if reaction conditions lead to a ligand depletion regime. 12
12 FACS gating scheme for identifying singlet GFP-positive Amplicon/Protein beads for the 25-variant library. 13
13 Number of sorted single beads as a function of read count threshold for the 25 variant library. 14
14 Distribution of the number of sorted beads per variant for the 25 variant library. 14
15 Distribution of total read counts for beads containing variants with at least 25 reads for the 25 variant library. 15
16 Correlation between GFP intensity and total read counts on single beads for the 25 variant library. 15
17 Measured Cy5 versus GFP intensities for all beads with at least 25 reads corresponding to a given variant for the 25 variant library. 16
18 Measured Cy5 versus GFP intensities for all beads with at least 25 reads corresponding to only a single variant for the 25 variant library. 17
19 Comparison of Cy5/GFP ratio distributions for beads displaying only a single variant and all beads. 18
20 Bootstrap-sampled median Cy5/GFP distributions for 10-bead APB populations for the 25-variant library. 19
21 Bootstrap-sampled median Cy5/GFP distributions for 20-bead APB populations for the 25-variant library. 20
22 Median Cy5/GFP CV% as a function of sampled bead count for the 25-variant library. 21
23 Comparison of APB Cy5/GFP ratios from the 25-variant library with Cas9-display binding scores. 22
24 Comparison of APB Cy5/GFP ratios from the 25-variant library with Prot-MaP fluorescence intensities. 23
25 Log-log FACS plots for the single-variant library at each antibody concentration. 24
26 Linear-linear FACS plots for the single-variant library at each antibody concentration. 24
27 Correlation between read counts and sorted bead counts for each concentration-bin sample in the single-variant library. 25
28 APB-TiteSeq simulation pipeline to determine likely upper and lower limits for resolvable affinities for APB-TiteSeq experiments. 25
29 Simulated affinity and frequency distributions used to determine likely upper and lower limits for resolvable affinities for single-variant library experiments. 26
30 Comparison between fitted and true affinities from simulations of the single-variant library. 26
31 Binding isotherms for single-variant library (page 1/8). 27
32 Binding isotherms for single-variant library (page 2/8). 28
33 Binding isotherms for single-variant library (page 3/8). 29
34 Binding isotherms for single-variant library (page 4/8). 30
35 Binding isotherms for single-variant library (page 5/8). 31
36 Binding isotherms for single-variant library (page 6/8). 32
37 Binding isotherms for single-variant library (page 7/8). 33
38 Binding isotherms for single-variant library (page 8/8). 34
39 Fit quality metrics for single-variant library. 35
40 Distributions of fitted $\log_{10}(K_d)$ values for the single-variant library. 35
41 Comparison of FP affinities and simulated affinity distribution for the single-variant library. 36
42 Log-log FACS plots for the four-position library at each antibody concentration (replicate 1). 37
43 Linear-linear FACS plots for the four-position library at each antibody concentration (replicate 1). 38
44 Log-log FACS plots for the four-position library at each antibody concentration (replicate 2). 39

45 Linear-linear FACS plots for the four-position library at each antibody concentration (replicate 2). . . . 40
46 Correlation between read counts and sorted bead counts for the four-position library (replicate 1). . . . . 41
47 Correlation between read counts and sorted bead counts for the four-position library (replicate 2). . . . . 42
48 Comparison of input and output library frequencies across replicates of the four-position library. . . . . 43
49 Histograms of $\log_2$(output/input) frequency ratios as a function of edit distance for the four-position library. 44
50 Amino acid enrichment in depleted double mutant FLAG variants from the four-position library. . . . . 45
51 Histograms of affinity and frequency distributions for simulating the four-position library. . . . . . . . . 46
52 Simulation method for determining dynamic range limits for the four-position library. . . . . . . . . . . 46
53 Comparison between individual replicate fits and joint fits for the four-position library. . . . . . . . . . 47
54 Fit quality metrics for four-position library. . . . . . . . . . . . . . . . . . . . . . . . . . . . . . 48
55 Binding isotherms for randomly-sampled variants with affinities <10 nM in the four-position library. . . 49
56 Binding isotherms for randomly-sampled variants with affinities between 10 nM and 100 nM in the four-position library. . . . . . . . . . . . . . . . . . . . . . . . . . . . . . . . . . . . . . . . 50
57 Binding isotherms for randomly-sampled variants with affinities between 100 nM and 1000 nM in the four-position library. . . . . . . . . . . . . . . . . . . . . . . . . . . . . . . . . . . . . . . . 51
58 Binding isotherms for randomly-sampled variants with affinities >1000 nM in the four-position library. . 52
59 Distribution of returned fitted $\log_{10}(K_d)$ values for the four-position library. . . . . . . . . . . . . . . . 53
60 Comparison of FP-measured affinities and simulated affinity distribution for the four-position library. . . 54
61 Performance of four-position library CNN during Fold 1 of 5-fold cross-validation. . . . . . . . . . . . 54
62 Performance of four-position library CNN during Fold 2 of 5-fold cross-validation. . . . . . . . . . . . 55
63 Performance of four-position library CNN during Fold 3 of 5-fold cross-validation. . . . . . . . . . . . 55
64 Performance of four-position library CNN during Fold 4 of 5-fold cross-validation. . . . . . . . . . . . 56
65 Performance of four-position library CNN during Fold 5 of 5-fold cross-validation. . . . . . . . . . . . 56
66 Correlation between predicted and measured $\log_{10}(K_d)$ from linear models of increasing complexity derived from CNN predictions. . . . . . . . . . . . . . . . . . . . . . . . . . . . . . . . . 57
67 Log-log FACS plot for five-position library at a single antibody concentration. . . . . . . . . . . . . . . 58
68 Linear-linear FACS plot for five-position library at a single antibody concentration. . . . . . . . . . . . 58
69 Correlation between read counts and sorted bead counts for the five-position library. . . . . . . . . . . . 59
70 Histograms of affinity and frequency distributions for simulating the five-position library. . . . . . . . . 59
71 APB-SortSeq simulation pipeline for determining upper and lower limits of resolution for the five-position library. . . . . . . . . . . . . . . . . . . . . . . . . . . . . . . . . . . . . . . . . . . . 60
72 Performance of five-position library CNN during Fold 1 of 5-fold cross-validation. . . . . . . . . . . . 60
73 Performance of five-position library CNN during Fold 2 of 5-fold cross-validation. . . . . . . . . . . . 61
74 Performance of five-position library CNN during Fold 3 of 5-fold cross-validation. . . . . . . . . . . . 61
75 Performance of five-position library CNN during Fold 4 of 5-fold cross-validation. . . . . . . . . . . . 62
76 Performance of five-position library CNN during Fold 5 of 5-fold cross-validation. . . . . . . . . . . . 62
77 Correlation between returned fitted and "true" input $\log_{10}(K_d)$ values from APB-TiteSeq binding simulations as a function of the number of ligand concentrations and mean DNA/bead parameters for a representative library. . . . . . . . . . . . . . . . . . . . . . . . . . . . . . . . . . . . . . 63
78 Comparison between Prot-MaP Limit of Detection (LoD) values APB-TiteSeq-measured $K_d$ values. . . 64
79 Comparison of APB-TiteSeq-measured $K_d$ values with PICASSO Cas9-Display binding scores. . . . . 64
80 Comparison of APB-TiteSeq-measured $K_d$ values with CasPlay Cas9-Display binding scores. . . . . . . 65
81 Microscopy images of concentration-dependent antibody binding for "superFLAG" and WT FLAG variants. 66
82 Microscopy-generated binding curves for "superFLAG", WT FLAG, and no FLAG variants. . . . . . . 67

### List of Supplementary Tables


1 Sequencing run summary for APB experiments. . . . . 68
2 Simulation input parameters for each experimental dataset. . . . . 69
3 Per-concentration simulation parameters for each experimental dataset. . . . . 70
4 Four-position CNN cross-validation results. . . . . 71
5 Five-position CNN cross-validation results. . . . . 71
6 Cost analysis for APB experiments. . . . . 72
7 FLAG library sequences used in APB experiments. . . . . 73
8 Primers used in APB experiments. . . . . 74
9 Amplicon sequences used in APB experiments. . . . . 74

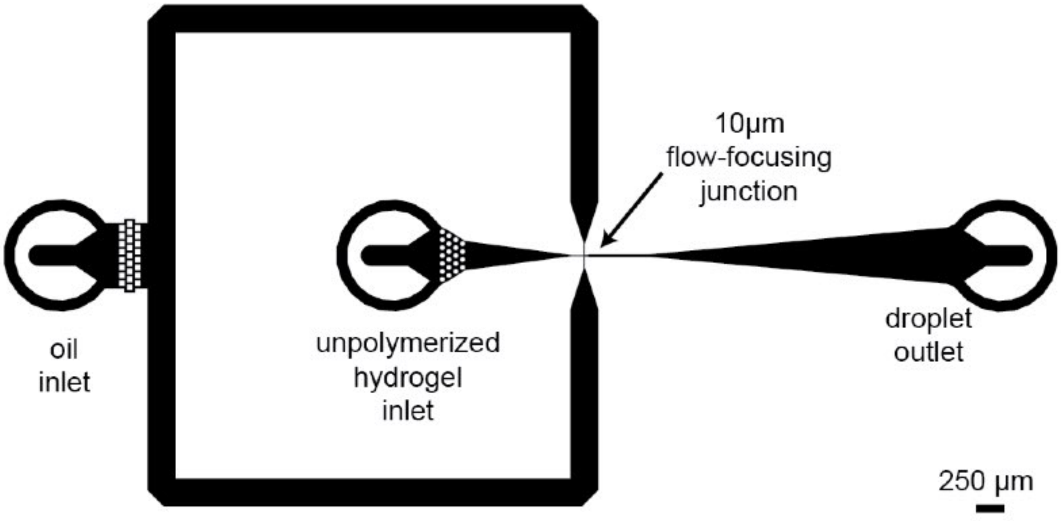


**Supplementary Figure 1 — Microfluidic droplet generator.**
Schematic of CAD file showing the droplet generator used to make Amplicon/Protein Beads.

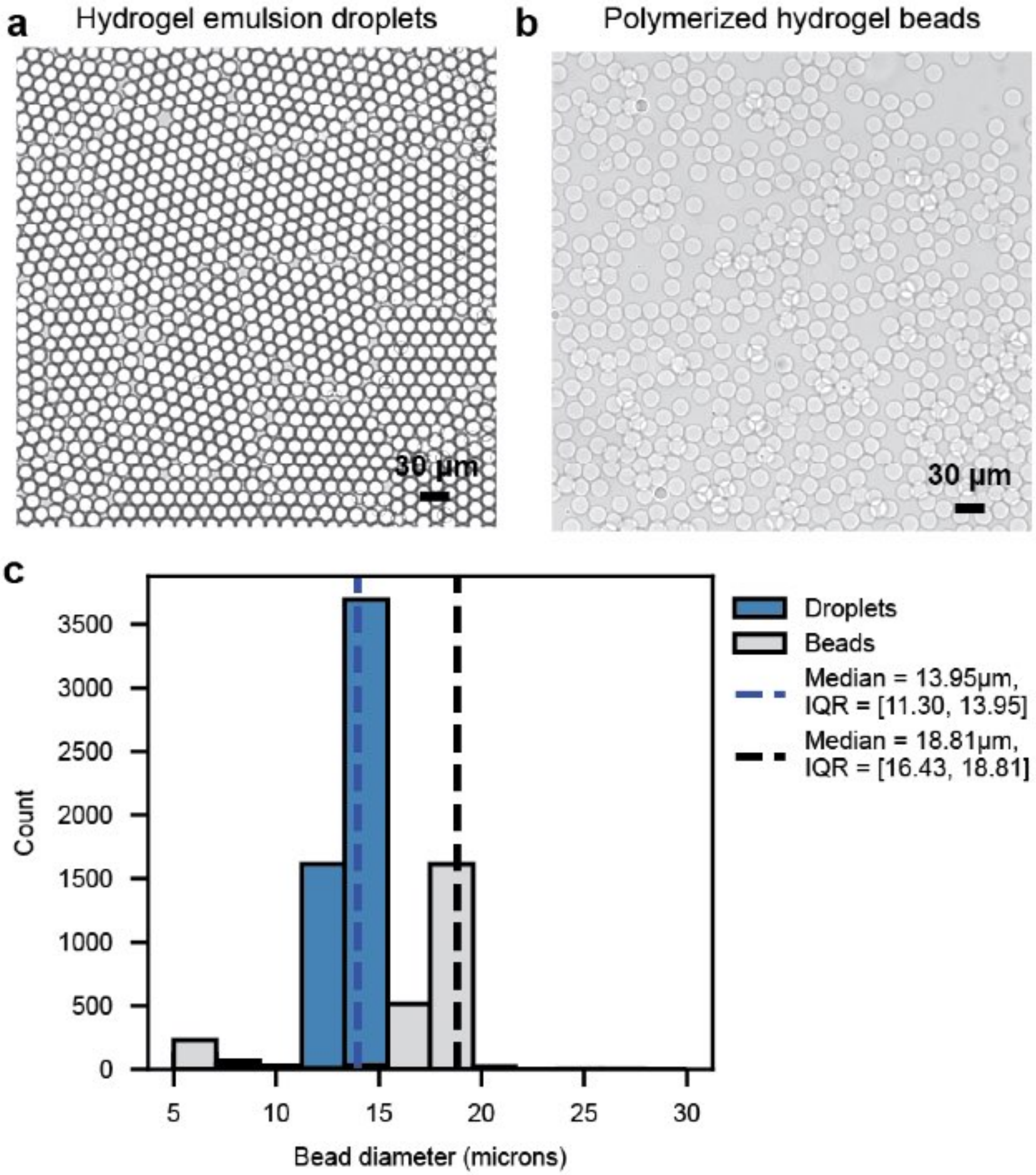


**Supplementary Figure 2 — Images and diameters of hydrogel droplets before and after polymerization into beads.**

(a) Bright field image of microfluidically-generated hydrogel droplets. Scale bar = 30 µm. (b) Bright field image of polymerized hydrogel beads. (c) Histogram of droplet and bead diameters, with annotated median and interquartile range [0.25, 0.75] in µm.

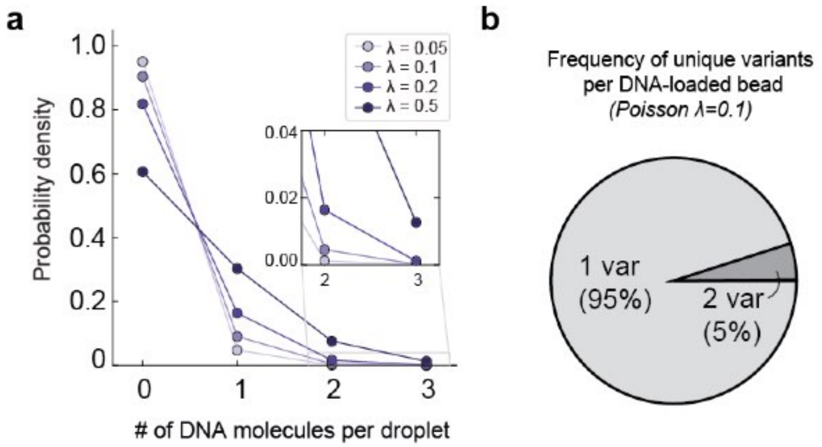


**Supplementary Figure 3 — Poisson-predicted probabilities of a given number of DNA templates per droplet as a function of $\lambda$.**

(a) Scatter plot showing the Poisson-predicted probability density of a given number of DNA molecules per droplet as a function of Poisson $\lambda$ values. (b) Pie chart showing the fraction of DNA-loaded droplets containing 1 or 2 template sequences at $\lambda$=0.1.

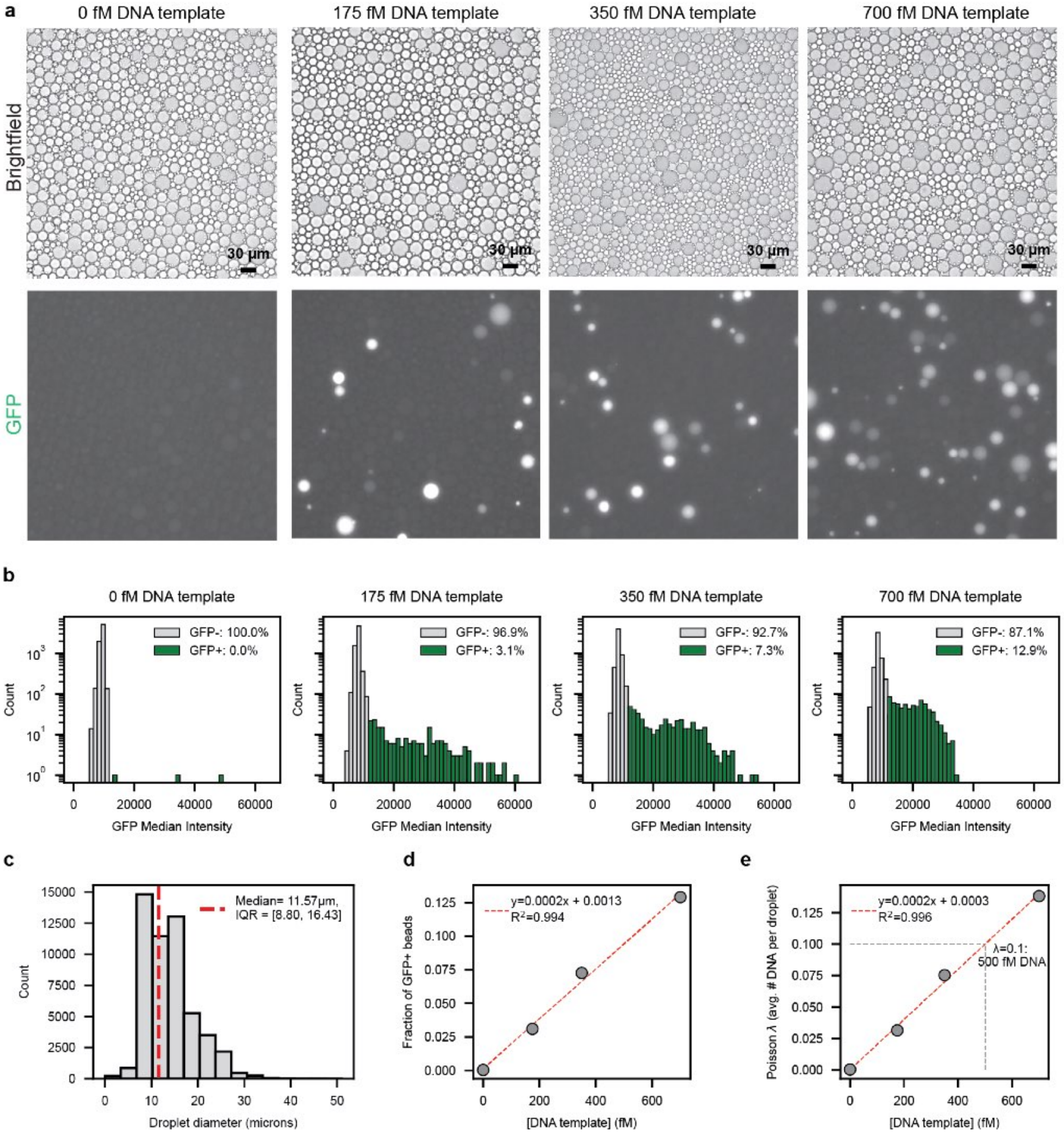


**Supplementary Figure 4 — DNA template dilutions in vortexed emulsion droplets to identify optimal DNA concentrations for APB-Display.**

(a) Representative bright field (top row) and fluorescence (bottom row) images from digital droplet PCR assay used to determine template concentrations near $\lambda$=0.1 for emulsion PCR. Droplets are loaded wth SybrGreen DNA intercalating dye such that DNA amplification can be detected in the GFP channel. Scale bar = 30 µm. (b) Histograms of median GFP intensities for each DNA loading condition. Gray bars: non-fluorescent droplets; green bars: fluorescent droplets. (c) Histogram of droplet diameters across images used for template dilution analysis. (d) Scatter plot of the fraction of GFP-positive beads as a function of the input DNA concentration. Markers indicate estimated GFP-positive fractions from panel b; red dashed line indicates linear regression fit. (e) Scatter plot of Poisson $\lambda$ (estimated number of DNA templates per droplet) as a function of the input DNA concentration. Markers indicate calculated $\lambda$ values from estimated GFP-positive fractions from panel b; red dashed line indicates linear regression fit. Gray dotted lines identify the concentration of DNA template predicted to yield $\lambda$=0.1.

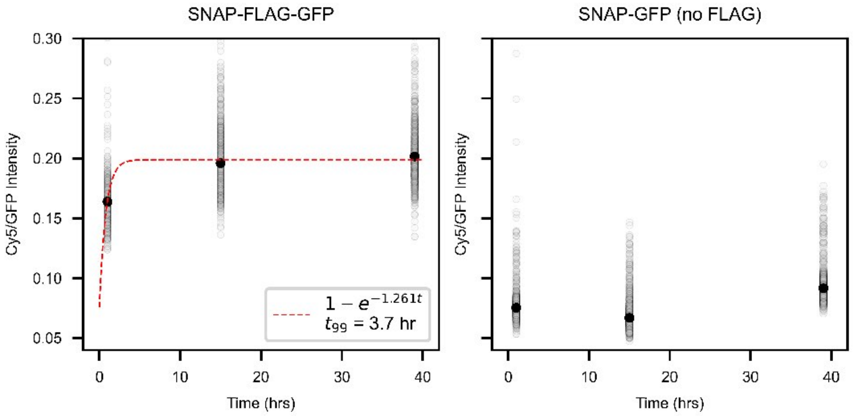


**Supplementary Figure 5 — Binding equilibration timecourse for SNAP-FLAG-eGFP and SNAP-eGFP constructs.**

Per-bead Cy5/GFP ratios as a function of incubation time with 6 nM M2 anti-FLAG and 6 nM Cy5-labeled secondary antibody. Black dots: median values. Red dotted line: exponential fit line to extract $t_{99}$, the time required to reach 99% of equilibrium saturation. Overnight incubation (>3.7 hours) appeared sufficient to reach equilibrium for SNAP-FLAG-eGFP constructs (left) without promoting non-specific interactions with constructs that lack the FLAG peptide (SNAP-eGFP, right).

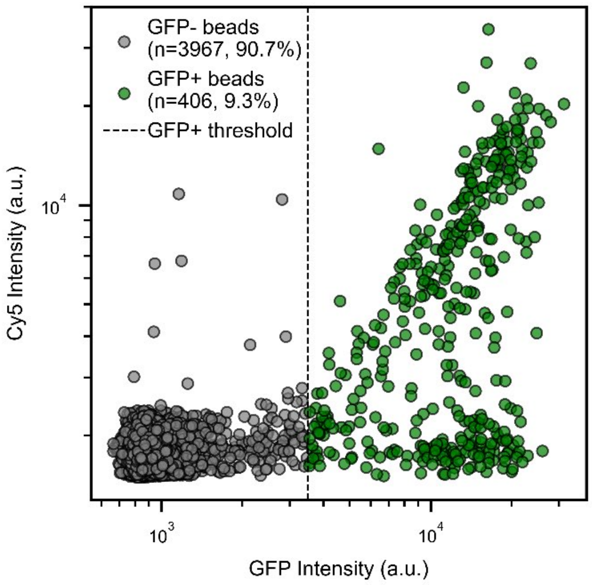


**Supplementary Figure 6 — Microscopy-based quantification of GFP and Cy5 intensities for the 25-variant FLAG library.**

Log-log scatter plot of per-bead Cy5 and GFP intensities from microscopy images of the 25-variant FLAG library (Cy5 exposure: 300ms; GFP exposure: 300ms). Black dotted line indicates the GFP-positive bead threshold, defined as the GFP intensity at which the Cy5 intensities begin rising linearly above background. Gray markers represent GFP-negative beads; green markers represent GFP-positive beads.

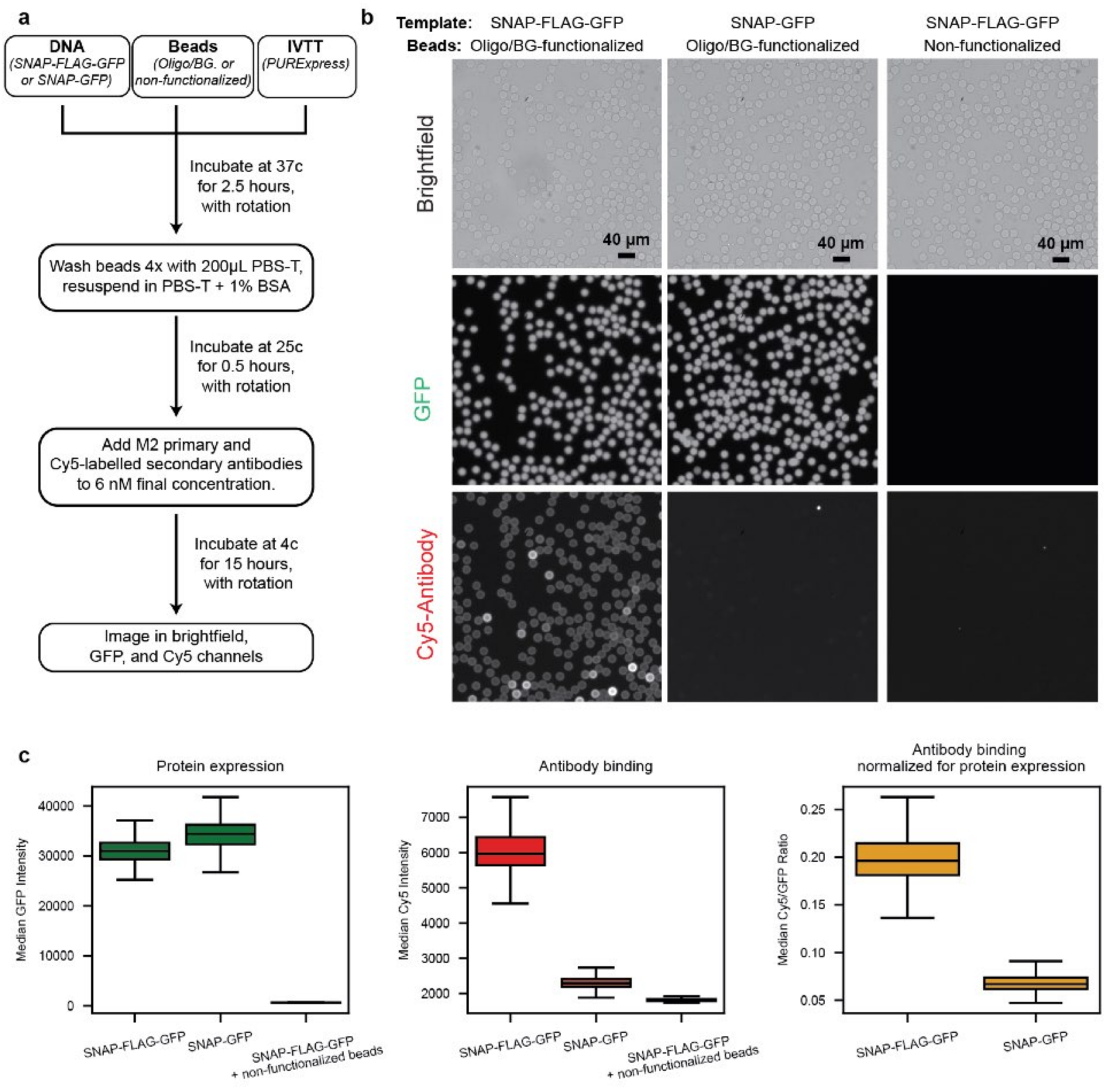


**Supplementary Figure 7 — Nonspecific binding controls for bead attachment and antibody staining.**

(a) Experimental procedure for testing: (1) if expressed proteins attach to beads via a SNAP/BG linkage and (2) if fluorescently-labeled M2 antibody and Cy5 secondary antibody bind specifically to FLAG variants displayed from beads. (b) Microscopy images of antibody-stained beads in bright field, GFP (300 ms), and Cy5 (300 ms) fluorescence channels. Scale bar = 40 µm. (c) Quantification of Cy5 and GFP intensities from the microscopy images in panel b. Left: per-bead GFP intensities; Middle: per-bead Cy5 intensities; Right: per-bead Cy5/GFP intensities (non-functionalized bead sample excluded due to lack of GFP signal).

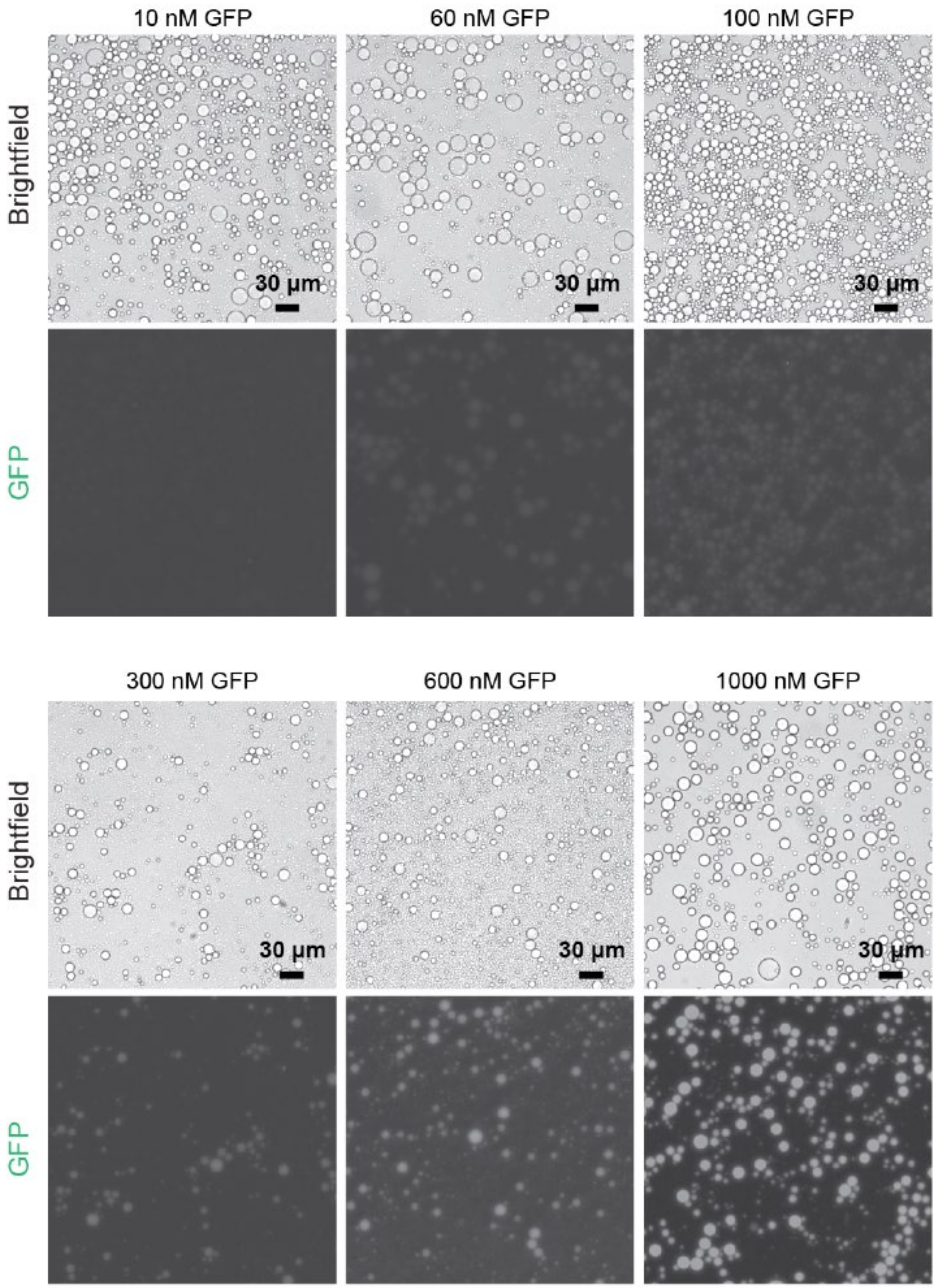


**Supplementary Figure 8 — Representative images of vortexed emulsions loaded with increasing concentrations of eGFP protein.**

Representative microscopy images of vortexed emulsions loaded with increasing concentrations of recombinant eGFP protein. Top rows: bright field images. Bottom rows: GFP channel images (300 ms exposure). Scale bar = 30 µm.

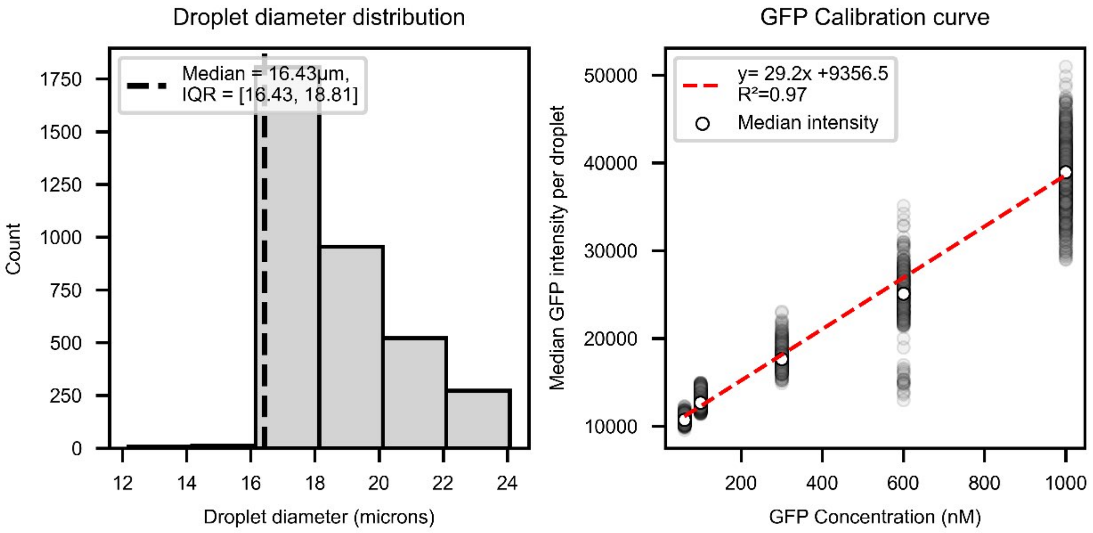


**Supplementary Figure 9 — Calibration curve for converting GFP intensities into concentrations.**

Left: Distribution of GFP-loaded droplet diameters used for concentration calibration. Right: Measured median eGFP intensity per droplet as a function of loaded GFP concentration. Gray markers indicate per-droplet intensities; white makers indicate the median per-droplet intensity at each GFP concentration; red dashed line indicates linear regression fit used to derive a calibration curve relating measured intensity to encapsulated concentration.

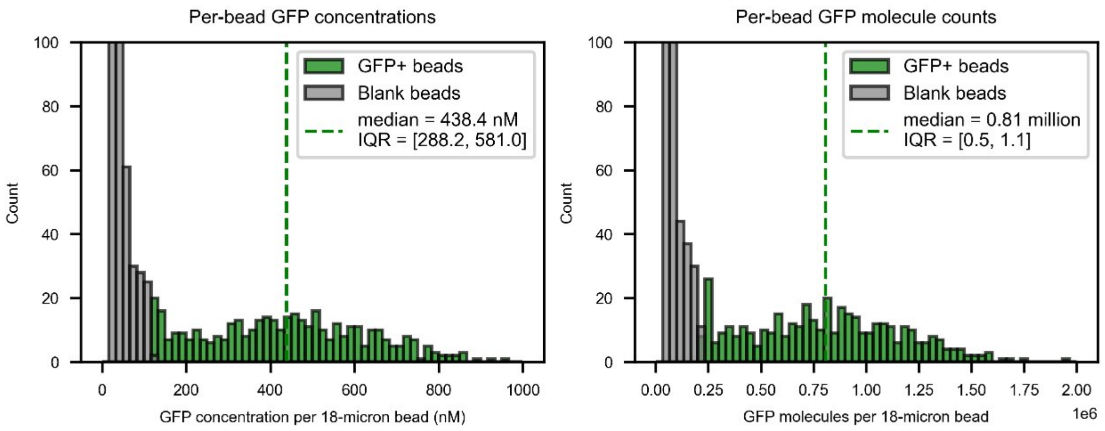


**Supplementary Figure 10 — Distribution of estimated per-bead GFP concentration and molecule counts for the 25-variant library.**

Distribution of estimated per-bead GFP concentration (left) and displayed protein molecule count (right) based on the GFP calibration curve in Supplementary Figure 9. Gray bars: GFP-negative beads; green bars: GFP-positive beads. Green dotted lines: median values for the GFP-positive bead population.

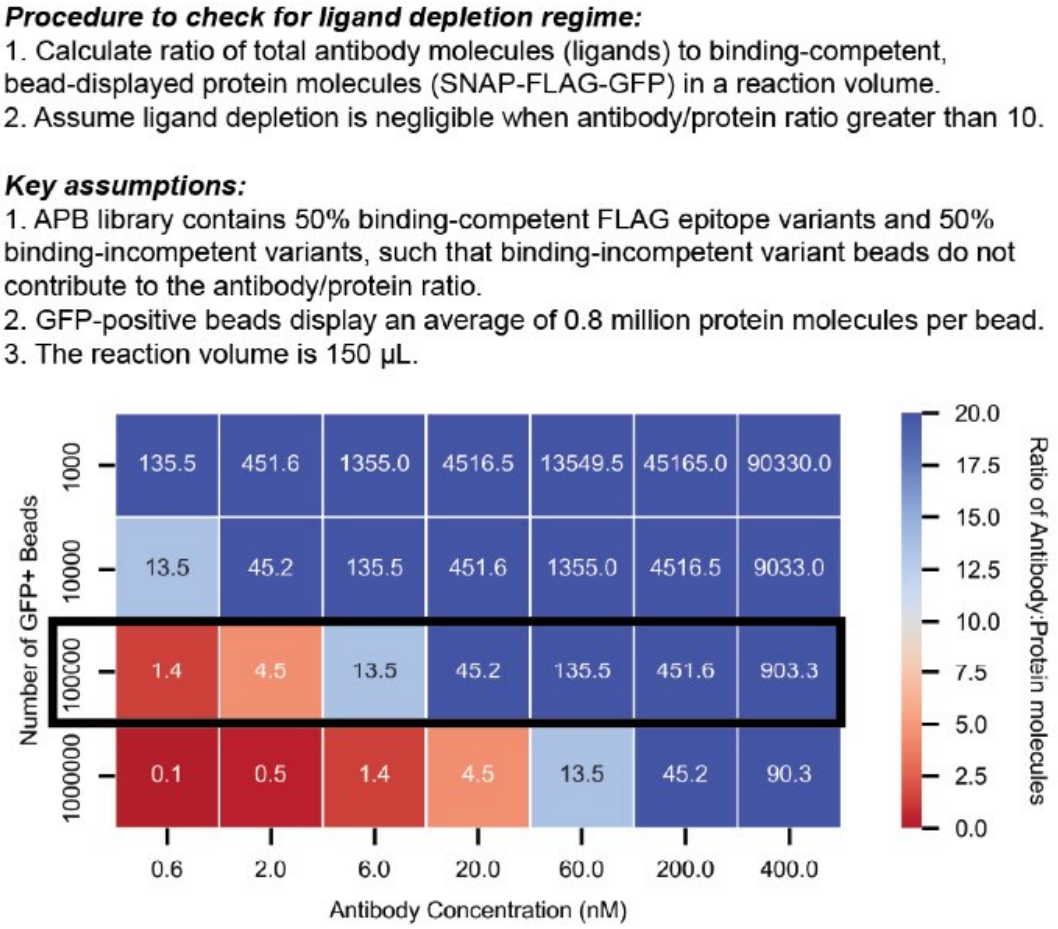


**Supplementary Figure 11 — Calculations to test if reaction conditions lead to a ligand depletion regime.**
Heat map showing calculated antibody/protein molecule ratios as a function of the number of GFP-positive beads and introduced antibody concentration within a 150 µL reaction volume. Estimated per-bead protein numbers were based on measurements shown in Supplementary Figure 10. Red colors indicate a likely titration regime and blue colors indicate conditions outside of this regime. Black box indicates the typical reaction conditions used in this paper; ligand concentrations at or above 6 nM should not result in ligand depletion.

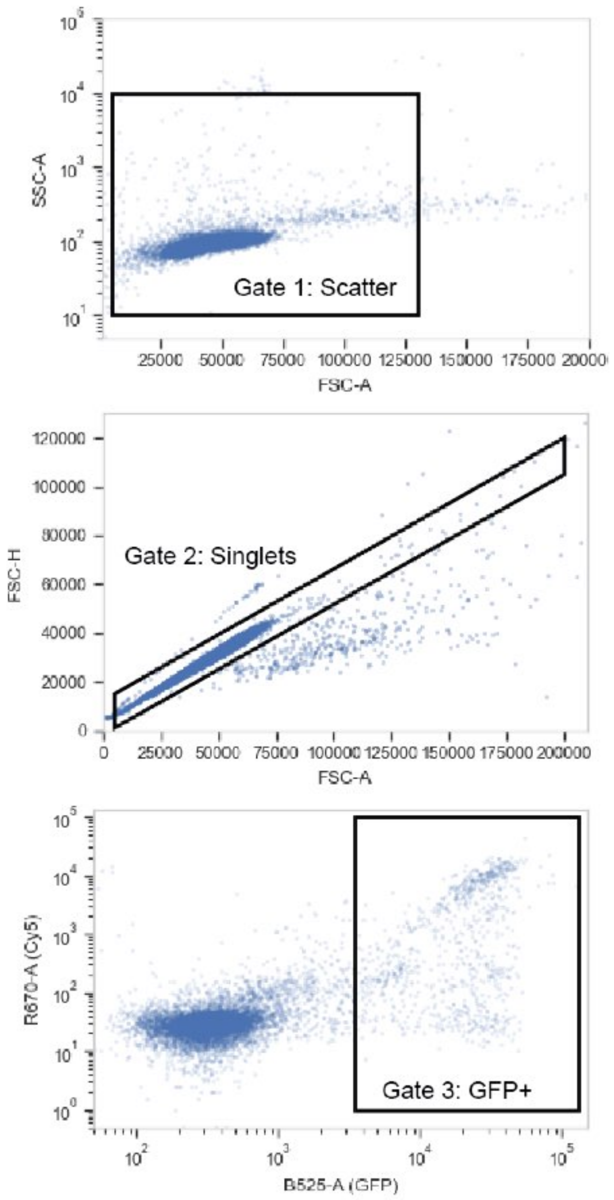


**Supplementary Figure 12 — FACS gating scheme for identifying singlet GFP-positive Amplicon/Protein beads for the 25-variant library.**

FACS plots showing gating scheme for identifying singlet GFP-positive Amplicon/Protein beads. Top: Scatter gate to exclude non-bead particles. Middle: Singlet gate to exlude bead clumps. Bottom: GFP-positive gate to exclude beads without displayed SNAP-FLAG-GFP proteins.

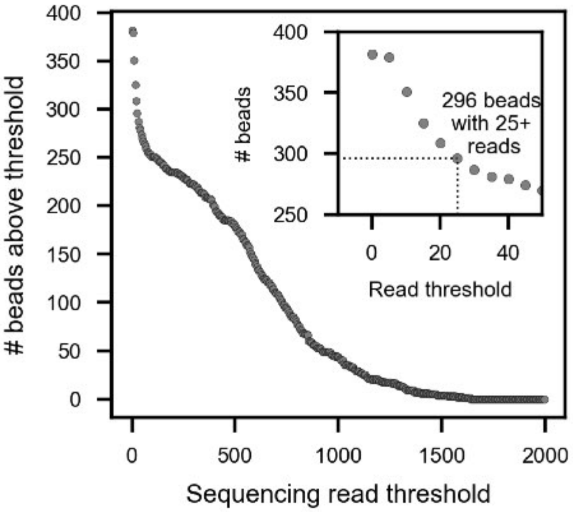


**Supplementary Figure 13 — Number of sorted single beads as a function of read count threshold for the 25 variant library.**

Number of sorted single beads with at least one variant sequence exceeding the read count threshold. Inset plot shows read counts between 0 and 50, with dotted lines marking the number of sorted beads with 25+ reads corresponding to an expected variant sequence.

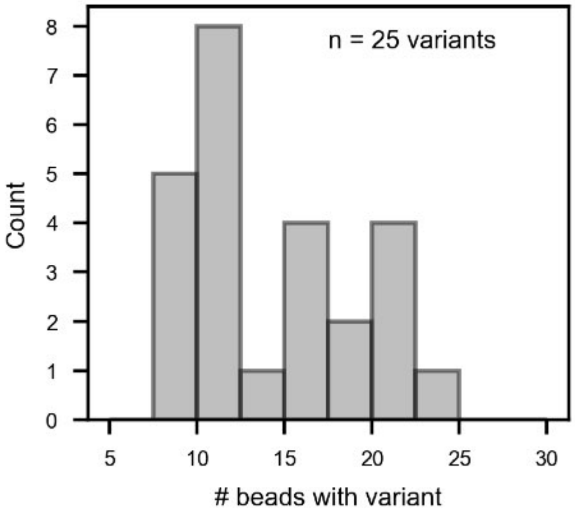


**Supplementary Figure 14 — Distribution of the number of sorted beads per variant for the 25 variant library.**

Distribution of the number of sorted beads with at least 25 reads for a single variant within the 25-variant FLAG library.

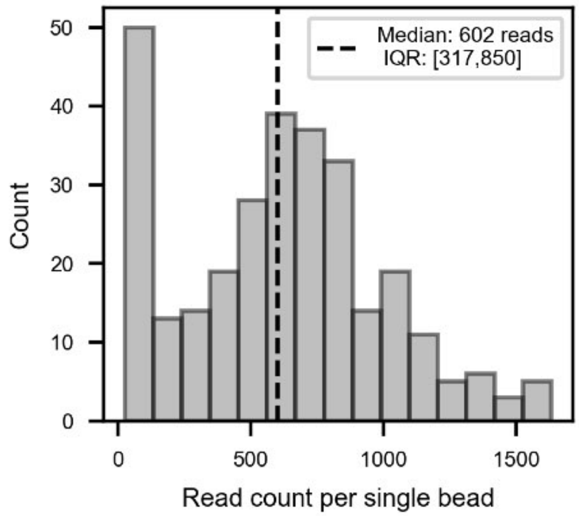


**Supplementary Figure 15 — Distribution of total read counts for beads containing variants with at least 25 reads for the 25 variant library.**

Distribution of total read counts for beads containing variants with at least 25 reads for at least 1 variant. Black dotted line: median read count.

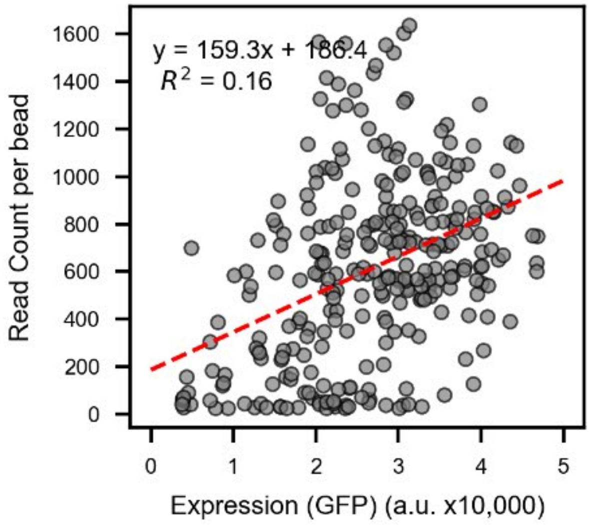


**Supplementary Figure 16 — Correlation between GFP intensity and total read counts on single beads for the 25 variant library.**

Scatter plot of GFP intensity (expression) versus total read counts on single beads with at least 25 reads for at least one variant. Red line denotes linear regression fit with annotated fit parameters.

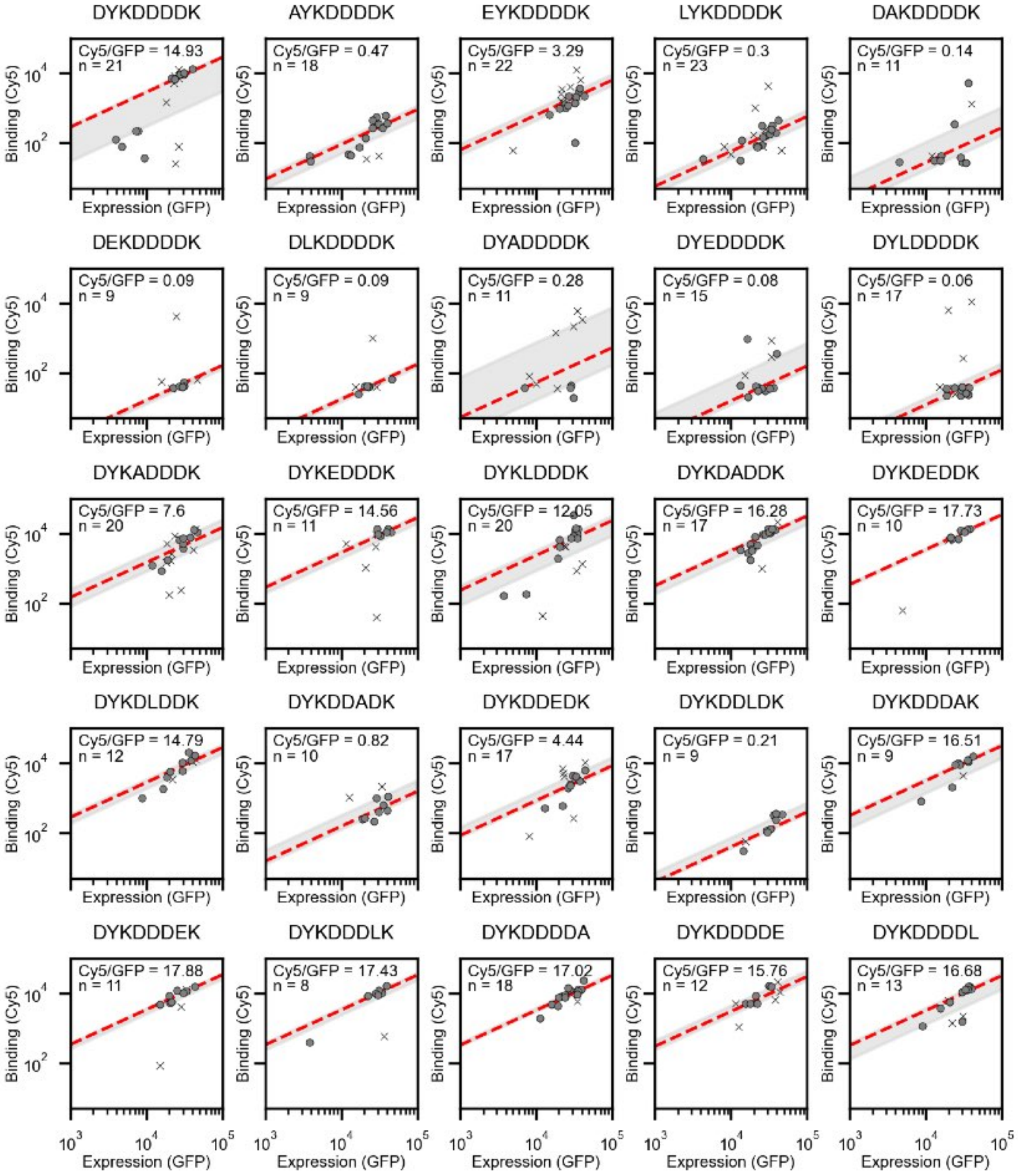


**Supplementary Figure 17 — Measured Cy5 versus GFP intensities for all beads with at least 25 reads corresponding to a given variant for the 25 variant library.**

Scatter plots showing measured Cy5 intensity versus measured GFP intensity for all beads displaying a given variant. Circles indicate beads displaying a single variant; X's indicate beads displaying more than one variant. The red line indicates the median Cy5/GFP ratio across all beads displaying a given variant and the gray fill indicates the interquartile range [0.25,0.75].

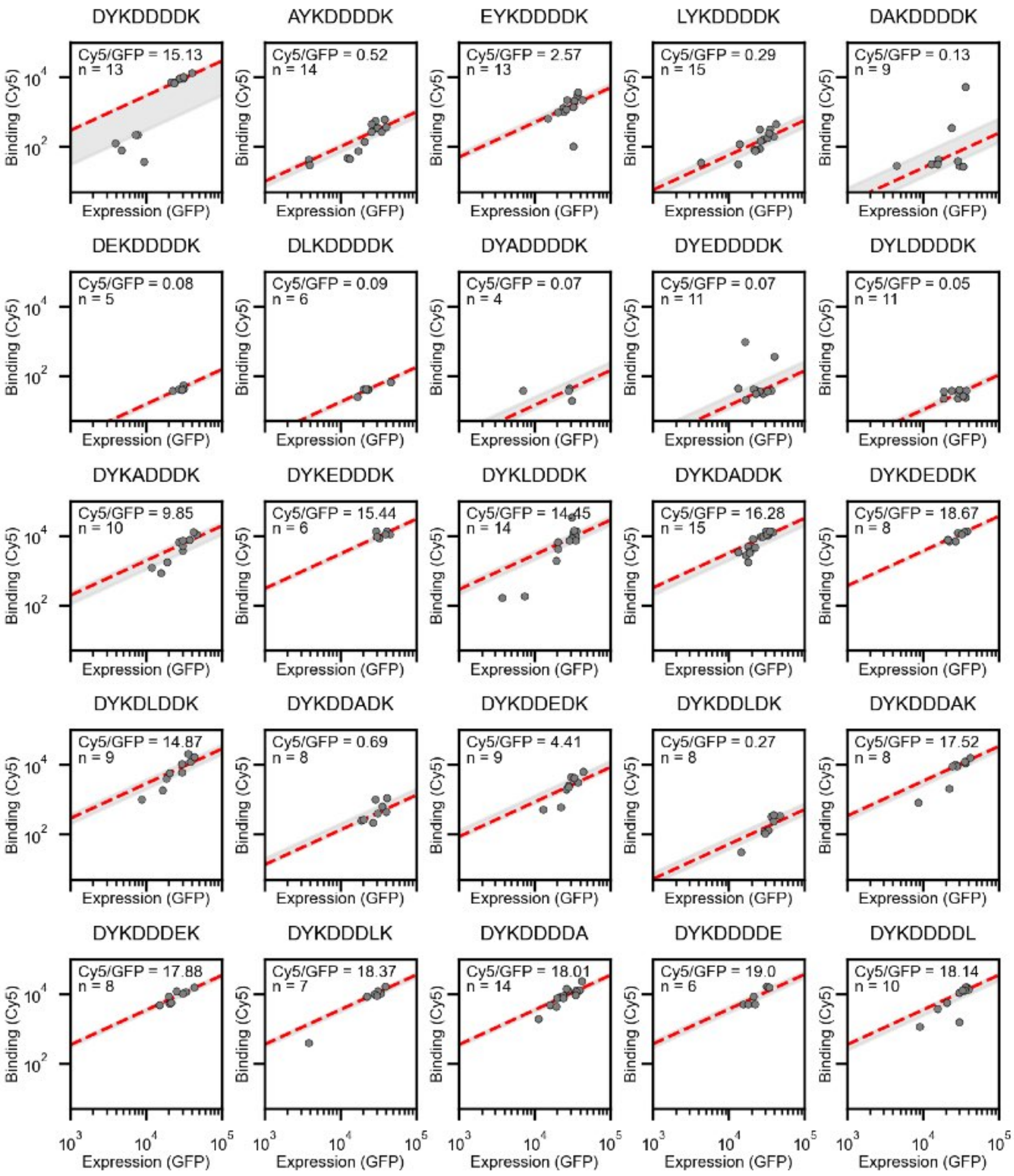


**Supplementary Figure 18 — Measured Cy5 versus GFP intensities for all beads with at least 25 reads corresponding to only a single variant for the 25 variant library.**

Scatter plots showing measured Cy5 intensity versus measured GFP intensity for all beads displaying only a single variant. Circles indicate beads displaying a single variant. The red line indicates the median Cy5/GFP ratio across all beads displaying a single variant and the gray fill indicates the interquartile range [0.25,0.75].

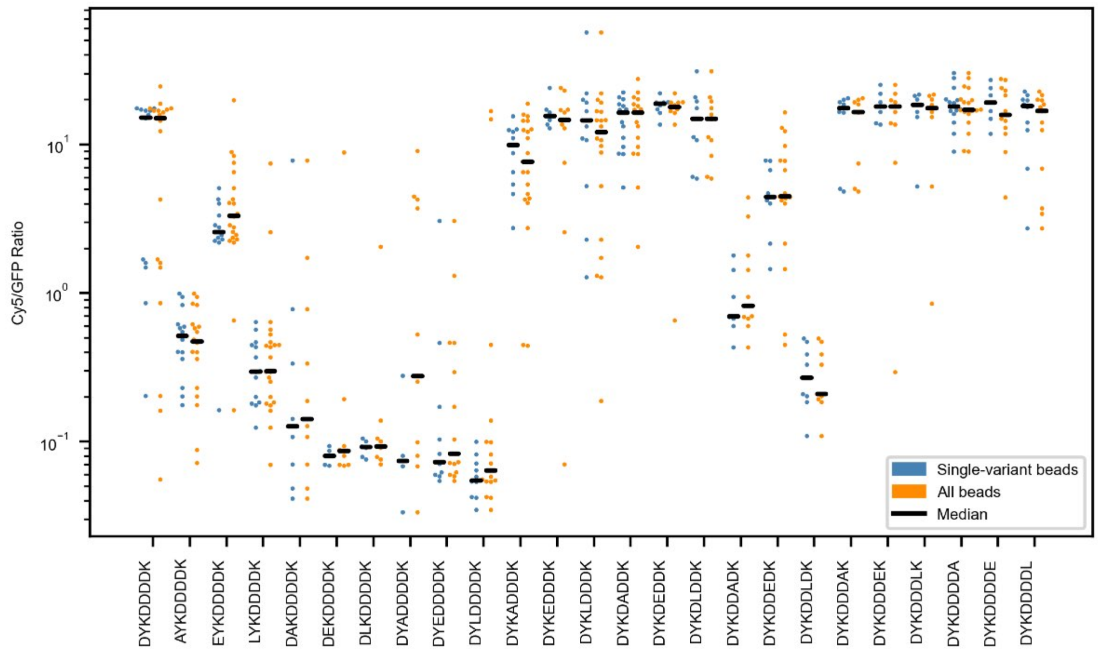


**Supplementary Figure 19 — Comparison of Cy5/GFP ratio distributions for beads displaying only a single variant and all beads.**

Distribution of per-bead Cy5/GFP ratios corresponding to a given variant sequence for beads containing only a single variant (blue) or all beads containing the given variant (orange). Black lines indicate population medians.

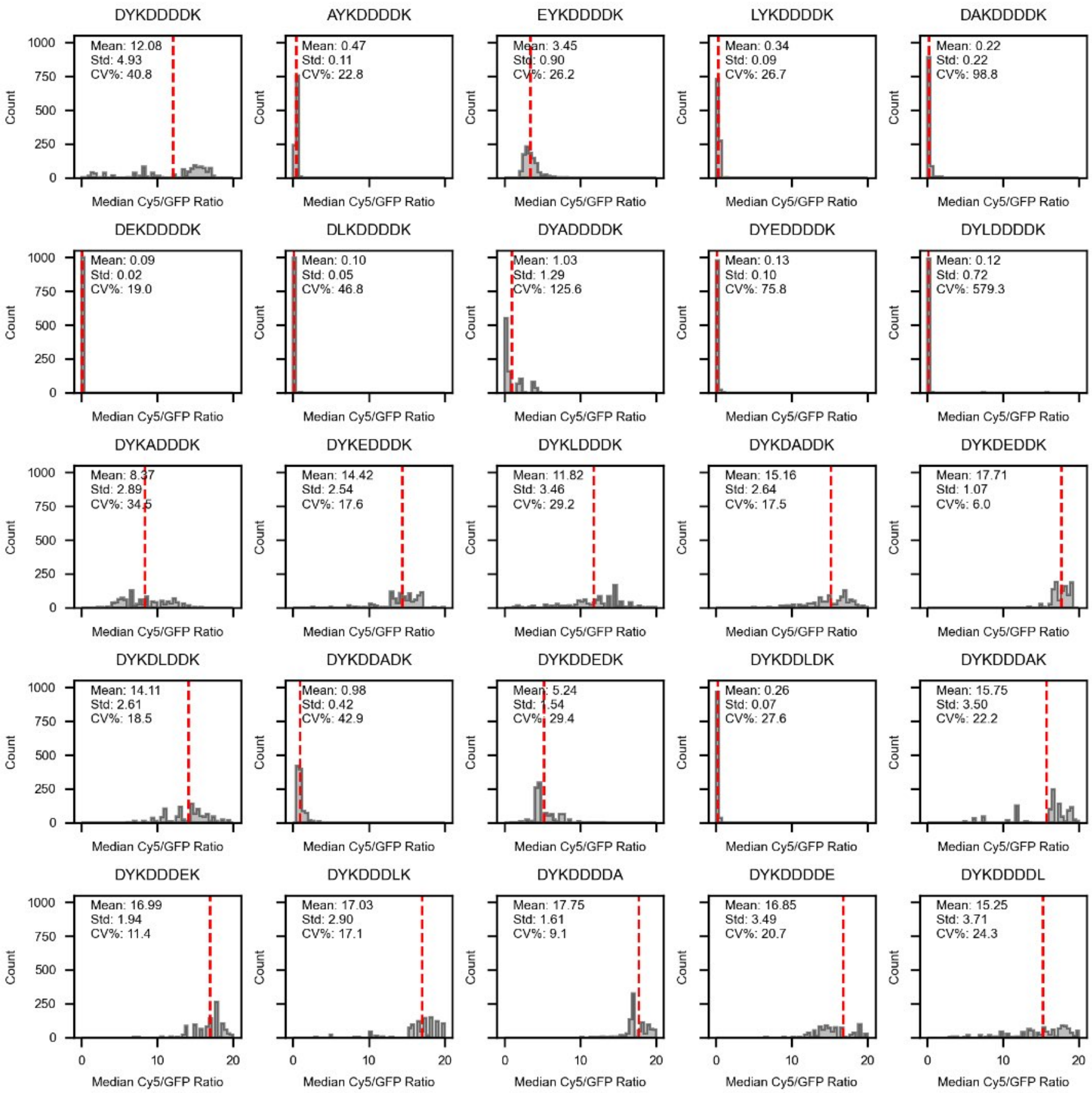


**Supplementary Figure 20 — Bootstrap-sampled median Cy5/GFP distributions for 10-bead APB populations for the 25-variant library.**

Distribution of bootstrap-sampled median Cy5/GFP values for 10-bead APB populations. Red dashed line indicates mean calculated from 1000 sampling iterations

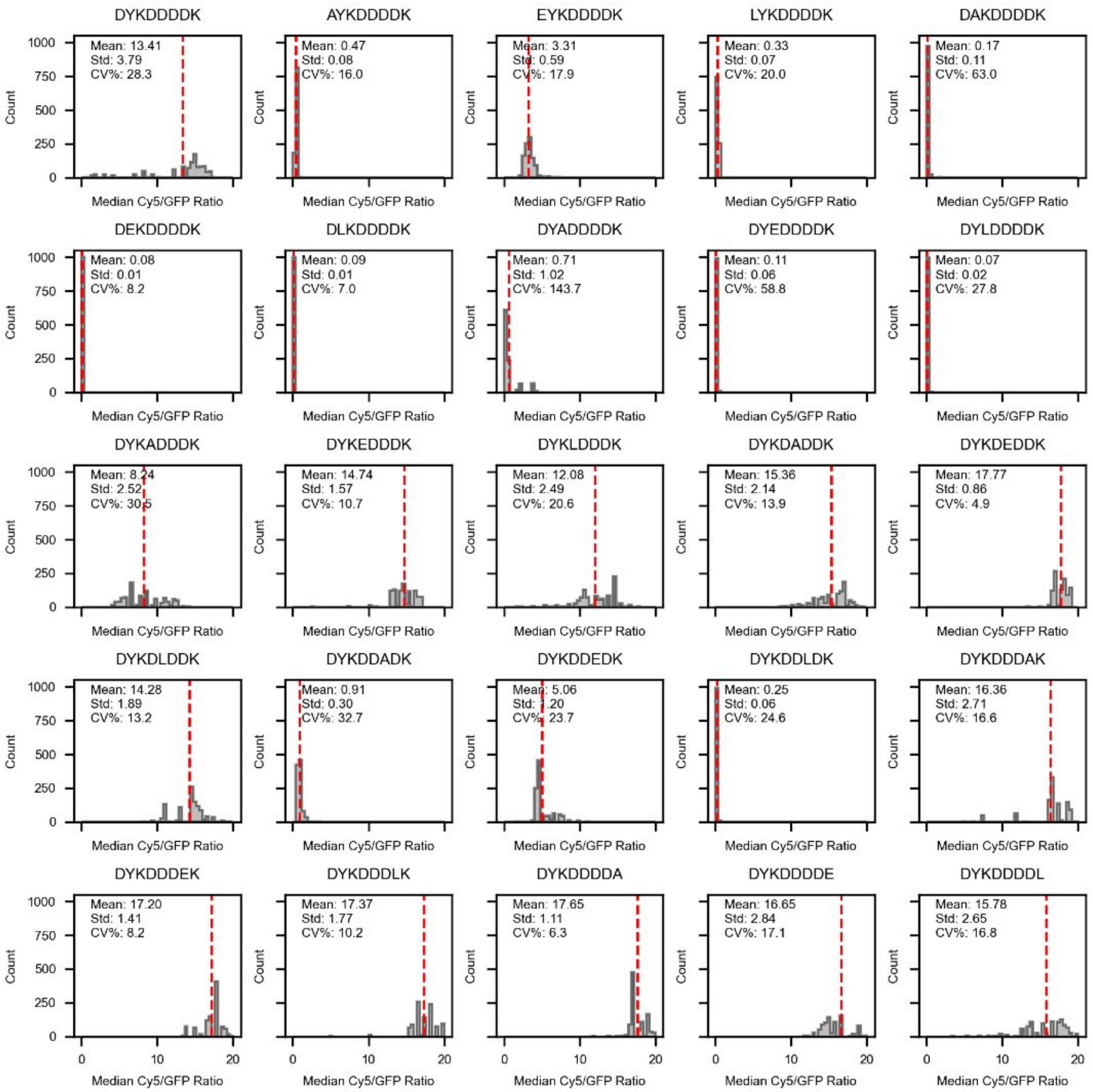


**Supplementary Figure 21 — Bootstrap-sampled median Cy5/GFP distributions for 20-bead APB populations for the 25-variant library.**

Distribution of bootstrap-sampled median Cy5/GFP values for 20-bead APB populations. Red line: population mean. Red dashed line indicates mean calculated from 1000 sampling iterations

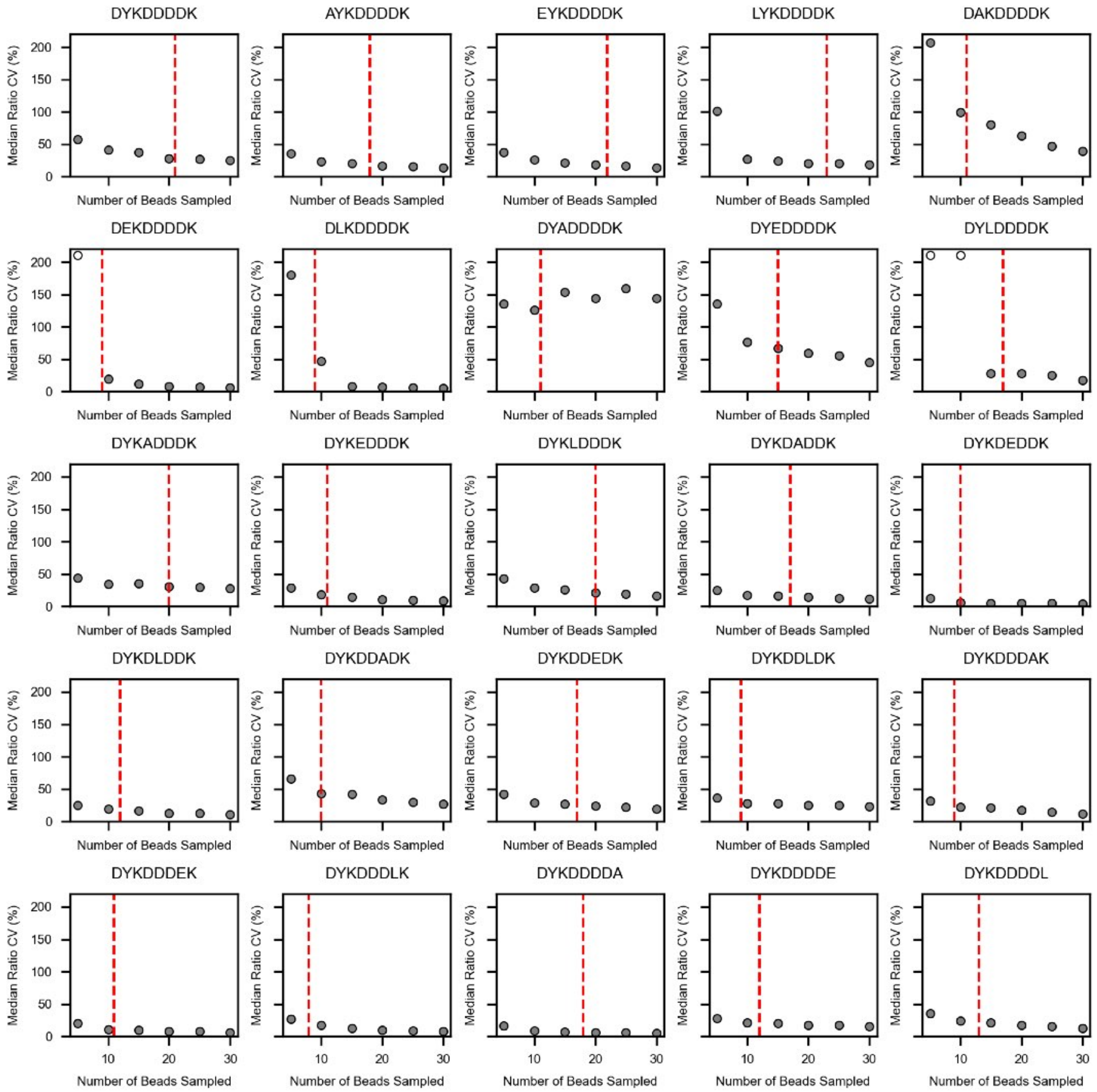


**Supplementary Figure 22 — Median Cy5/GFP CV% as a function of sampled bead count for the 25-variant library.**

Median Cy5/GFP CV% as a function of the number of beads sampled in each of 1000 bootstrap iterations for each of 25 FLAG variants. The upper limit of CV% is set to 210%; gray markers indicate values below this limit and white markers indicate values at or above this limit. Red dashed line indicates the total number of beads measured for each variant.

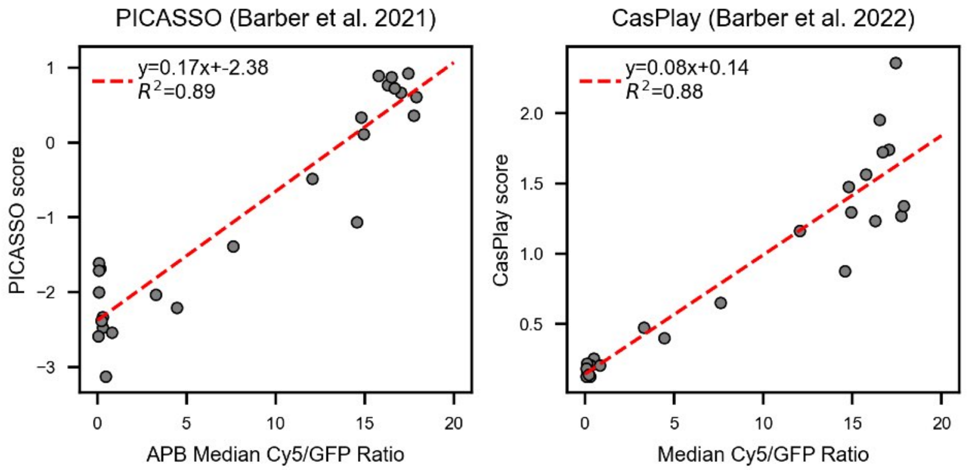


**Supplementary Figure 23 — Comparison of APB Cy5/GFP ratios from the 25-variant library with Cas9-display binding scores.**

Median Cy5/GFP ratios for 25 FLAG variants displayed on APBs versus binding score (log2(anti-FLAG/anti-His)) from PICASSO [52] (left) and binding score (output/input reads) from CasPlay [49] (right). Red dashed lines indicate linear regression fits with annotated fit parameters.

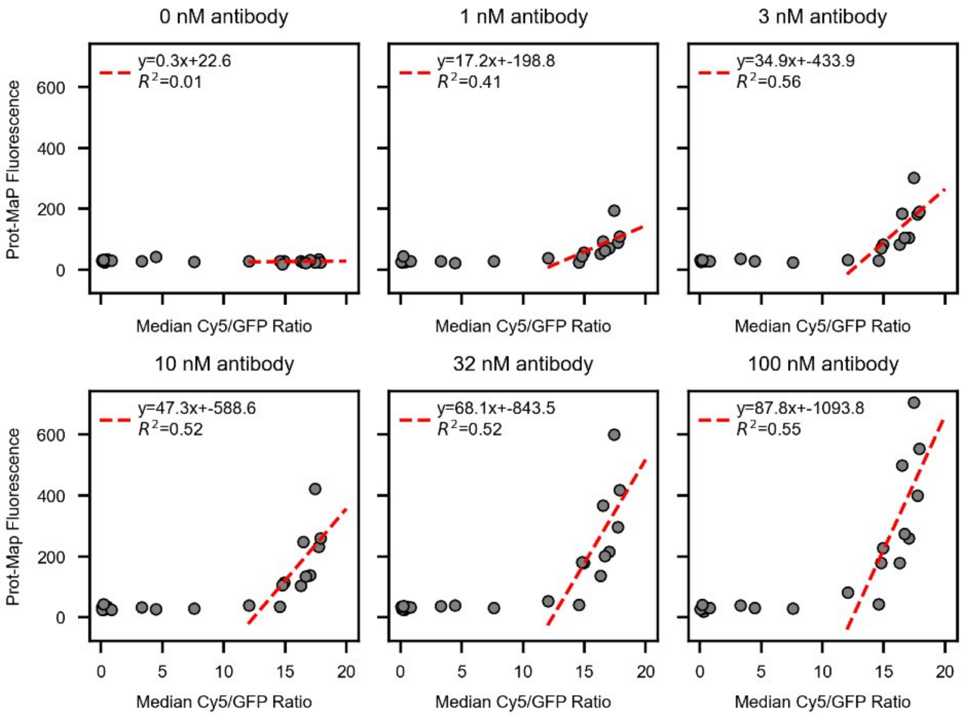


**Supplementary Figure 24 — Comparison of APB Cy5/GFP ratios from the 25-variant library with Prot-MaP fluorescence intensities.**

Median Cy5/GFP ratios for 25 FLAG variants displayed on APBs versus fluorescence intensities from Prot-MaP [50] as a function of M2 antibody concentration. Red dashed lines indicate linear regression fits. Variants with APB Cy5/GFP ratio below 10 were excluded from linear regression analysis because they fall outside the dynamic range of Prot-MaP.

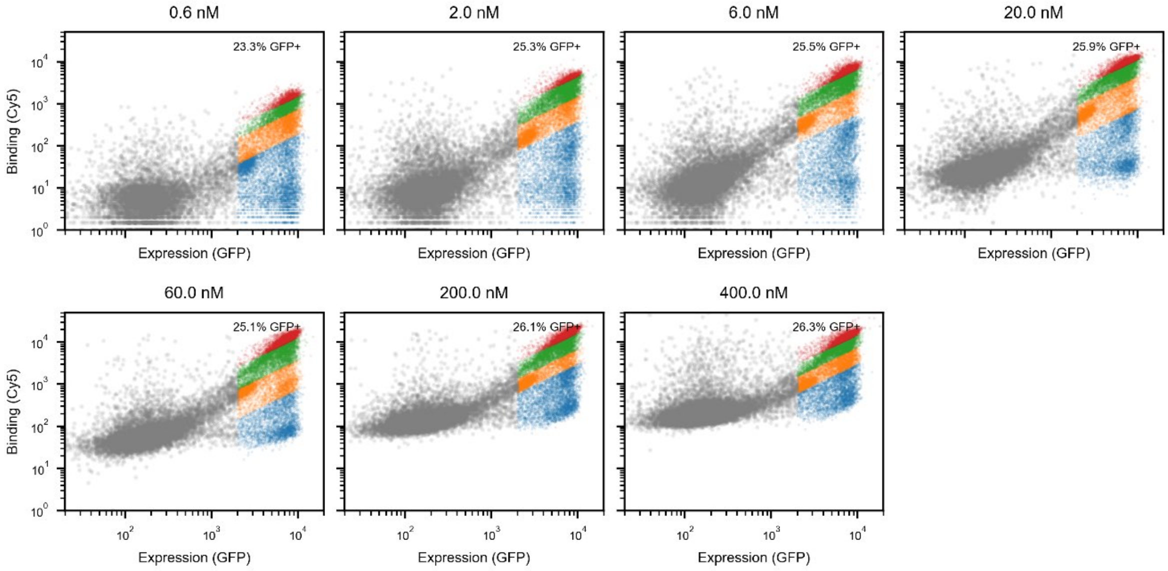


**Supplementary Figure 25 — Log-log FACS plots for the single-variant library at each antibody concentration.**

Log-log FACS plot of Cy5 versus GFP intensities for recorded beads at each antibody concentration. Gray: blank beads; blue: bin 1; orange: bin 2; green: bin 3; red: bin 4.

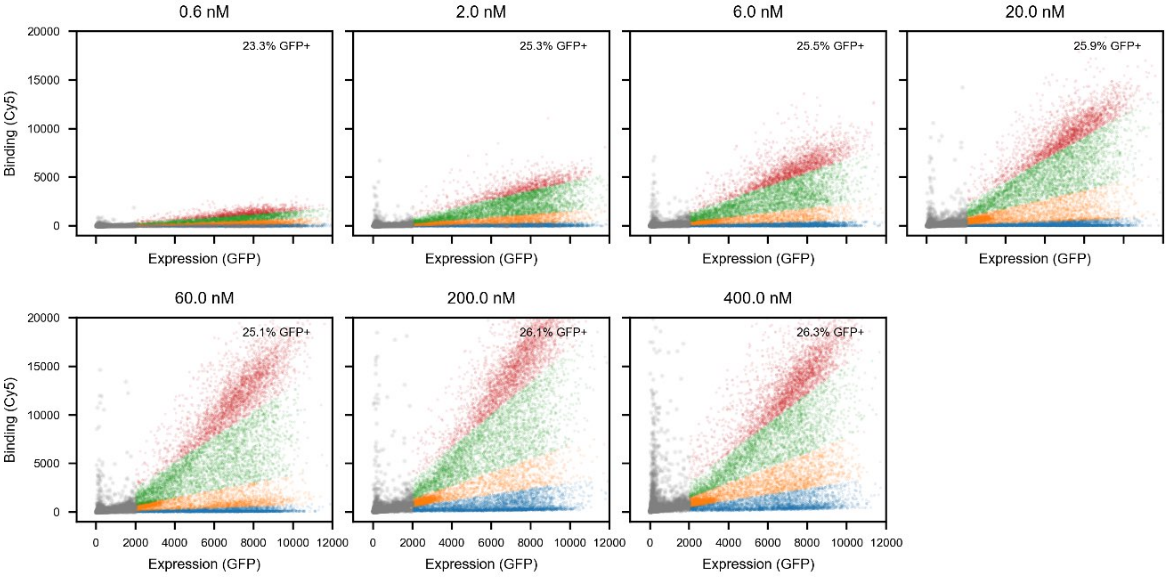


**Supplementary Figure 26 — Linear-linear FACS plots for the single-variant library at each antibody concentration.**

Linear-linear FACS plot of Cy5 versus GFP intensities for recorded beads at each antibody concentration. Gray: blank beads; blue: bin 1; orange: bin 2; green: bin 3; red: bin 4.

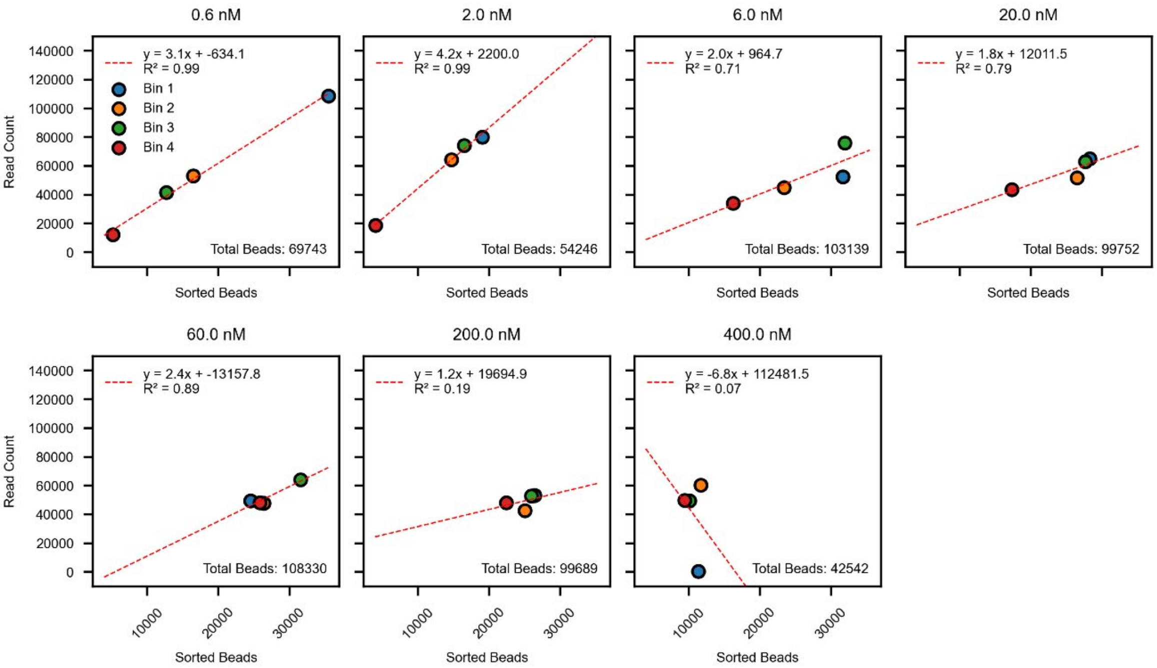


**Supplementary Figure 27 — Correlation between read counts and sorted bead counts for each concentration-bin sample in the single-variant library.**

Correlation between read counts and sorted bead counts for each sort bin and concentration. Blue: bin 1; orange: bin 2; green: bin 3; red: bin 4. Red dashed line indicates linear regression. Read counts are normalized by the number of sorted beads in each sample, such that the slope of the linear regression approximates the sequencing depth for all bins at a given concentration (*e.g.* a slope of 3.1 means that ~3.1 DNA amplicons were recovered from each sorted bead, assuming uniform amplification). In the 400 nM sample, bin 1 reads were poorly sampled relative to all other sort bins. Normalizing the read counts by the normalized beads for this sample means that each read count corresponds to >1 sorted bead.

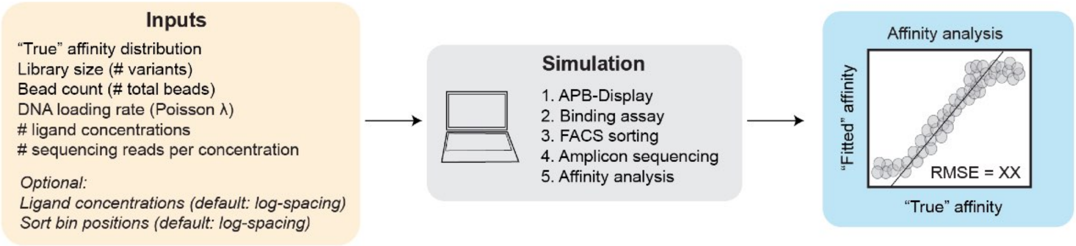


**Supplementary Figure 28 — APB-TiteSeq simulation pipeline to determine likely upper and lower limits for resolvable affinities for APB-TiteSeq experiments.**

Schematic of the APB-TiteSeq simulation pipeline showing user-defined inputs, simulated steps within the binding assay, and final comparison between apparent and true affinities.

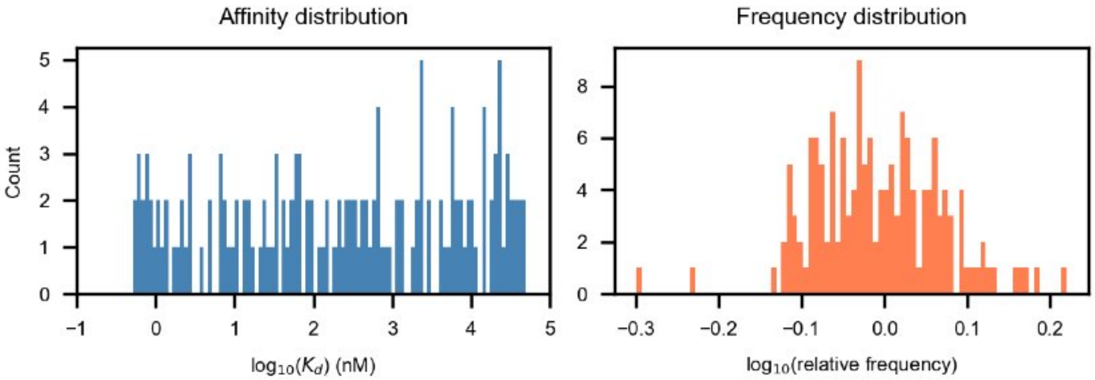


**Supplementary Figure 29 — Simulated affinity and frequency distributions used to determine likely upper and lower limits for resolvable affinities for single-variant library experiments.**

Left: Histogram of 153 log-normally-distributed affinities between 0.5nM-50µM used for simulating the single-variant library. Right: Histogram of variant frequencies in the simulated library (1.6-fold skew between 10th and 90th most abundant variants).

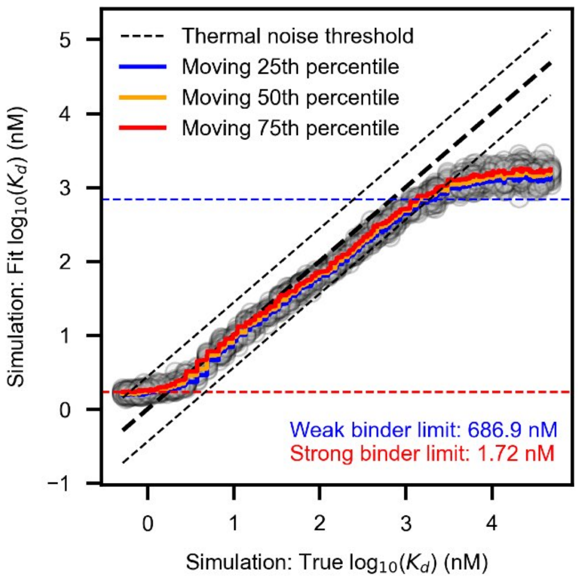


**Supplementary Figure 30 — Comparison between fitted and true affinities from simulations of the single-variant library.**

Scatter plot comparing simulated "fitted" $\log_{10}(K_d)$ values versus "true" $\log_{10}(K_d)$ values. Each point represents 1 simulation replicate and data shown include 5 simulation replicates. Black dashed lines denote thermal noise threshold (0.44 $\log_{10}(K_d)$ units, corresponding to 0.6 kcal/mol or 1 kT); colored lines denote moving percentiles of fitted $\log_{10}(K_d)$ values. We set the resolvable limit for weak and strong binders to be where the moving 25th and 75th percentiles for the $\log_{10}(K_d)$) cross the lower (blue dashed line) and upper (red dashed line) thermal noise thresholds, respectively.

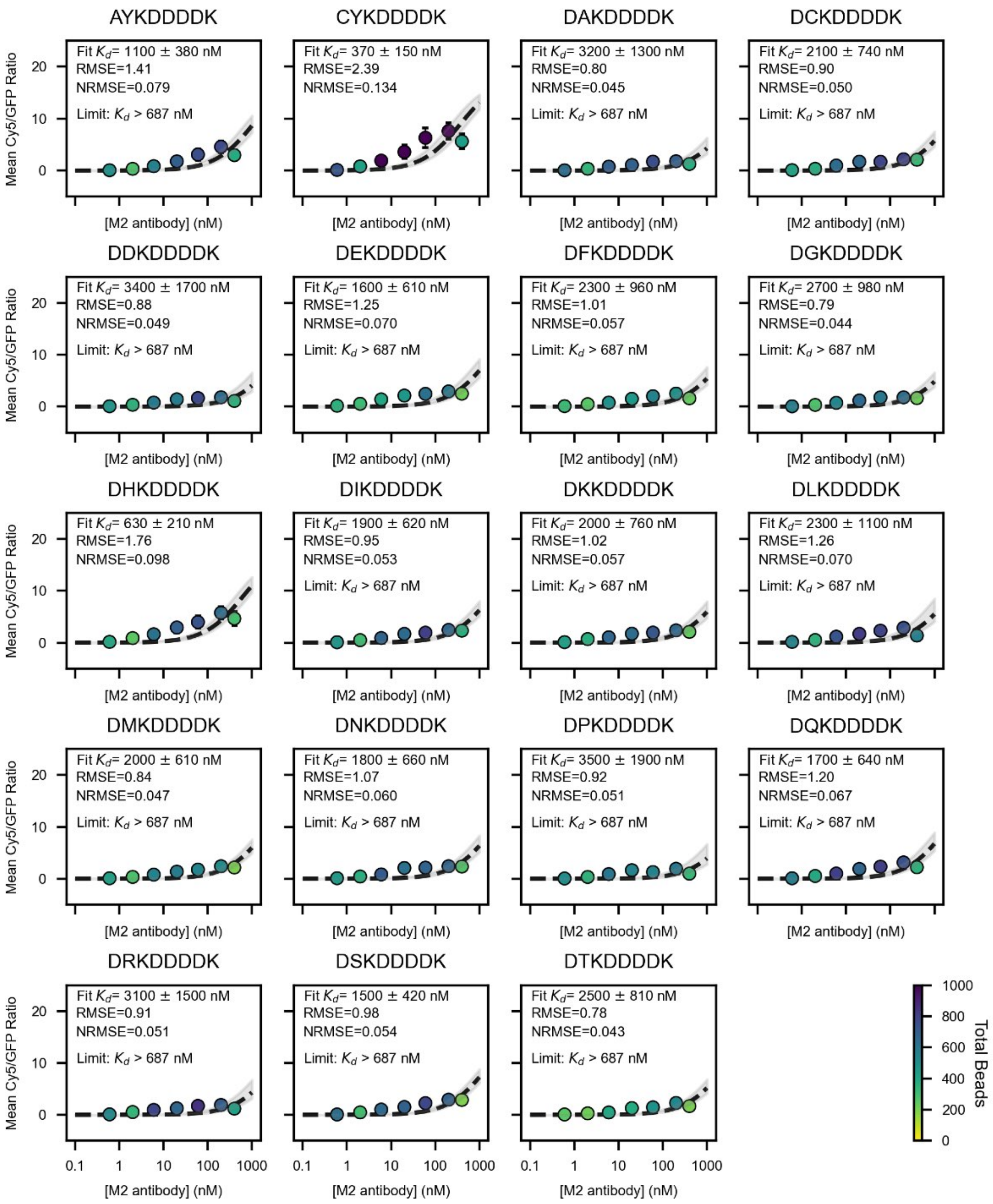


**Supplementary Figure 31 — Binding isotherms for single-variant library (page 1/8).**

Mean Cy5/GFP ratios (mean ± SD) as a function of M2 antibody concentration for all FLAG variants within the single-variant library. Gray fit curves (mean ± SD) show global fits to a Langmiur isotherm with a shared maximum saturation value (global $R_{max}$) and individual fitted $K_d$ value for each variant. Marker colors denote the number of sorted beads at each concentration; annotations report raw fitted $K_d$ values and whether measurements could be considered lower or upper limits.

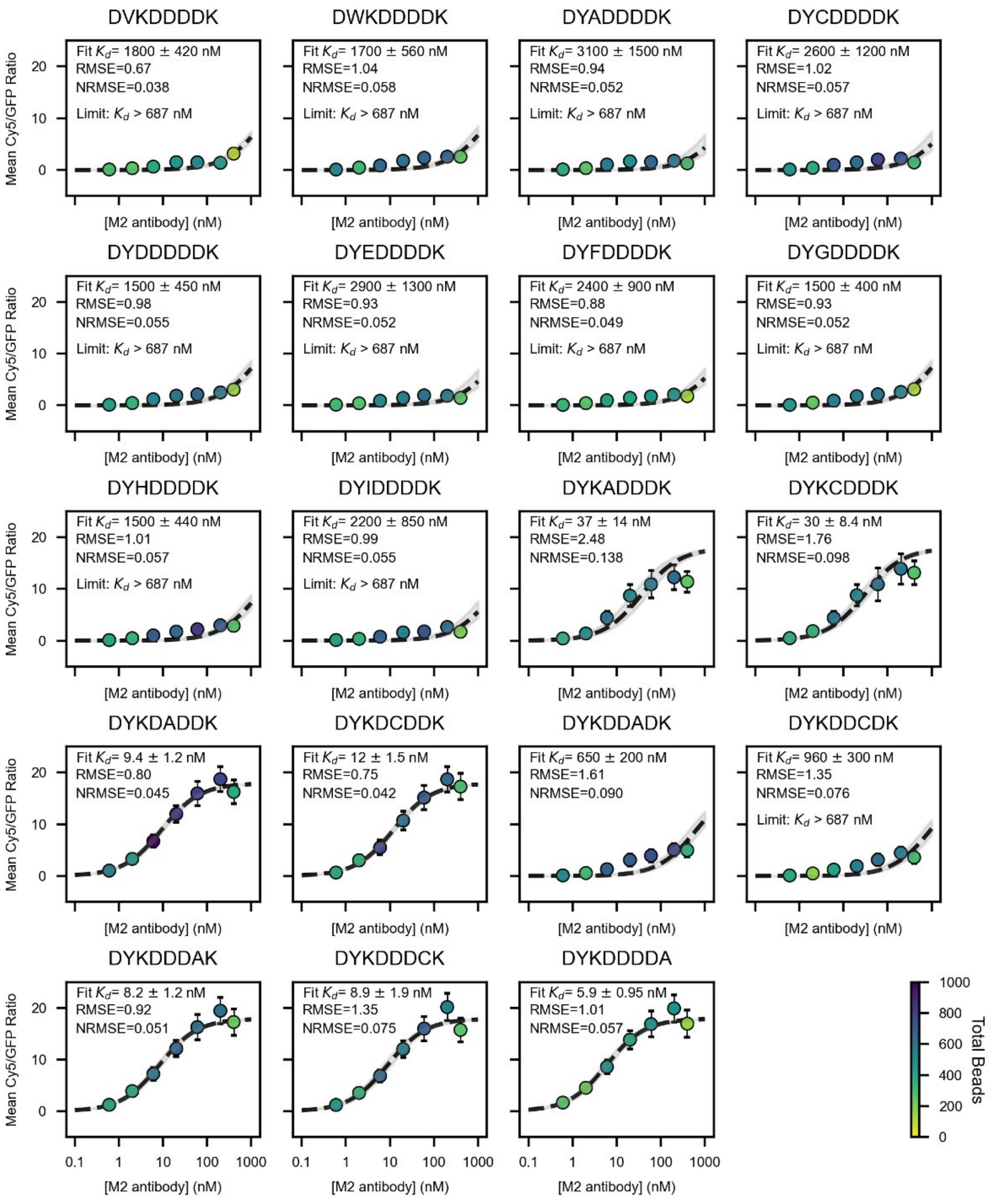


**Supplementary Figure 32 — Binding isotherms for single-variant library (page 2/8).**

Mean Cy5/GFP ratios (mean ± SD) as a function of M2 antibody concentration for all FLAG variants within the single-variant library. Gray fit curves (mean ± SD) show global fits to a Langmiur isotherm with a shared maximum saturation value (global $R_{max}$) and individual fitted $K_d$ value for each variant. Marker colors denote the number of sorted beads at each concentration; annotations report raw fitted $K_d$ values and whether measurements could be considered lower or upper limits.

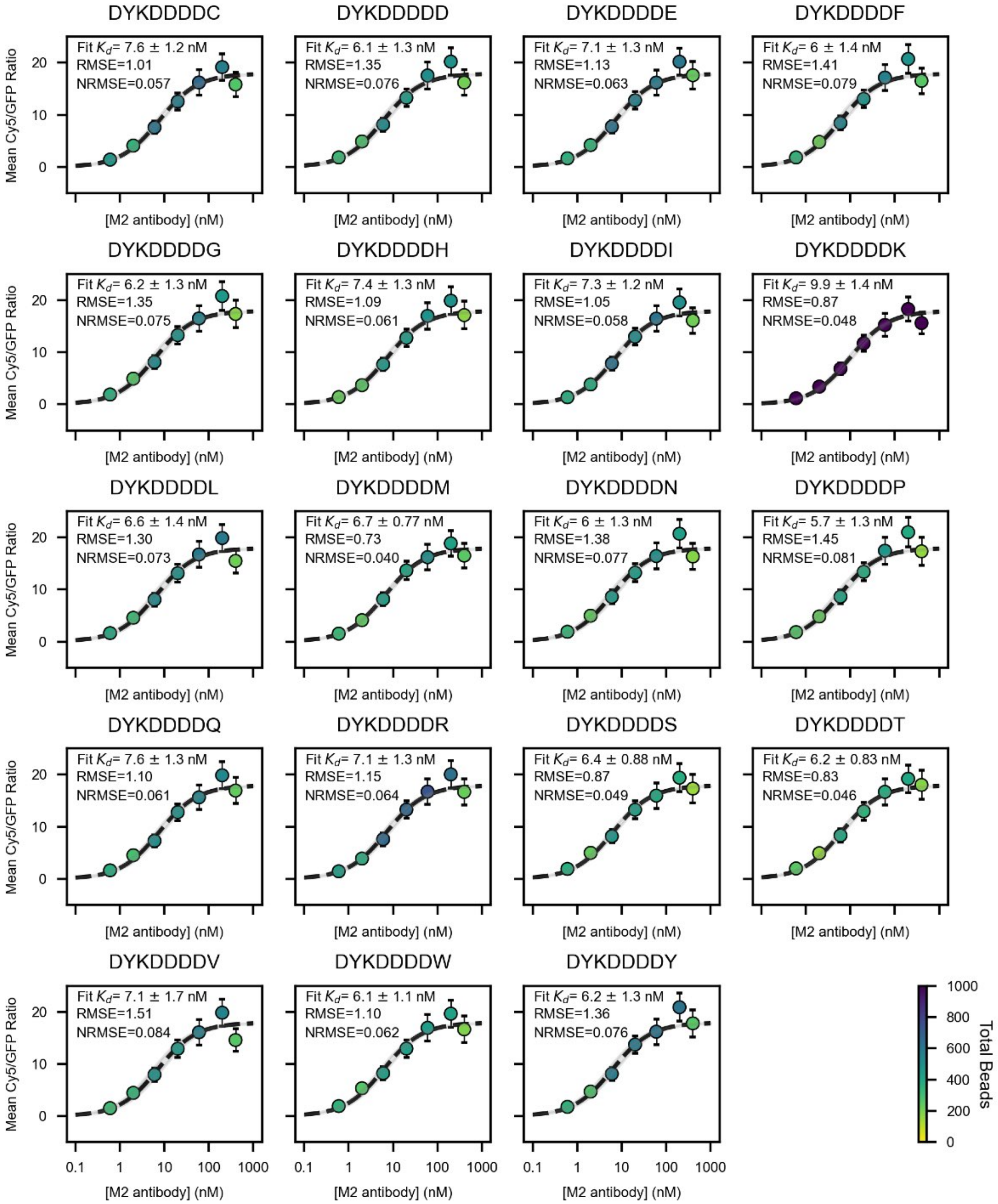


**Supplementary Figure 33 — Binding isotherms for single-variant library (page 3/8).**

Mean Cy5/GFP ratios (mean ± SD) as a function of M2 antibody concentration for all FLAG variants within the single-variant library. Gray fit curves (mean ± SD) show global fits to a Langmiur isotherm with a shared maximum saturation value (global $R_{max}$) and individual fitted $K_d$ value for each variant. Marker colors denote the number of sorted beads at each concentration; annotations report raw fitted $K_d$ values and whether measurements could be considered lower or upper limits.

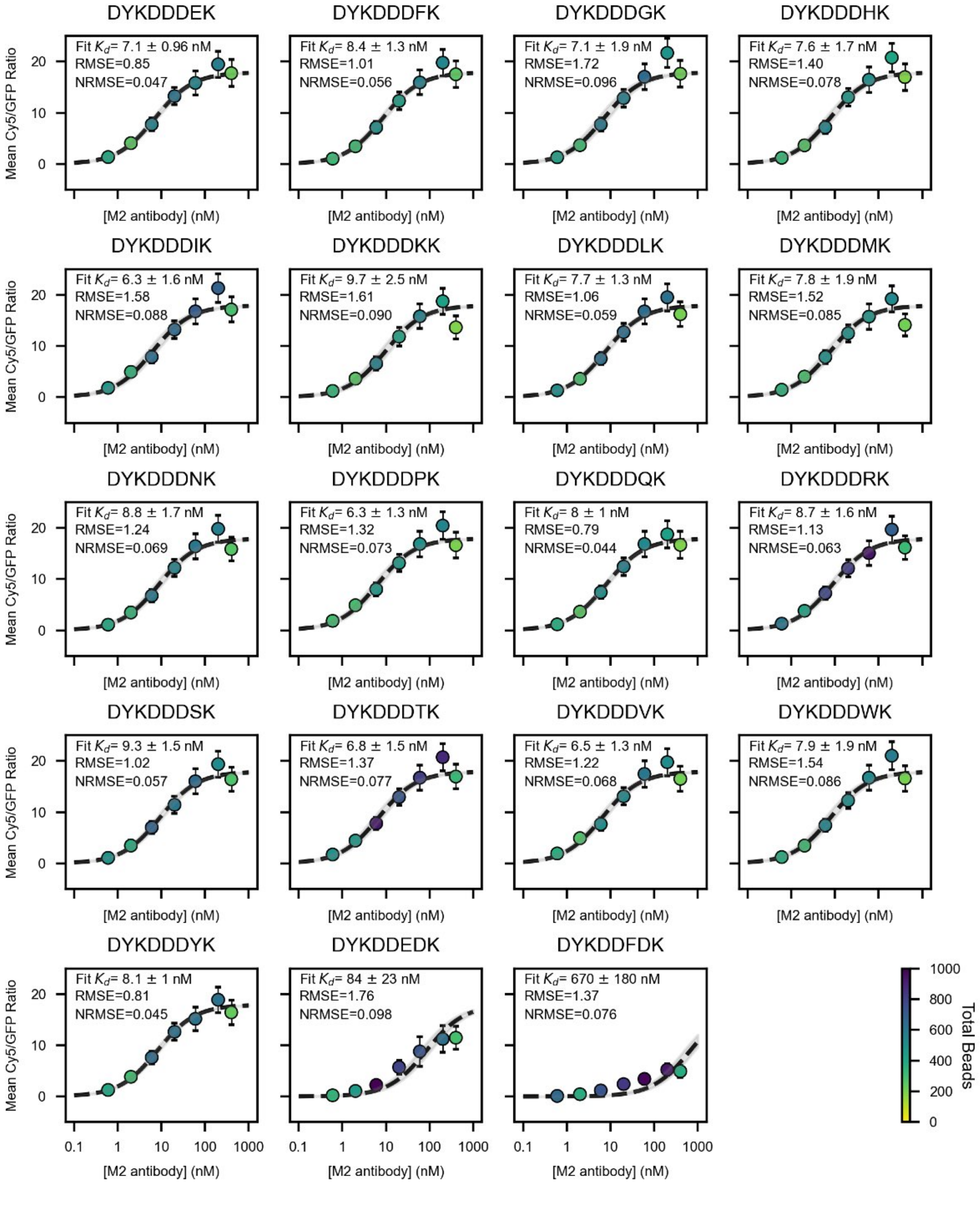


**Supplementary Figure 34 — Binding isotherms for single-variant library (page 4/8).**

Mean Cy5/GFP ratios (mean ± SD) as a function of M2 antibody concentration for all FLAG variants within the single-variant library. Gray fit curves (mean ± SD) show global fits to a Langmuir isotherm with a shared maximum saturation value (global $R_{max}$) and individual fitted $K_d$ value for each variant. Marker colors denote the number of sorted beads at each concentration; annotations report raw fitted $K_d$ values and whether measurements could be considered lower or upper limits.

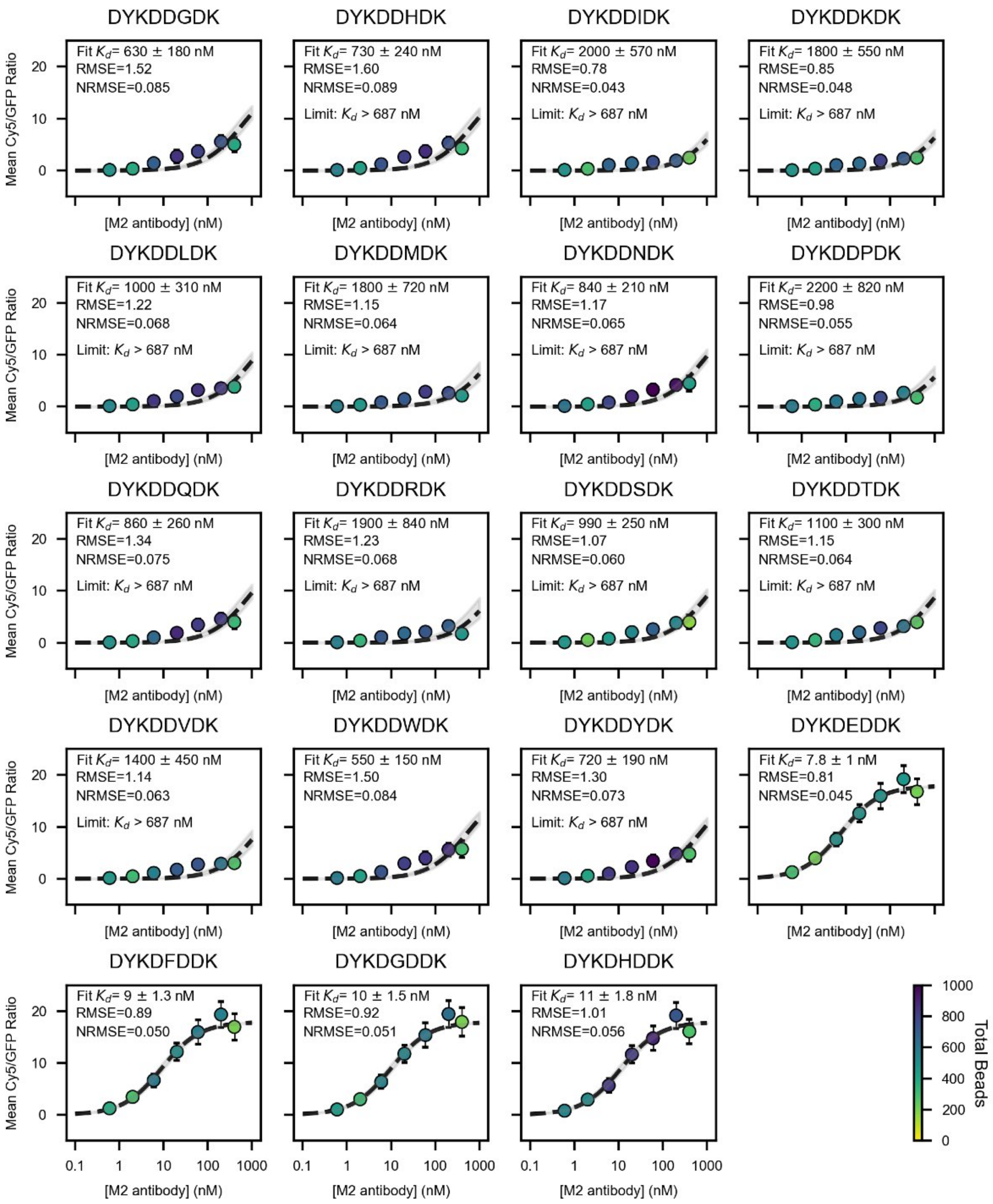


**Supplementary Figure 35 — Binding isotherms for single-variant library (page 5/8).**

Mean Cy5/GFP ratios (mean ± SD) as a function of M2 antibody concentration for all FLAG variants within the single-variant library. Gray fit curves (mean ± SD) show global fits to a Langmuir isotherm with a shared maximum saturation value (global $R_{max}$) and individual fitted $K_d$ value for each variant. Marker colors denote the number of sorted beads at each concentration; annotations report raw fitted $K_d$ values and whether measurements could be considered lower or upper limits.

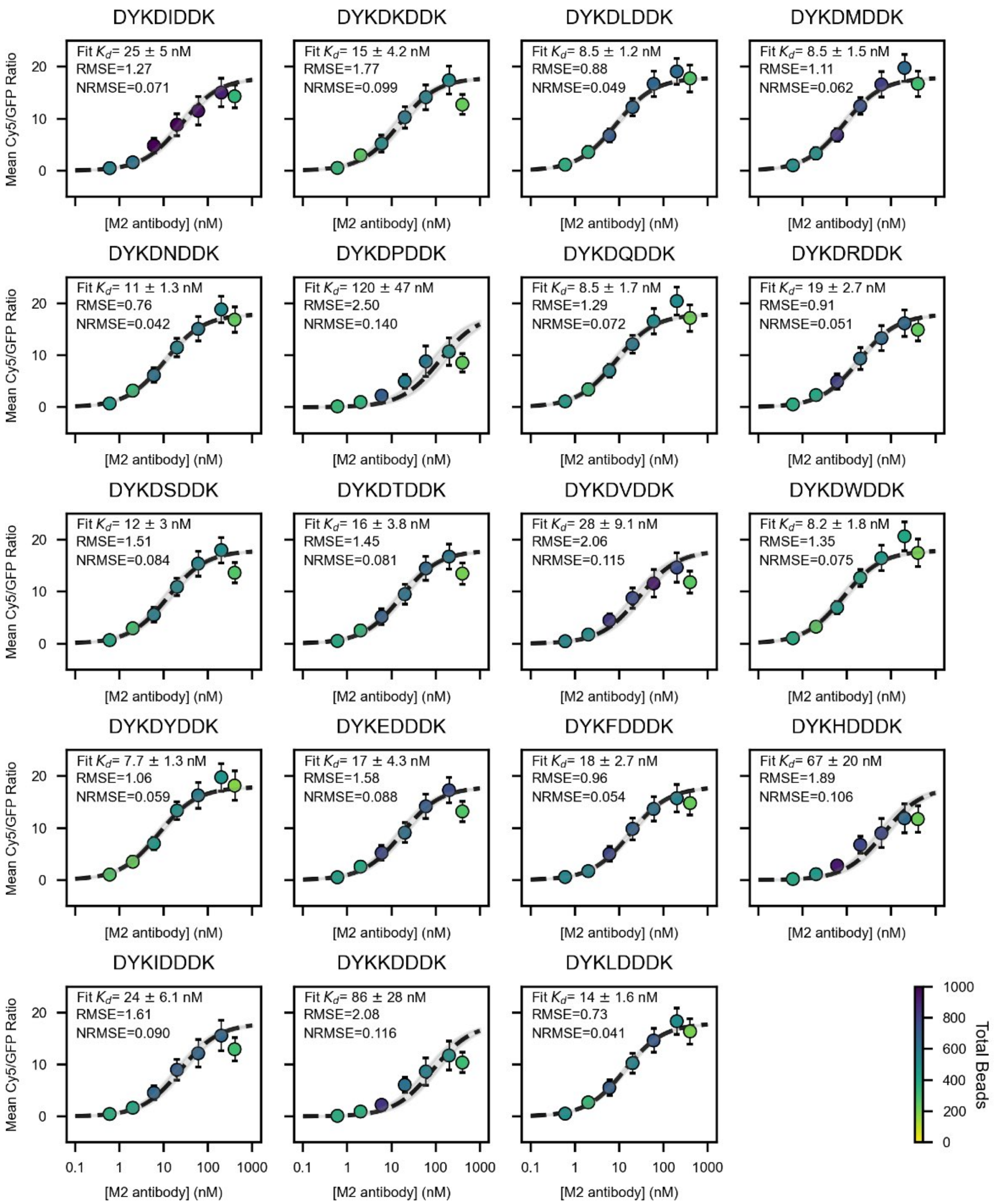


**Supplementary Figure 36 — Binding isotherms for single-variant library (page 6/8).**

Mean Cy5/GFP ratios (mean ± SD) as a function of M2 antibody concentration for all FLAG variants within the single-variant library. Gray fit curves (mean ± SD) show global fits to a Langmuir isotherm with a shared maximum saturation value (global $R_{max}$) and individual fitted $K_d$ value for each variant. Marker colors denote the number of sorted beads at each concentration; annotations report raw fitted $K_d$ values and whether measurements could be considered lower or upper limits.

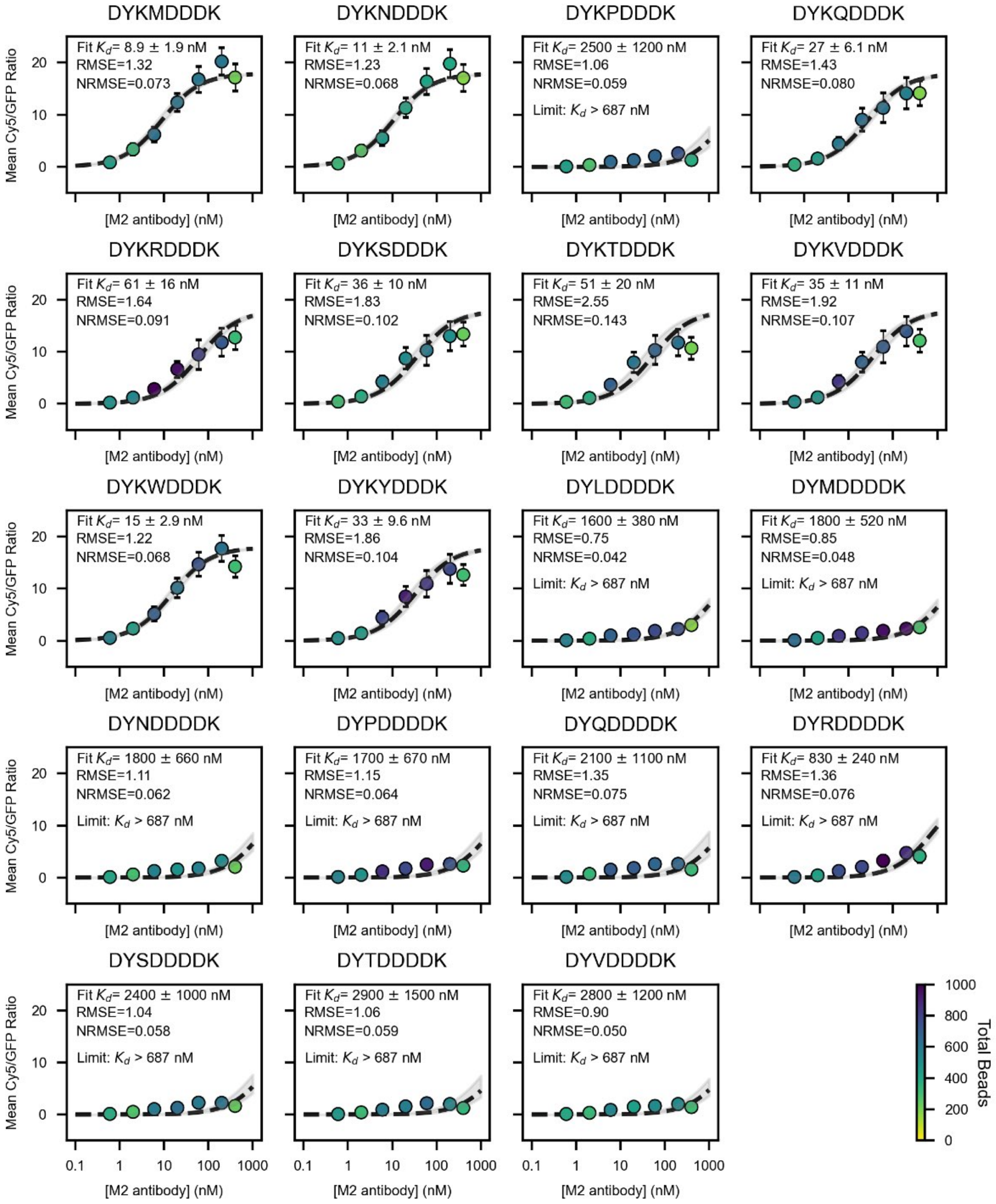


**Supplementary Figure 37 — Binding isotherms for single-variant library (page 7/8).**

Mean Cy5/GFP ratios (mean ± SD) as a function of M2 antibody concentration for all FLAG variants within the single-variant library. Gray fit curves (mean ± SD) show global fits to a Langmuir isotherm with a shared maximum saturation value (global $R_{max}$) and individual fitted $K_d$ value for each variant. Marker colors denote the number of sorted beads at each concentration; annotations report raw fitted $K_d$ values and whether measurements could be considered lower or upper limits.

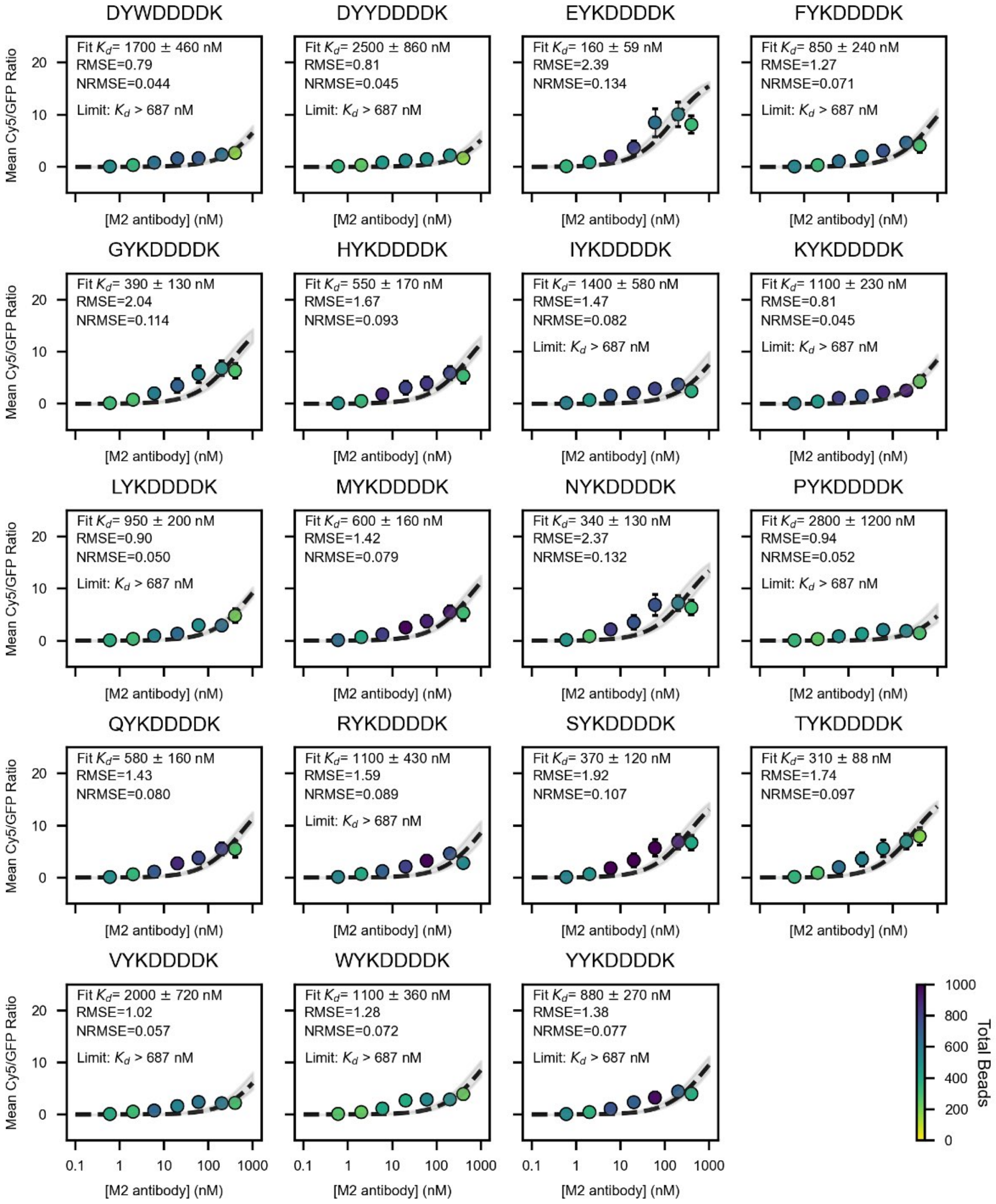


**Supplementary Figure 38 — Binding isotherms for single-variant library (page 8/8).**

Mean Cy5/GFP ratios (mean ± SD) as a function of M2 antibody concentration for all FLAG variants within the single-variant library. Gray fit curves (mean ± SD) show global fits to a Langmuir isotherm with a shared maximum saturation value (global $R_{max}$) and individual fitted $K_d$ value for each variant. Marker colors denote the number of sorted beads at each concentration; annotations report raw fitted $K_d$ values and whether measurements could be considered lower or upper limits.

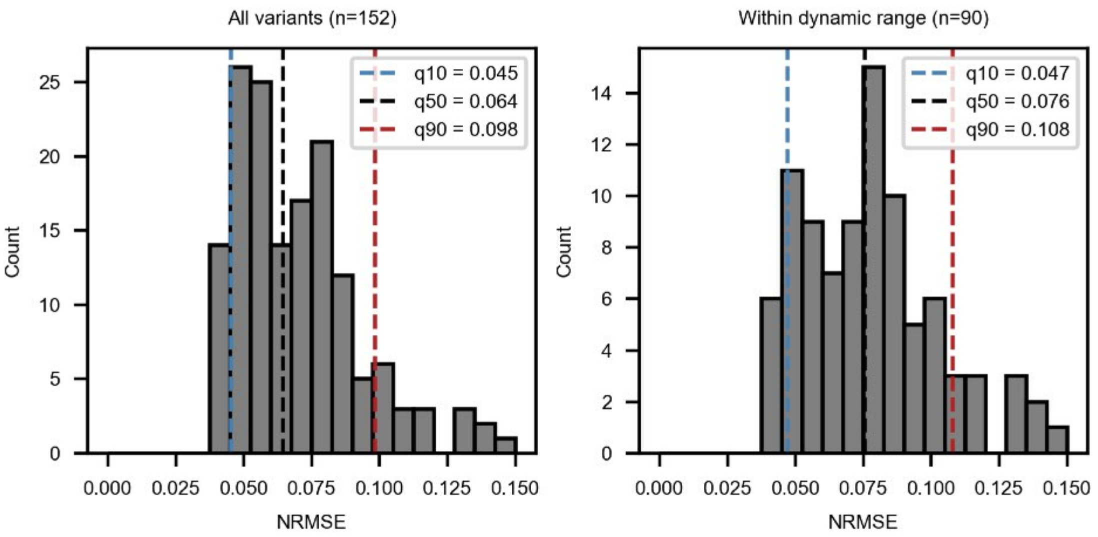


**Supplementary Figure 39 — Fit quality metrics for single-variant library.**

Distribution of normalized RMSE values (Cy5/GFP ratio RMSE / $R_{\max}$) for fitted $K_d$ values for all single-variants (including those with fitted $K_d$ above and below likely resolvable limits; left) and only those with $K_d$ within the simulated likely resolvable limits (right). Blue dotted line: 10th percentile; black dotted line: 50th percentile, red dotted line: 90th percentile.

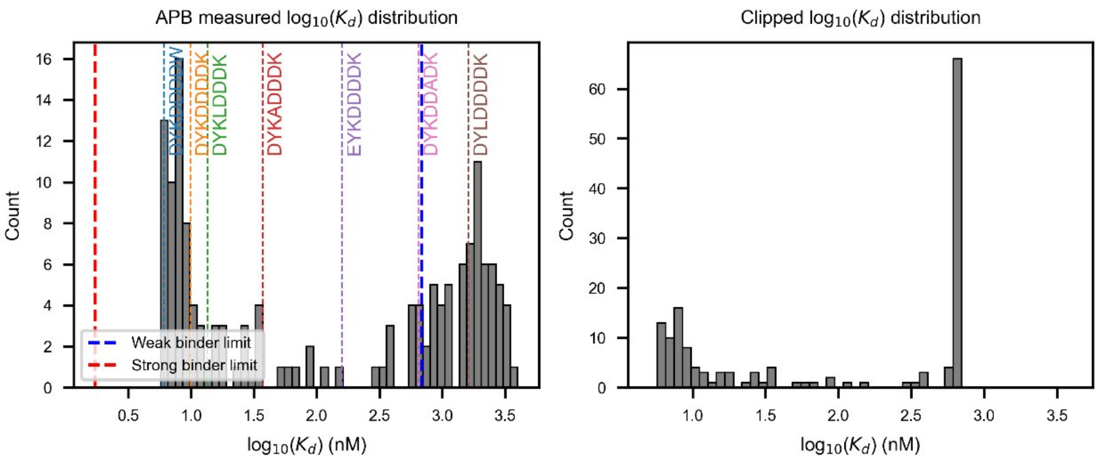


**Supplementary Figure 40 — Distributions of fitted $\log_{10}(K_d)$ values for the single-variant library.**

Distribution of fitted $\log_{10}(K_d)$ values for the single-variant librar without (left) and with (right) imposing lower and upper limits on resolvable affinities. Colored lines and text denote 'calibration' variants with orthogonal FP measurements. Blue dotted line: weak binder limit from simulation; red dotted line: strong binder limit from simulation.

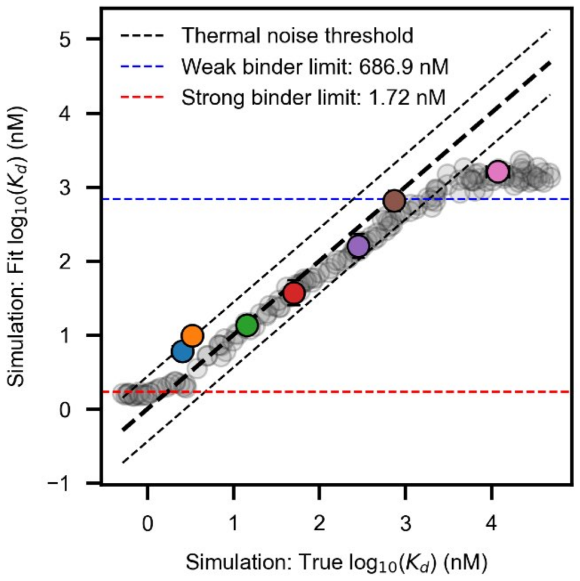


**Supplementary Figure 41 — Comparison of FP affinities and simulated affinity distribution for the single-variant library.**

Scatter plot comparing "fitted" $\log_{10}(K_d)$ values from APB-TiteSeq simulations versus "true" $\log_{10}(K_d)$ values used as inputs. Each point represents 1 simulation replicate. Black dashed lines denote thermal noise threshold (0.44 $\log_{10}(K_d)$ units, corresponding to 0.6 kcal/mol or 1 kT). Overlaid colored points show measured $\log_{10}(K_d)$ from APB-TiteSeq experiments versus (y-axis); FP-measured $\log_{10}(K_d)$ values for calibration variants (x-axis). Blue: DYKDDDDW, orange: DYKDDDDK, green: DYKLDDDK, red: DYKADDDK, purple: EYKDDDDK, brown: DYKDDADK, pink: DYLDDDDK. Blue and red dotted lines denote resolution limits for weak and strong binders, respectively, from simulations.

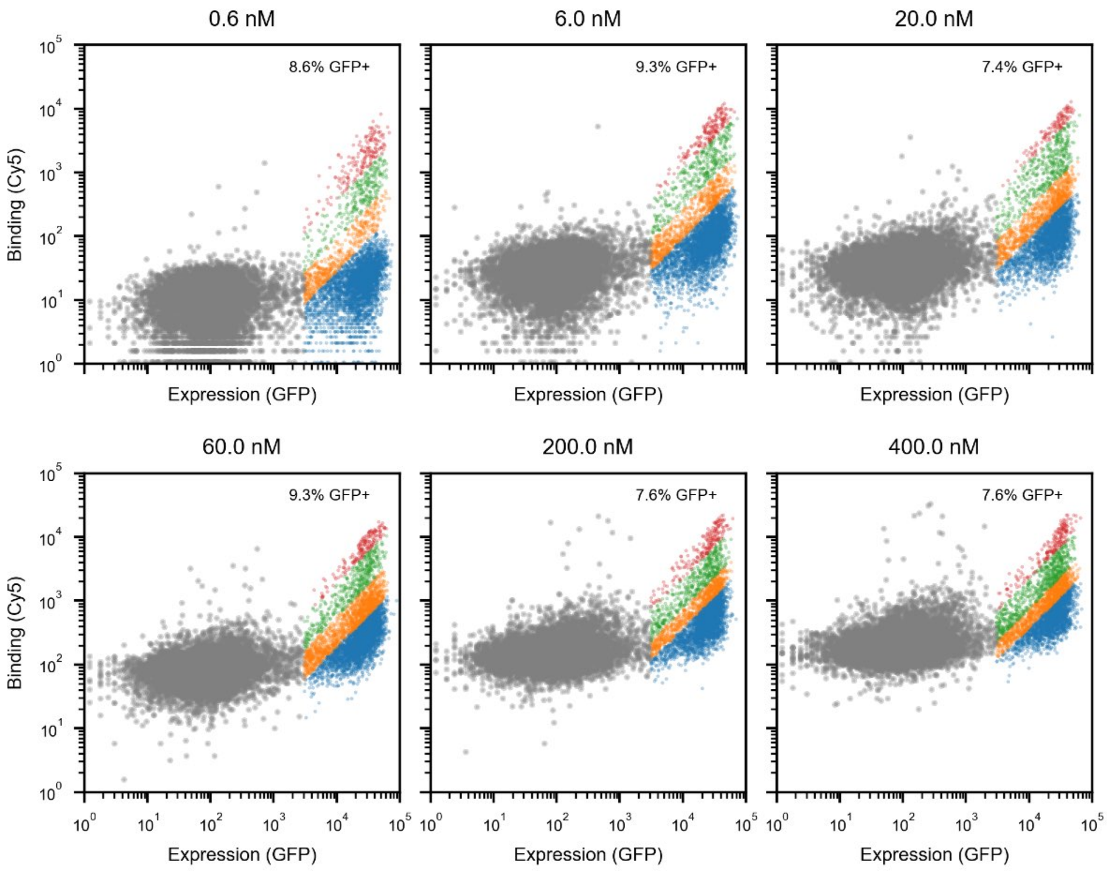


**Supplementary Figure 42 — Log-log FACS plots for the four-position library at each antibody concentration (replicate 1).**

Log-log FACS plot of Cy5 versus GFP intensities for recorded beads at each antibody concentration (four-position library, replicate 1). Gray: blank beads; blue: bin 1; orange: bin 2; green: bin 3; red: bin 4.

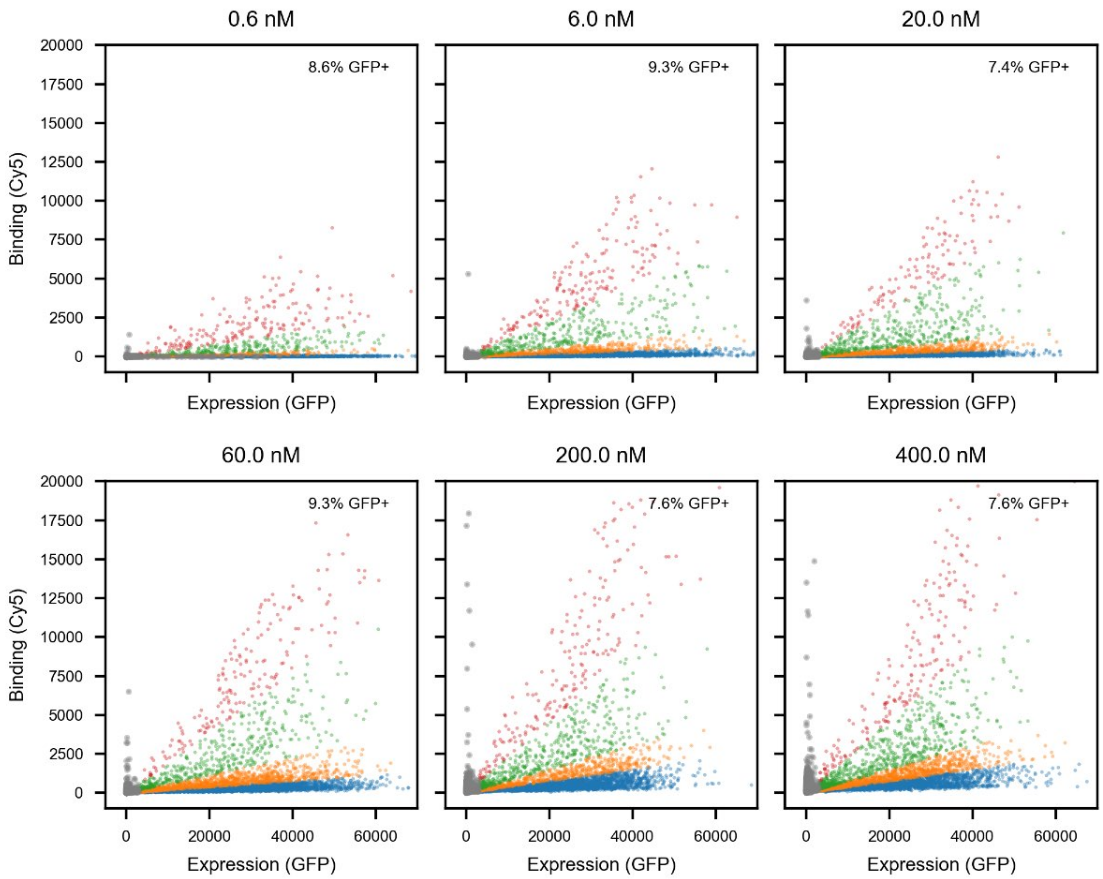


**Supplementary Figure 43 — Linear-linear FACS plots for the four-position library at each antibody concentration (replicate 1).**

Linear-linear FACS plot of Cy5 versus GFP intensities for recorded beads at each antibody concentration (four-position library, replicate 1). Gray: blank beads; blue: bin 1; orange: bin 2; green: bin 3; red: bin 4.

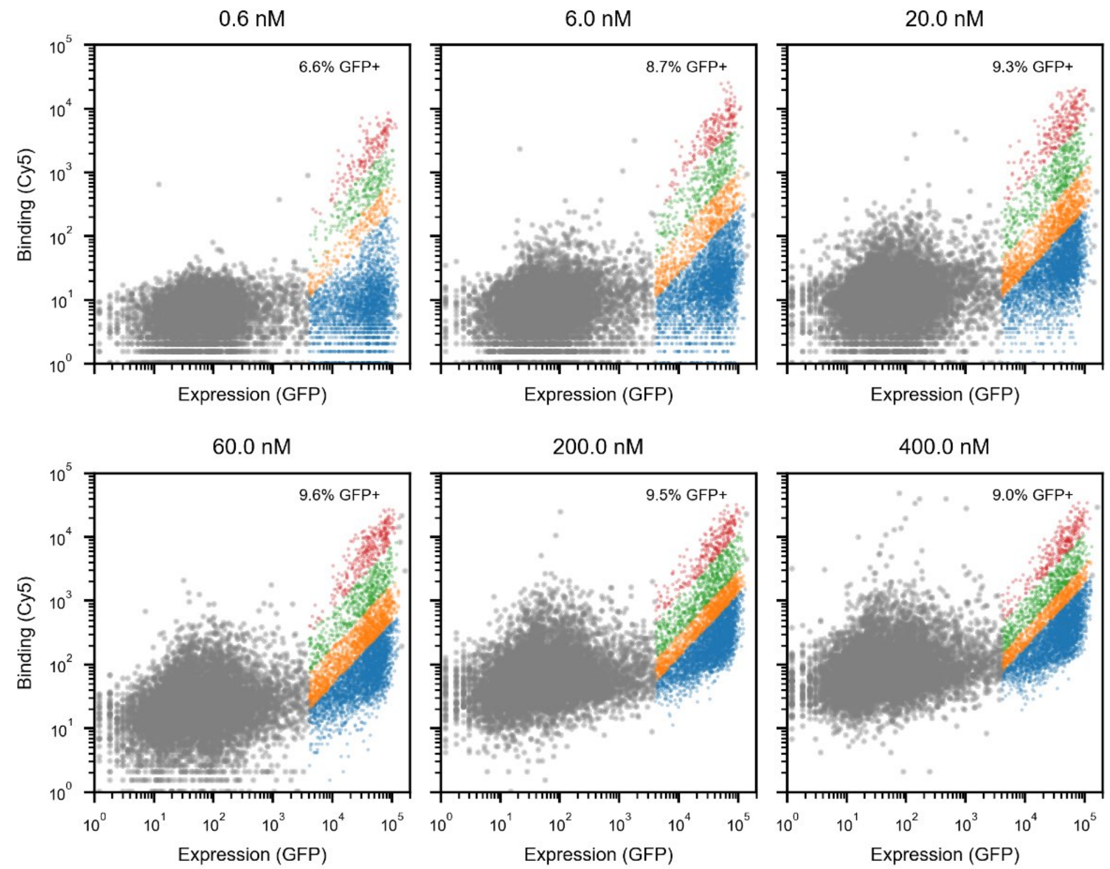


**Supplementary Figure 44 — Log-log FACS plots for the four-position library at each antibody concentration (replicate 2).**

Log-log FACS plot of Cy5 versus GFP intensities for recorded beads at each antibody concentration (four-position library, replicate 2). Gray: blank beads; blue: bin 1; orange: bin 2; green: bin 3; red: bin 4.

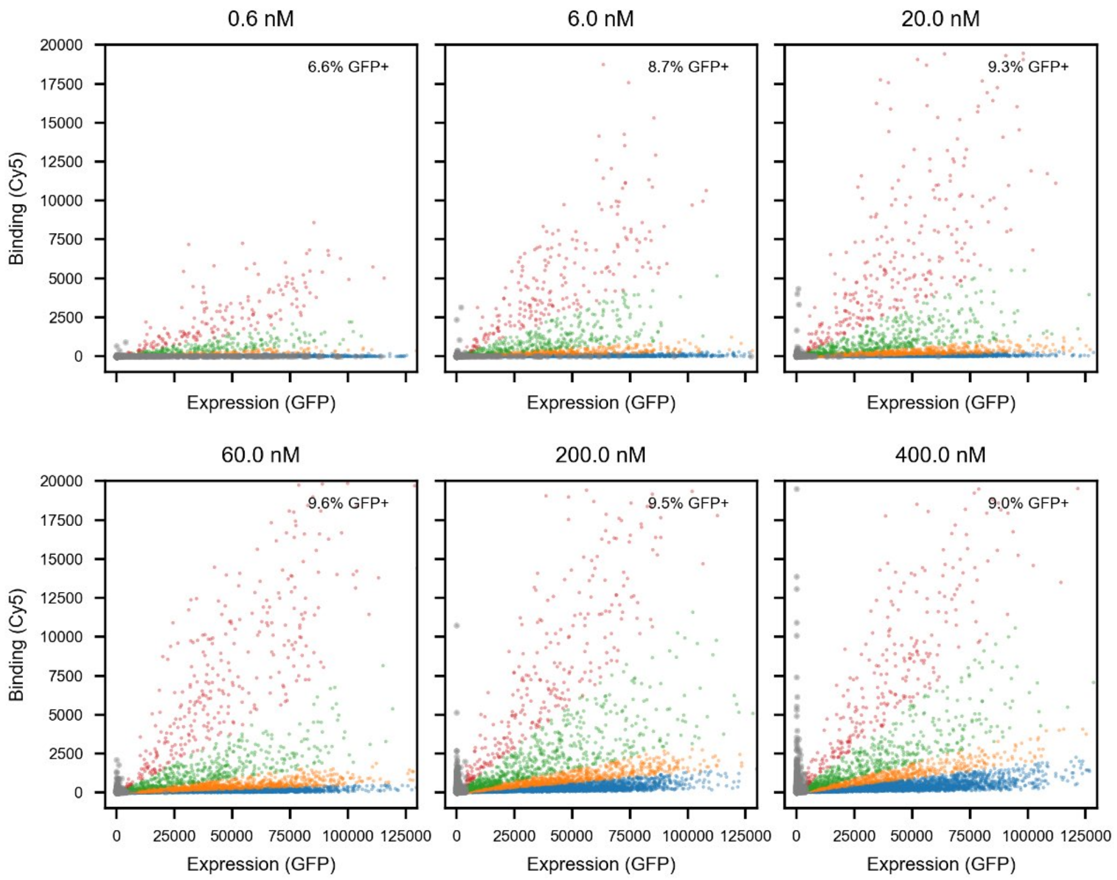


**Supplementary Figure 45 — Linear-linear FACS plots for the four-position library at each antibody concentration (replicate 2).**

Linear-linear FACS plot of Cy5 versus GFP intensities for recorded beads at each antibody concentration (four-position library, replicate 2). Gray: blank beads; blue: bin 1; orange: bin 2; green: bin 3; red: bin 4.

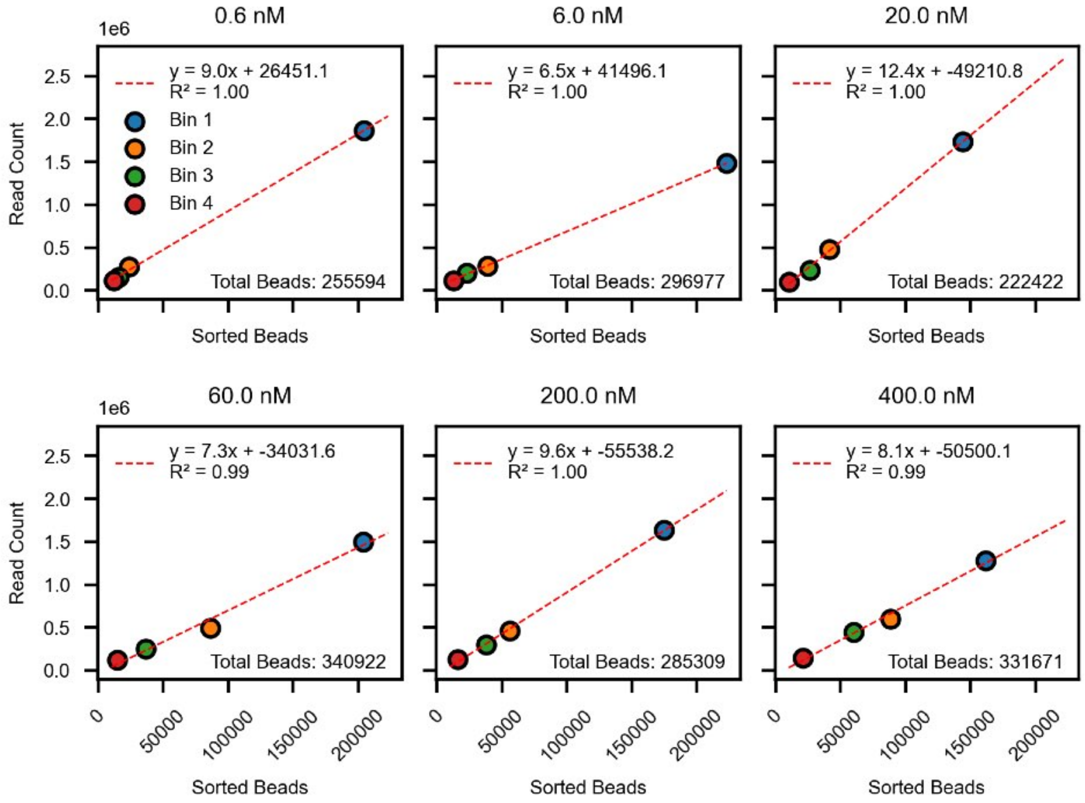


**Supplementary Figure 46 — Correlation between read counts and sorted bead counts for the four-position library (replicate 1).**

Read counts as a function of the number of sorted beads for each sort bin and concentration for the four-position library (replicate 1).

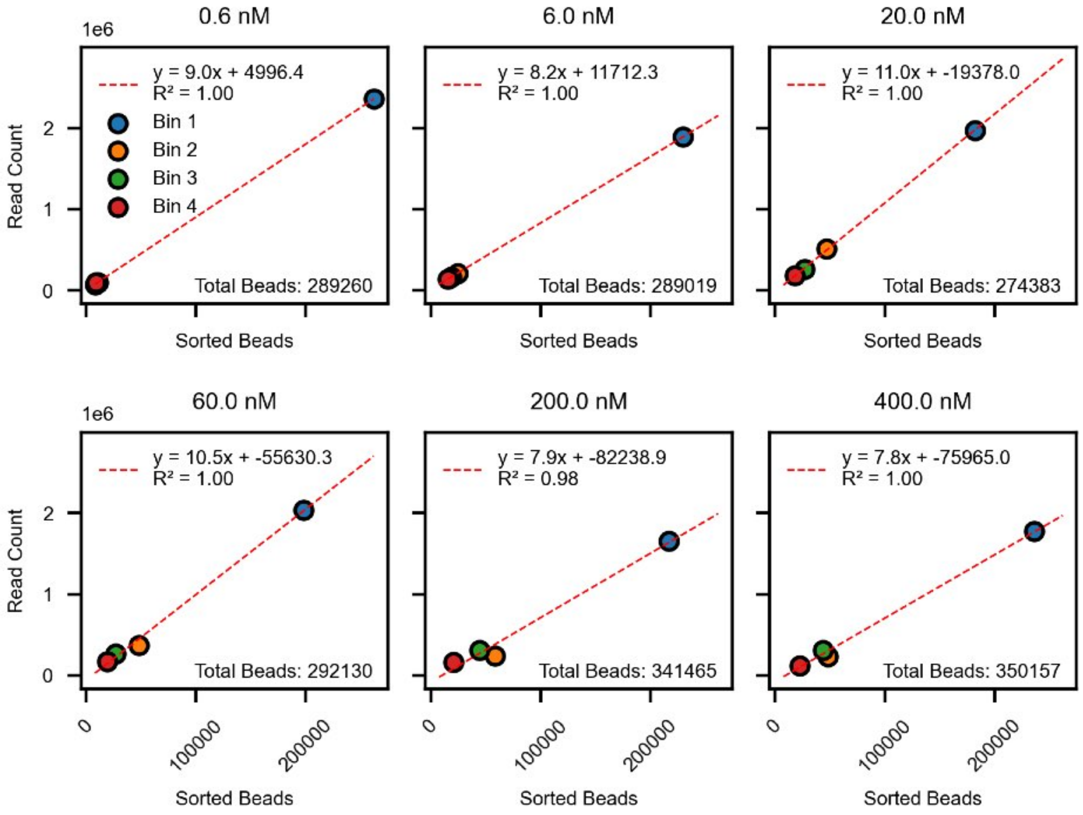


**Supplementary Figure 47 — Correlation between read counts and sorted bead counts for the four-position library (replicate 2).**

Read counts as a function of the number of sorted beads for each sort bin and concentration for the four-position library (replicate 2).

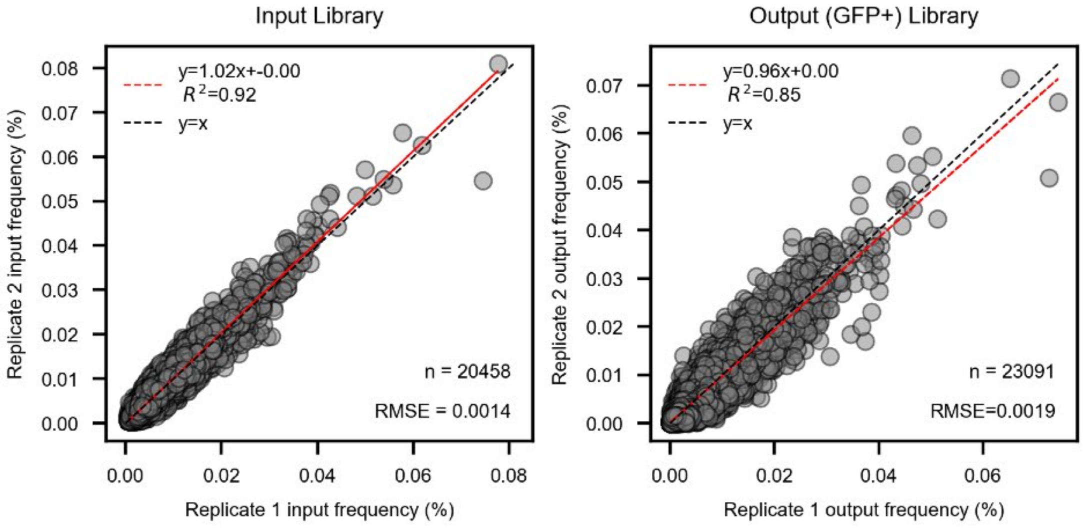


**Supplementary Figure 48 — Comparison of input and output library frequencies across replicates of the four-position library.**

Scatter plots comparing input (left) and output (right) frequencies for replicate 2 versus replicate 1. Each marker represents a single variant. Black dashed line: identity line; red dashed line: linear regression fit with annotated fit parameters.

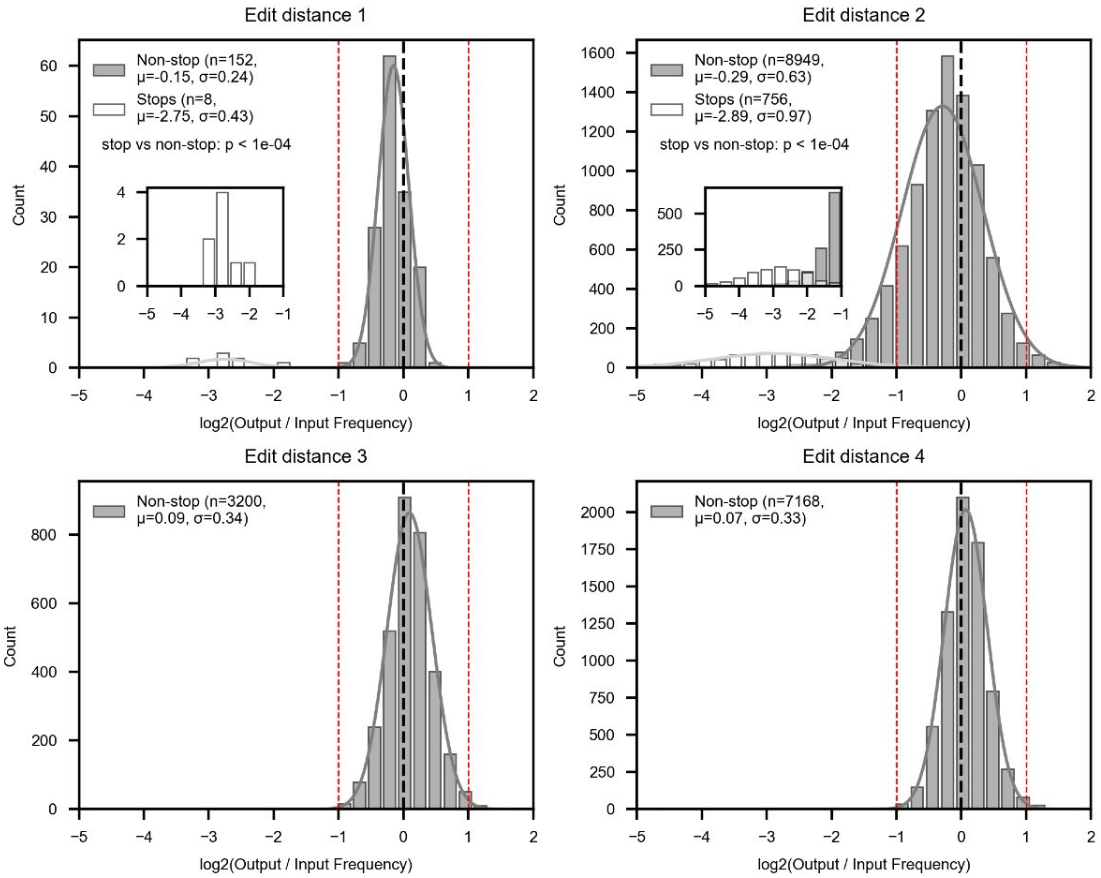


**Supplementary Figure 49 — Histograms of $\log_2$(output/input) frequency ratios as a function of edit distance for the four-position library.**

Histograms of $\log_2$(output/input) frequency ratios as a function of edit distance for the four-position library. Black dashed line indicates no frequency difference between input and output frequency; red dashed lines indicate 2-fold enrichment or depletion ($\log_2$(output/input)±1). Gray bars: variants without premature stop codons; white bars: variants containing premature stop codons. Gray and white lines represent Gaussian fits to each distribution, respectively (mean $\mu$ and standard deviation $\sigma$ reported in legend). Inset plots show zoomed-in views of depleted variant distributions ($\log_2$(output/input)<-1). p-values are from a two-sided permutation test on the difference of means between stop-codon and non-stop variant $\log_2$(output/input) ratios (n = 10,000 permutations). Values reported as $p < 10^{-4}$ indicate that no permutations produced a test statistic as extreme as the observed difference, giving an upper bound of 1/10,000.

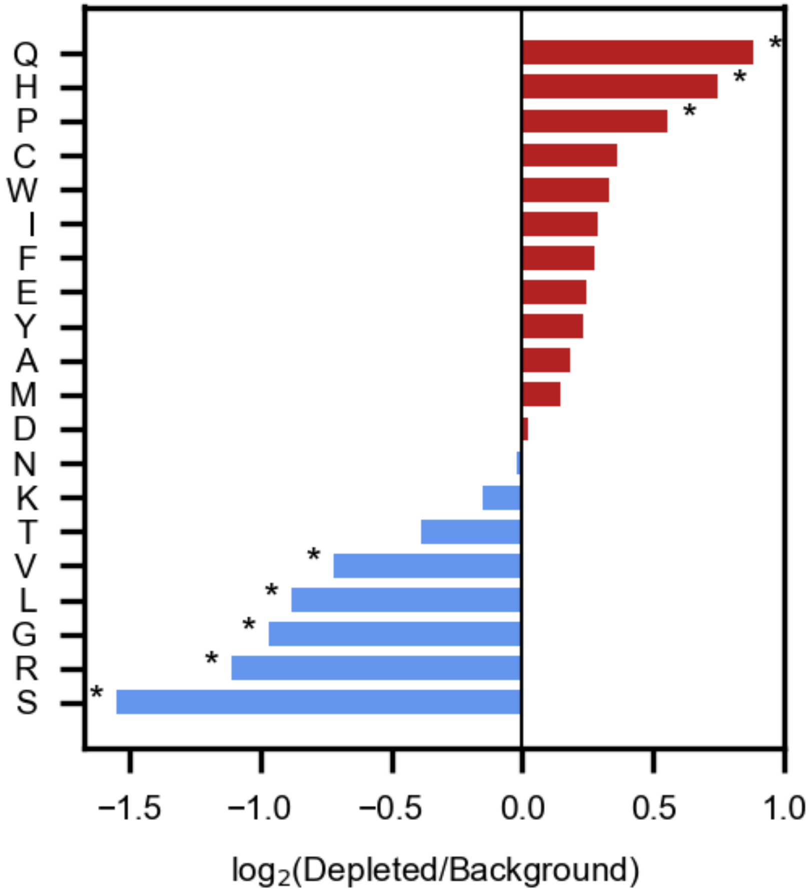


**Supplementary Figure 50 — Amino acid enrichment in depleted double mutant FLAG variants from the four-position library.**

Bar plot showing enrichment of specific residues within the >2-fold depleted double mutant population (calculated as $log_2$(frequency in depleted variants / frequency in all double mutants), where frequency reflects the fraction of mutation events introducing each residue at each position). Asterisks indicate amino acids with statistically significant enrichment or depletion (Fisher's exact test, Bonferroni correction, $p < 0.05$).

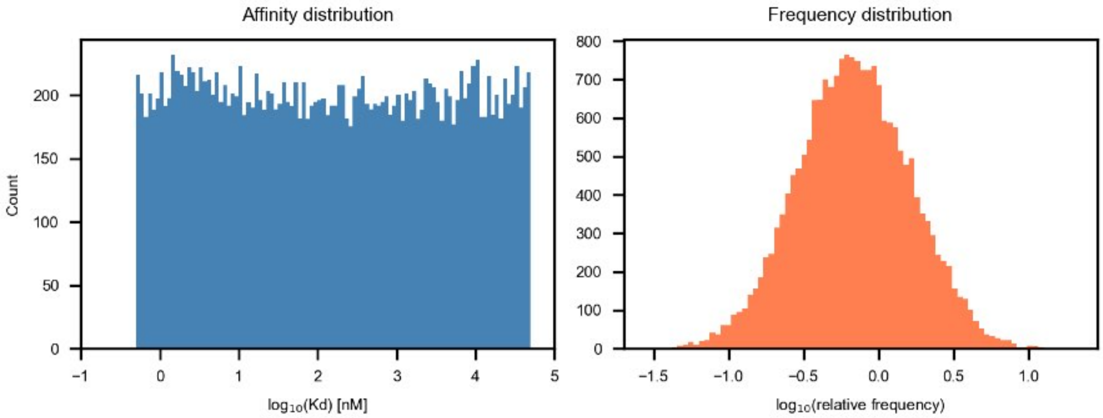


**Supplementary Figure 51 — Histograms of affinity and frequency distributions for simulating the four-position library.**

Left: Histogram of 20,000 log-normally-distributed affinities between 0.5-50,000 nM used for simulating the four-position library. Right: Histogram of variant frequencies in the simulated library (10.2-fold skew between 10th and 90th most abundant variants).

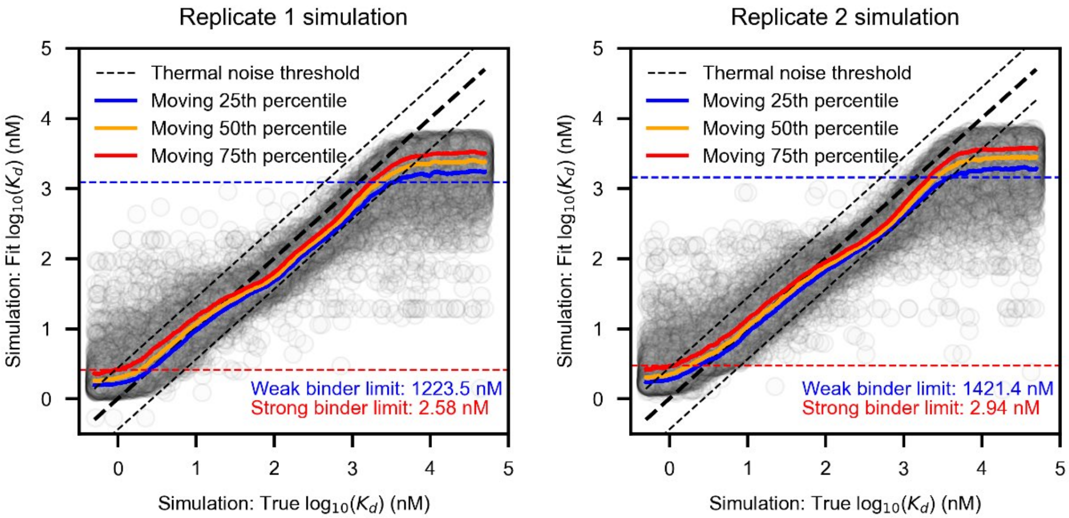


**Supplementary Figure 52 — Simulation method for determining dynamic range limits for the four-position library.**

Scatter plot comparing "fitted" $\log_{10}(K_d)$ values from APB-TiteSeq simulations for the four-position library versus "true" $\log_{10}(K_d)$ values used as inputs for replicate 1 (left) and replicate 2 (right). Each point represents 1 simulation replicate and data shown include 3 simulation replicates. Black dashed lines denote thermal noise threshold (0.44 $\log_{10}(K_d)$ units, corresponding to 0.6 kcal/mol or 1 kT); colored lines denote moving percentiles of fitted $\log_{10}(K_d)$ values. Blue and red dotted lines denote resolution limits for weak and strong binders (where the 25th and 75th percentile lines cross the thermal noise threshold, respectively).

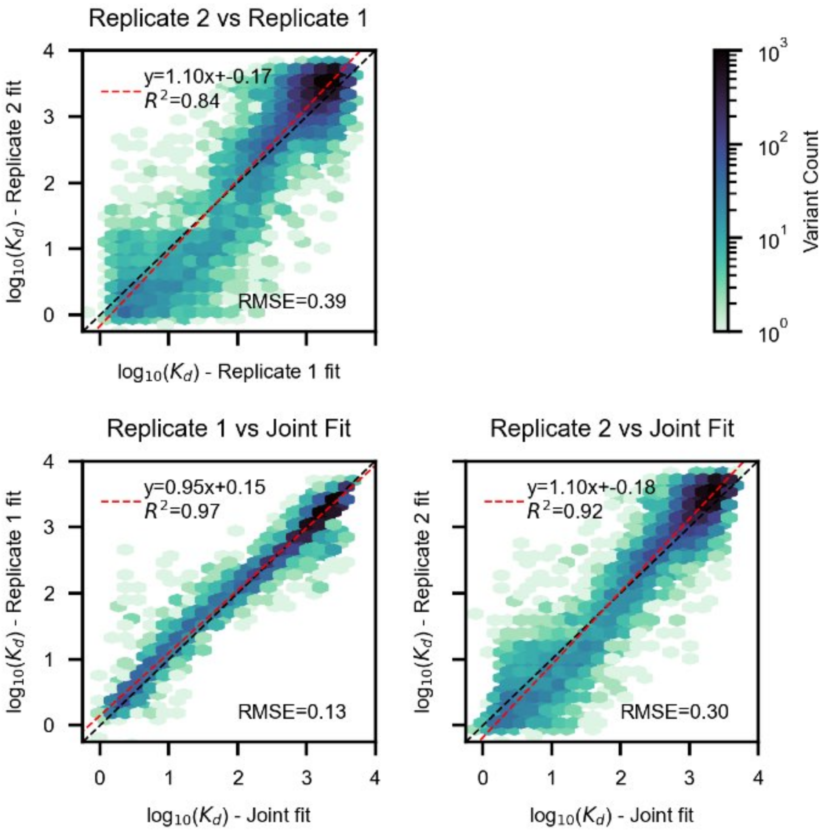


**Supplementary Figure 53 — Comparison between individual replicate fits and joint fits for the four-position library.**

Top: Hexbin plot of returned fitted $\log_{10}(K_d)$ values from replicate 2 versus replicate 1 (each analyzed separately). Bottom: Hexbin plots of returned fitted $\log_{10}(K_d)$ values from replicate 1 (left) and replicate 2 (right) versus returned values from fitting both datasets concurrently. Hexbins are colored by the number of variants with a given affinity range (log scale). Black dashed line indicates 1:1 line; red dashed line indicates linear regression with annotated fit parameters.

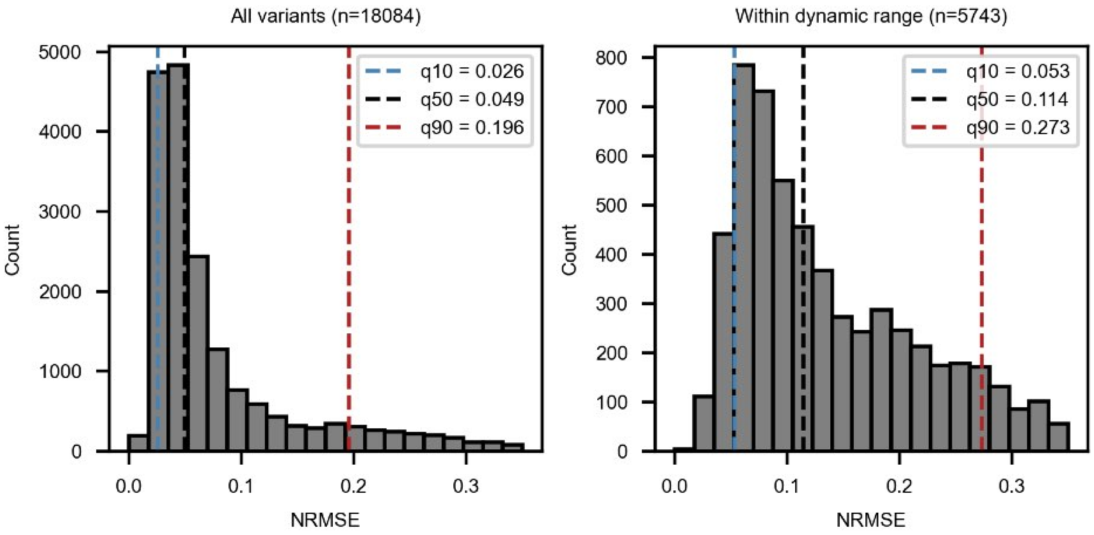


**Supplementary Figure 54 — Fit quality metrics for four-position library.**

Distribution of normalized RMSE values (Cy5/GFP ratio RMSE / $R_{\max}$) for fitted $K_d$ values for all variants in the four-position library (left) and those with $K_d$ values within the simulated dynamic range boundaries (right). Blue dotted line: 10th percentile; black dotted line: 50th percentile, red dotted line: 90th percentile.

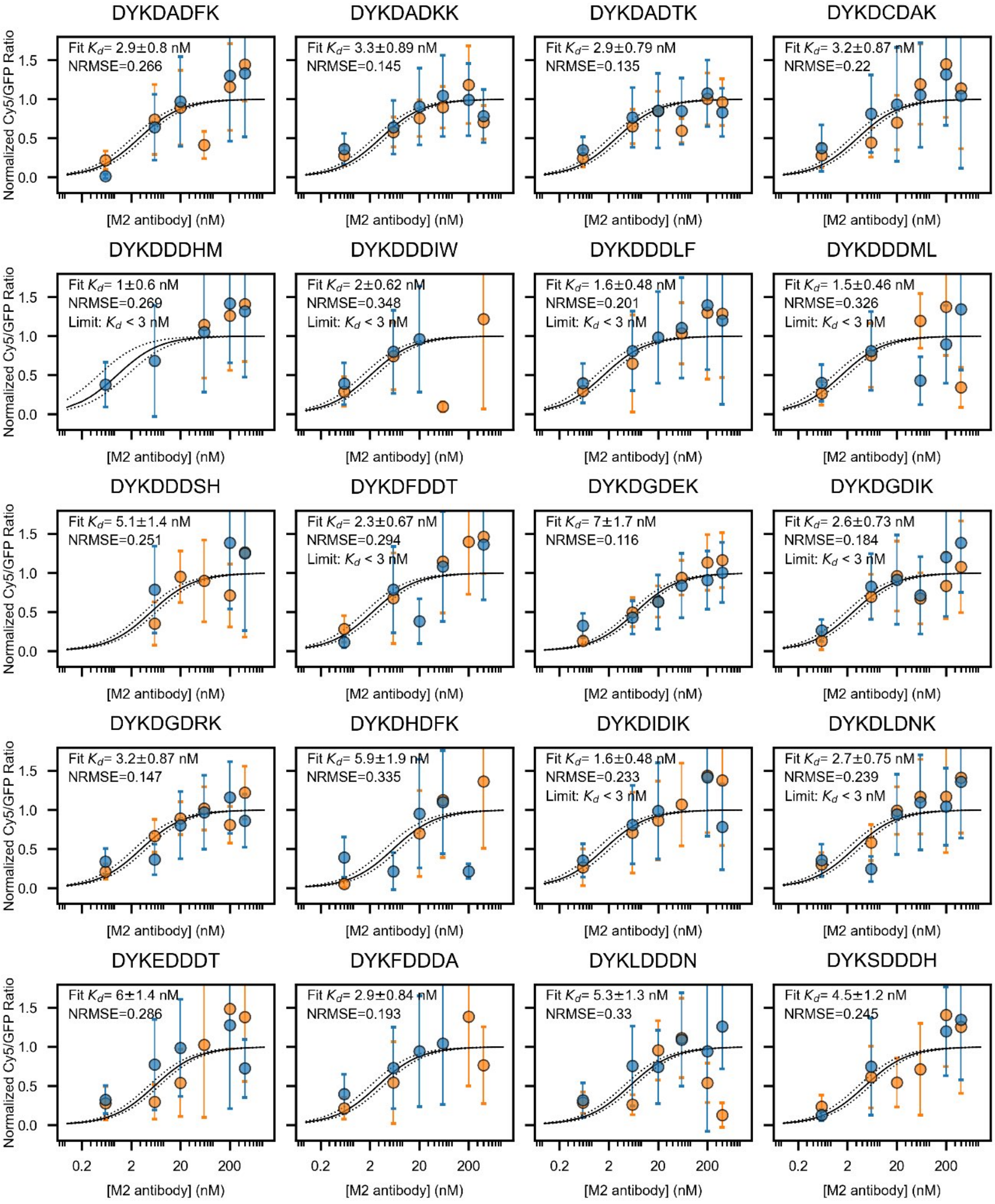


**Supplementary Figure 55 — Binding isotherms for randomly-sampled variants with affinities <10 nM in the four-position library.**

Normalized Cy5/GFP ratios as a function of M2 antibody concentration and Langmuir isotherm fits for randomly-sampled variants with estimated affinities <10 nM. Markers denote mean Cy5/GFP ratios (mean ± SD) normalized to experiment-specific $R_{\max}$; error bars denote standard deviation. Orange markers: experimental replicate 1; blue markers: experimental replicate 2. Black curves (±SD) are jointly fit to both experimental replicates. Fitted $K_d$ values outside the upper and lower limits established from simulation are also reported as limits.

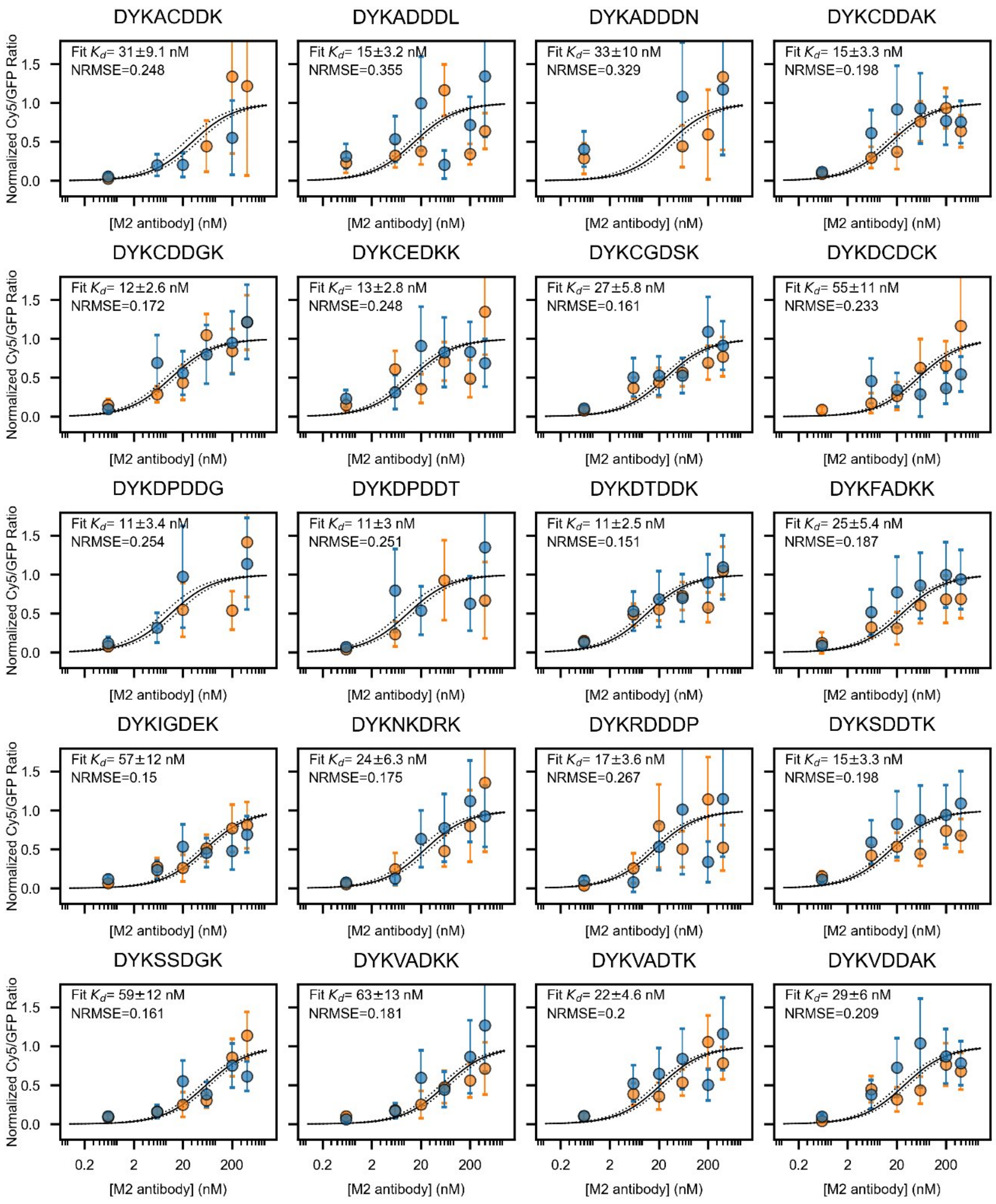


**Supplementary Figure 56 — Binding isotherms for randomly-sampled variants with affinities between 10 nM and 100 nM in the four-position library.**

Normalized Cy5/GFP ratios as a function of M2 antibody concentration and Langmuir isotherm fits for randomly-sampled variants with estimated affinities >10 nM and <100 nM. Markers denote mean Cy5/GFP ratios (mean ± SD) normalized to experiment-specific $R_{\max}$; error bars denote standard deviation. Orange markers: experimental replicate 1; blue markers: experimental replicate 2. Black curves (±SD) are jointly fit to both experimental replicates. Fitted $K_d$ values outside the upper and lower limits established from simulation are also reported as limits.

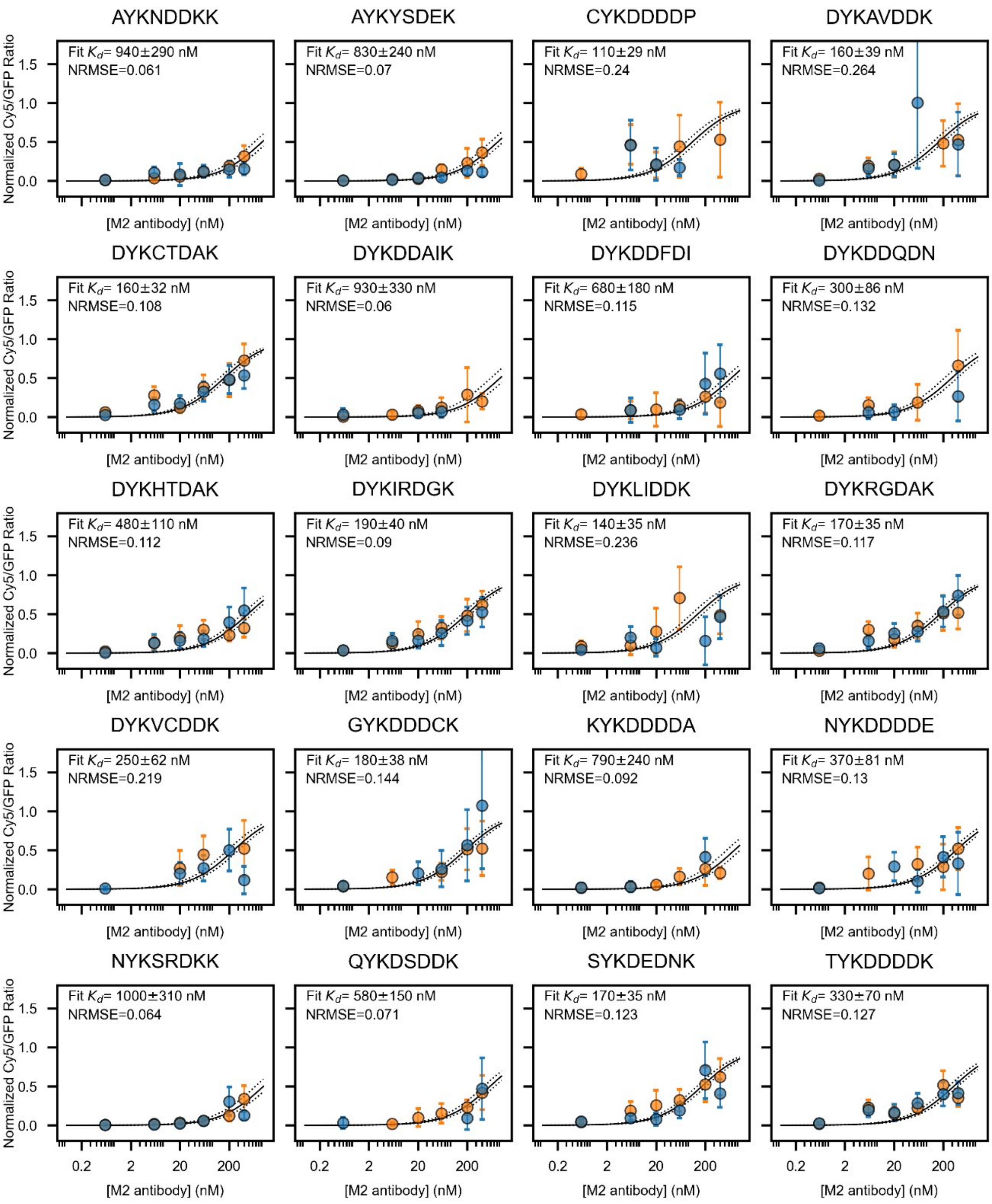


**Supplementary Figure 57 — Binding isotherms for randomly-sampled variants with affinities between 100 nM and 1000 nM in the four-position library.**

Normalized Cy5/GFP ratios as a function of M2 antibody concentration and Langmuir isotherm fits for randomly-sampled variants with estimated affinities >100 nM and <1000 nM. Markers denote mean Cy5/GFP ratios (mean ± SD) normalized to experiment-specific $R_{\max}$; error bars denote standard deviation. Orange markers: experimental replicate 1; blue markers: experimental replicate 2. Black curves (±SD) are jointly fit to both experimental replicates. Fitted $K_d$ values outside the upper and lower limits established from simulation are also reported as limits.

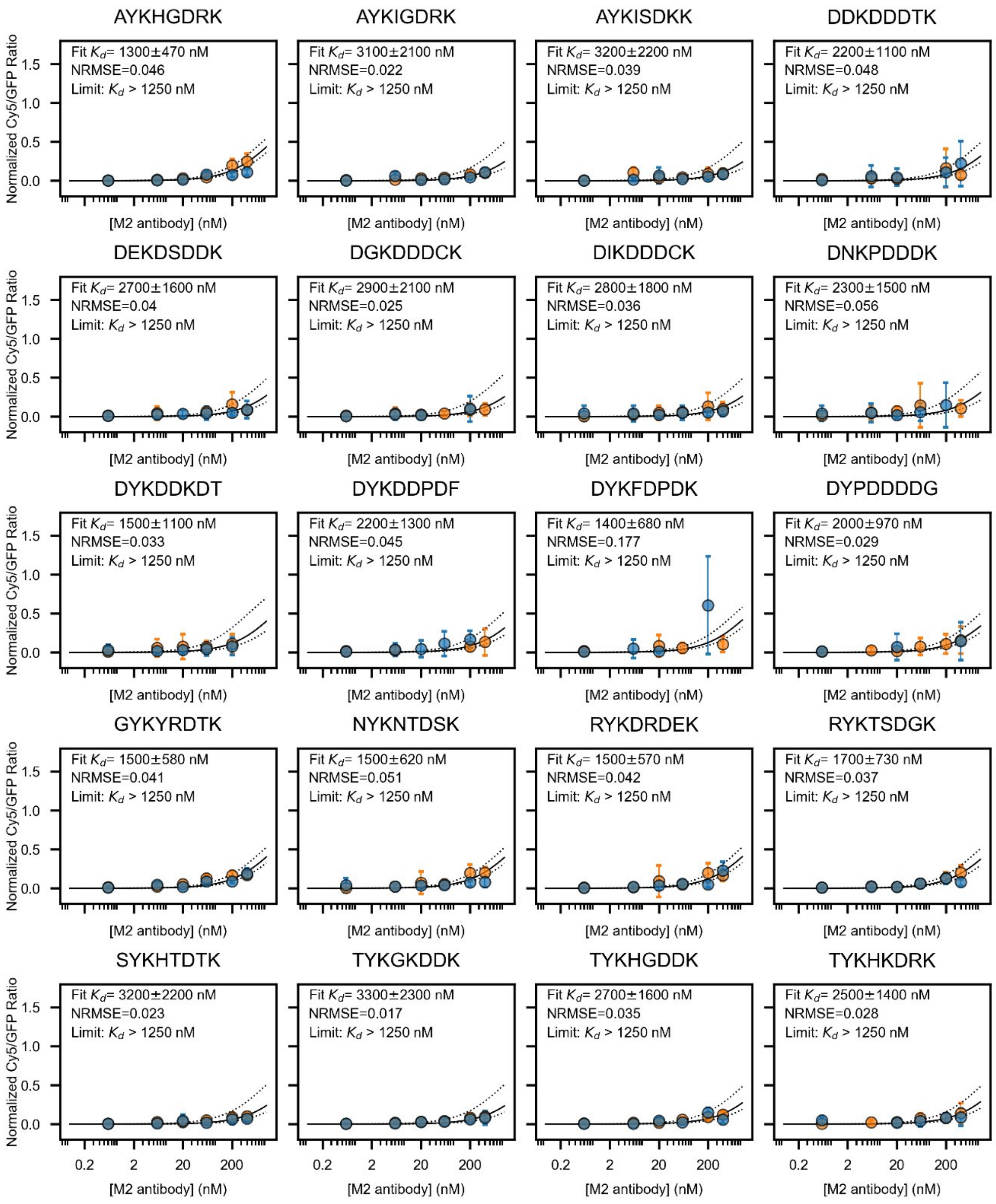


**Supplementary Figure 58 — Binding isotherms for randomly-sampled variants with affinities >1000 nM in the four-position library.**

Normalized Cy5/GFP ratios as a function of M2 antibody concentration and Langmuir isotherm fits for randomly-sampled variants with estimated affinities >1000 nM. Markers denote mean Cy5/GFP ratios (mean ± SD) normalized to experiment-specific $R_{\text{max}}$; error bars denote standard deviation. Orange markers: experimental replicate 1; blue markers: experimental replicate 2. Black curves (±SD) are jointly fit to both experimental replicates. Fitted $K_d$ values outside the upper and lower limits established from simulation are also reported as limits.

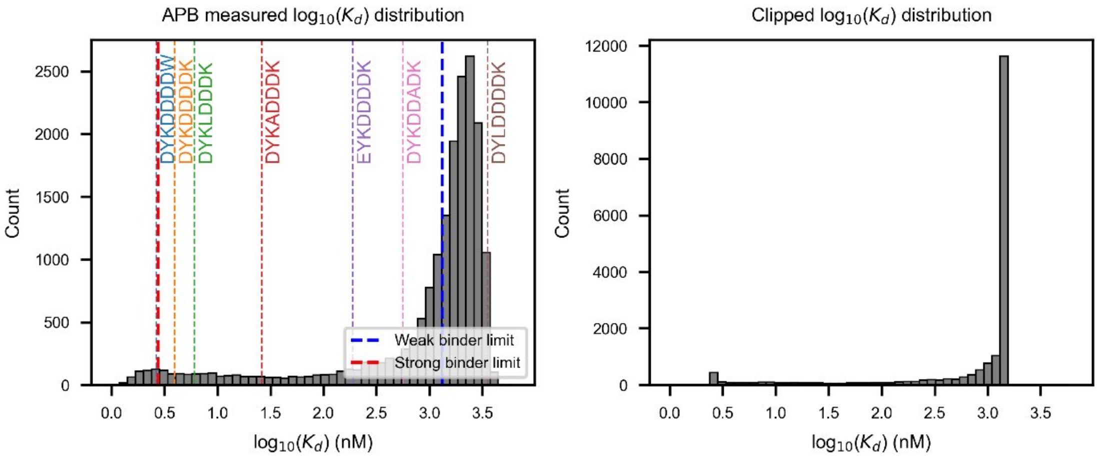


**Supplementary Figure 59 — Distribution of returned fitted $\log_{10}(K_d)$ values for the four-position library.** Distribution of all returned fitted $\log_{10}(K_d)$ values (left) and fitted $\log_{10}(K_d)$ values after imposing upper and lower resolution limits (right) for the four-position library. Colored lines and text: calibration variants. Blue dotted line: average weak binder limit from simulation; red dotted line: average strong binder limit from simulation.

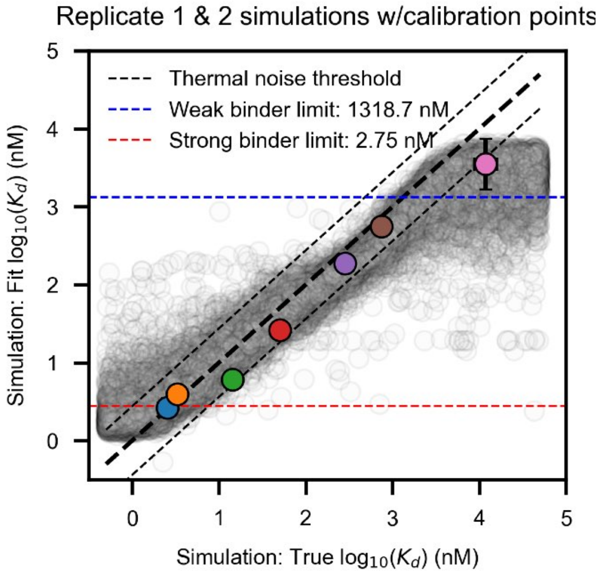


**Supplementary Figure 60 — Comparison of FP-measured affinities and simulated affinity distribution for the four-position library.**

Scatter plot of returned "fitted" $\log_{10}(K_d)$ values from the APB-TiteSeq simulation versus "true" $\log_{10}(K_d)$ values used as input. Each point represents a single simulation replicate based on parameters from the replicate 1 experiment and a single simulation replicate based on parameters from the replicate 2 experiment; the graph shows results from 5 simulations. Black dashed lines denote thermal noise threshold (0.44 $\log_{10}(K_d)$ units, corresponding to 0.6 kcal/mol or 1 kT). Overlaid colored points show measured $\log_{10}(K_d)$ from APB-TiteSeq versus FP-measured $\log_{10}(K_d)$ values for calibration variants (x-axis). Blue dotted line: average weak binder limit from simulation; red dotted line: average strong binder limit from simulation.

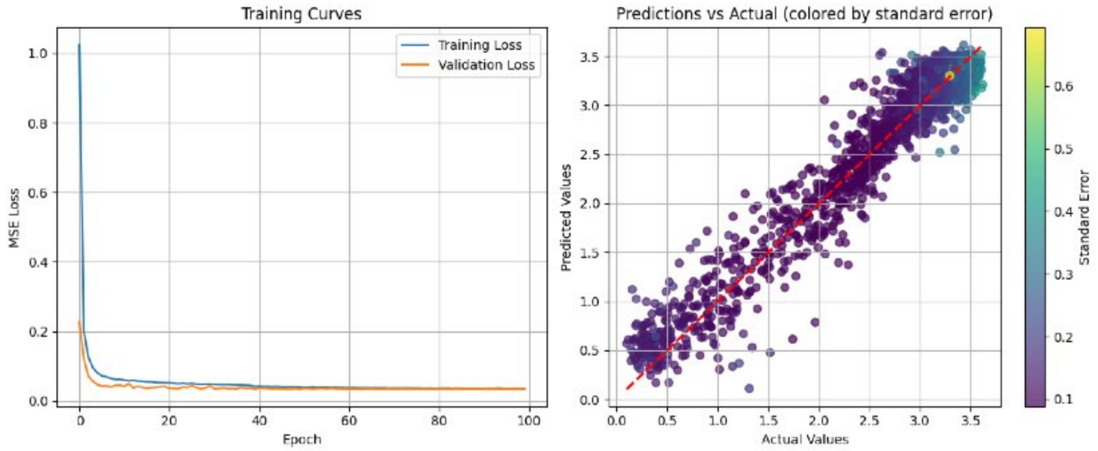


**Supplementary Figure 61 — Performance of four-position library CNN during Fold 1 of 5-fold cross-validation.**

Left: Training (blue) and validation (orange) MSE loss as a function of training epoch. Right: Predicted versus actual (experimentally-measured) $\log_{10}(K_d)$ values in nM, colored by standard error. Performance metrics are reported in Supplementary Table 4. Red dashed line indicates the identity line.

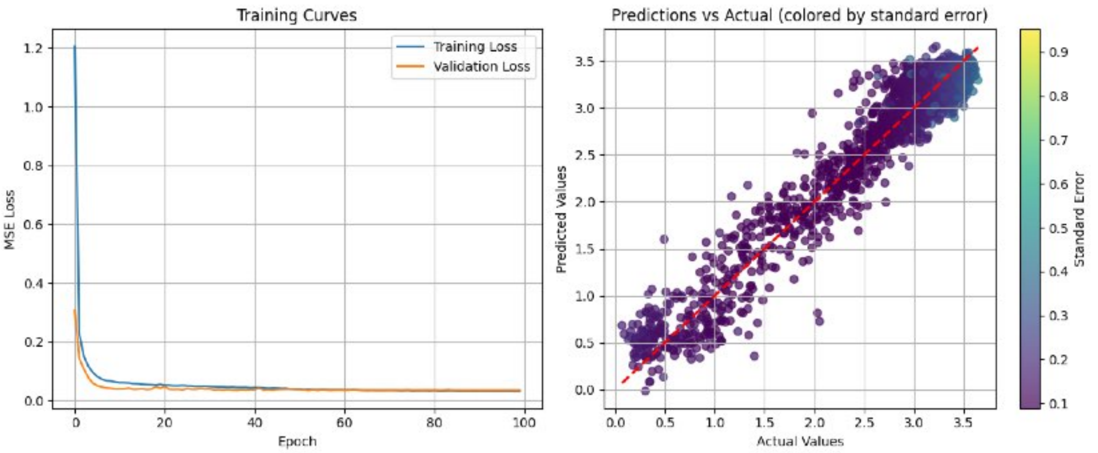


**Supplementary Figure 62 — Performance of four-position library CNN during Fold 2 of 5-fold cross-validation.** Left: Training (blue) and validation (orange) MSE loss as a function of training epoch. Right: Predicted versus actual (experimentally-measured) $\log_{10}(K_d)$ values in nM, colored by standard error. Performance metrics are reported in Supplementary Table 4. Red dashed line indicates the identity line.

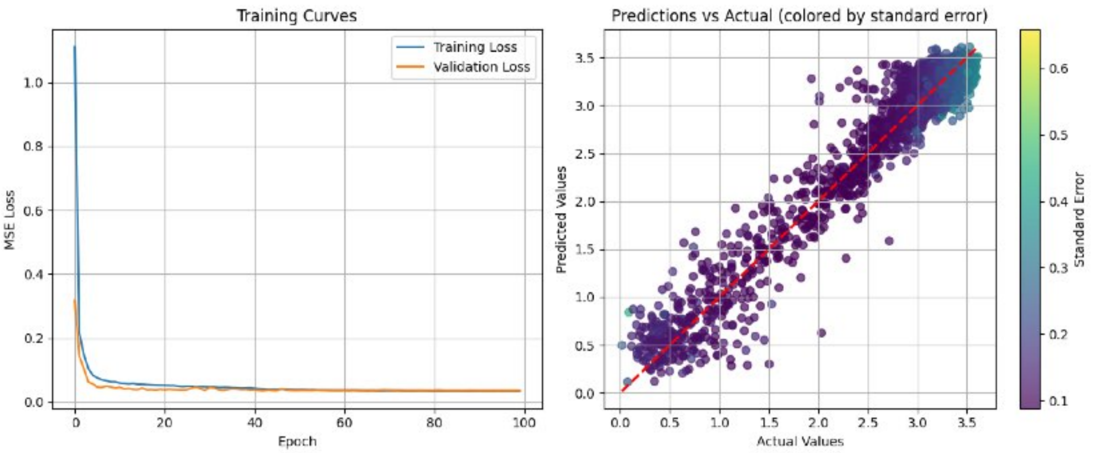


**Supplementary Figure 63 — Performance of four-position library CNN during Fold 3 of 5-fold cross-validation.** Left: Training (blue) and validation (orange) MSE loss as a function of training epoch. Right: Predicted versus actual (experimentally-measured) $\log_{10}(K_d)$ values in nM, colored by standard error. Performance metrics are reported in Supplementary Table 4. Red dashed line indicates the identity line.

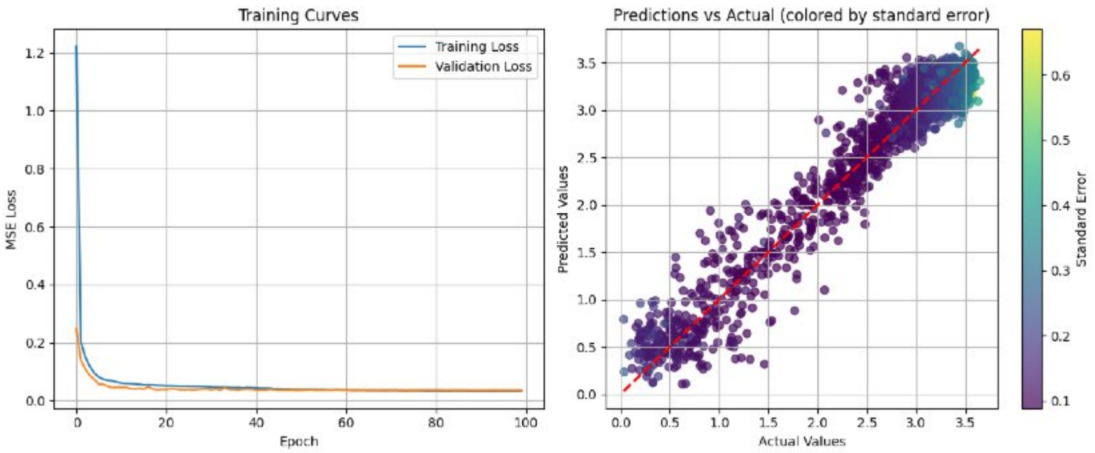


**Supplementary Figure 64 — Performance of four-position library CNN during Fold 4 of 5-fold cross-validation.** Left: Training (blue) and validation (orange) MSE loss as a function of training epoch. Right: Predicted versus actual (experimentally-measured) $\log_{10}(K_d)$ values in nM, colored by standard error. Performance metrics are reported in Supplementary Table 4. Red dashed line indicates the identity line.

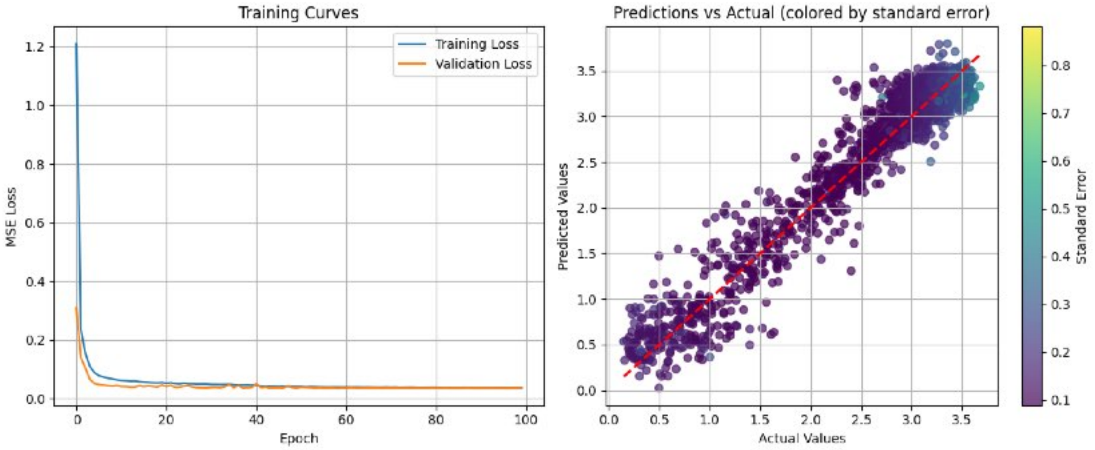


**Supplementary Figure 65 — Performance of four-position library CNN during Fold 5 of 5-fold cross-validation.** Left: Training (blue) and validation (orange) MSE loss as a function of training epoch. Right: Predicted versus actual (experimentally-measured) $\log_{10}(K_d)$ values in nM, colored by standard error. Performance metrics are reported in Supplementary Table 4. Red dashed line indicates the identity line.

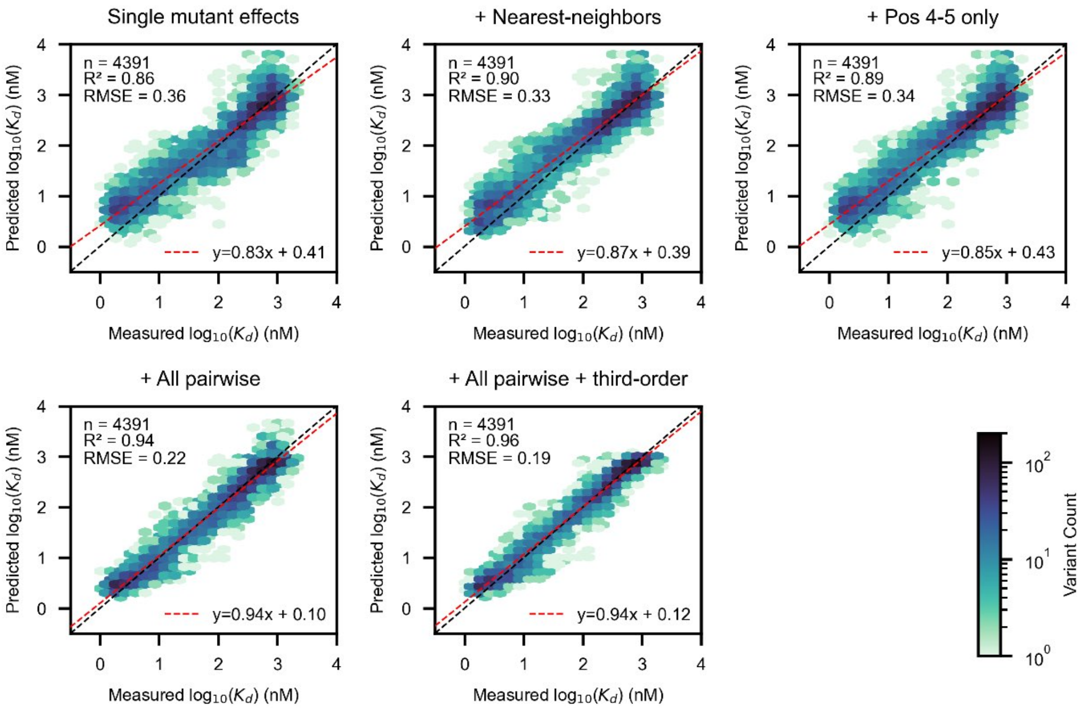


**Supplementary Figure 66 — Correlation between predicted and measured $\log_{10}(K_d)$ from linear models of increasing complexity derived from CNN predictions.**

Hexbin plots showing predicted versus measured $\log_{10}(K_d)$ for linear models including increasing numbers of terms constructed from CNN predictions. Predictions are made using all measured variants with $\log_{10}(K_d) < 3$. Hexbins are colored by the number of variants with a given affinity range (log scale).

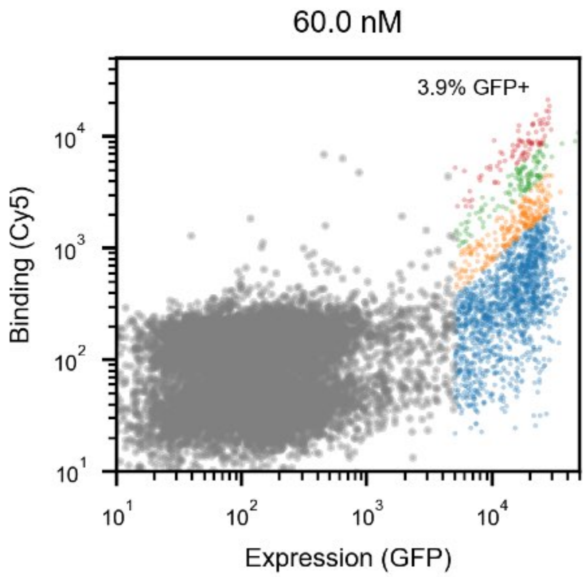


**Supplementary Figure 67 — Log-log FACS plot for five-position library at a single antibody concentration.**

Log-log FACS plot of Cy5 versus GFP intensities for recorded beads at 60 nM antibody concentration (five-position library). Gray: blank beads; blue: bin 1; orange: bin 2; green: bin 3; red: bin 4.

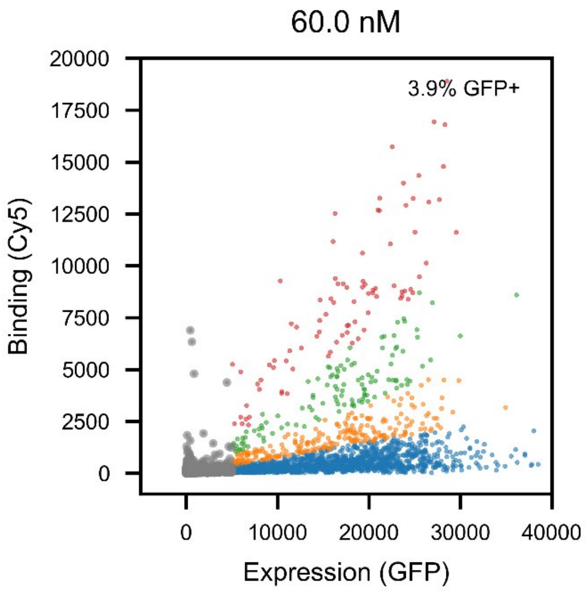


**Supplementary Figure 68 — Linear-linear FACS plot for five-position library at a single antibody concentration.**

Linear-linear FACS plot of Cy5 versus GFP intensities for recorded beads at 60 nM antibody concentration (five-position library). Gray: blank beads; blue: bin 1; orange: bin 2; green: bin 3; red: bin 4.

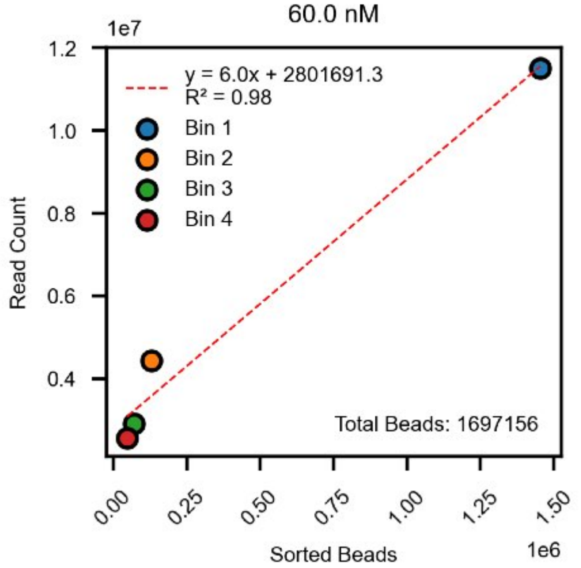


**Supplementary Figure 69 — Correlation between read counts and sorted bead counts for the five-position library.**

Read counts versus the number of sorted beads for each sort bin for the five-position library.

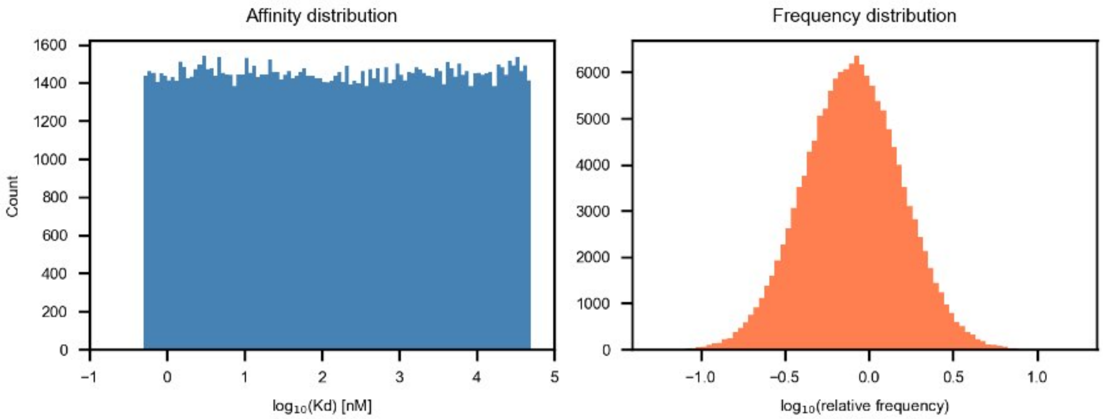


**Supplementary Figure 70 — Histograms of affinity and frequency distributions for simulating the five-position library.**

Left: Histogram of 145,000 log-normally-distributed affinities between 0.5-50,000 nM used for simulating the four-position library. Right: Histogram of variant frequencies in the simulated library (5.6-fold skew between 10th and 90th most abundant variants).

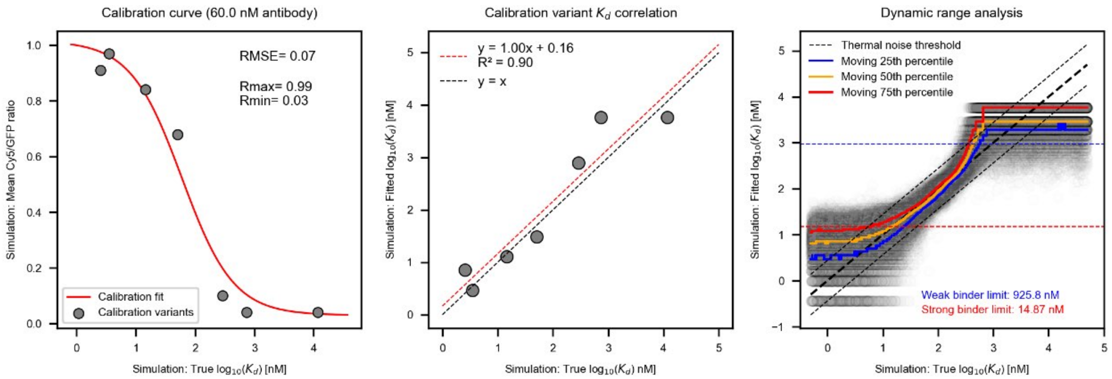


**Supplementary Figure 71 — APB-SortSeq simulation pipeline for determining upper and lower limits of resolution for the five-position library.**

(Left) Binding isotherm fit (red line) to seven simulated calibration variants with fluorescence polarization (FP)-derived 'true' $\log_{10}(K_d)$ values, used to extract global assay parameters $R_{\text{max}}$ and $R_{\text{min}}$ from the mean Cy5/GFP fluorescence ratio at 60 nM ligand. (Center) Correlation between simulation-fitted $\log_{10}(K_d)$ and true FP-derived $\log_{10}(K_d)$ for the seven calibration variants. Black dashed line indicates 1:1 agreement; red dashed line indicates unweighted linear regression. (Right) Dynamic range analysis across the full simulated library. Gray points show individual variant affinities; colored lines show moving 25th, 50th, and 75th percentiles of fitted $\log_{10}(K_d)$ as a function of true $\log_{10}(K_d)$. The black dashed thermal noise threshold (0.44 $\log_{10}$ units, equivalent to 0.6 kcal/mol ΔΔ G) defines the precision limit; the assay reliably resolves variants in the range 15–930 nM.

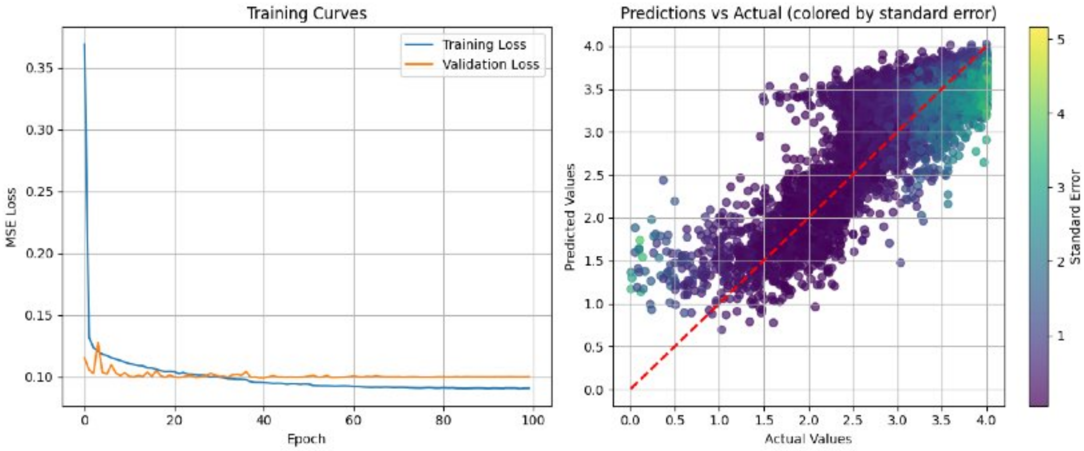


**Supplementary Figure 72 — Performance of five-position library CNN during Fold 1 of 5-fold cross-validation.**

Left: Training (blue) and validation (orange) MSE loss as a function of training epoch. Right: Correlation between predicted and actual (experimentally-measured) $\log_{10}(K_d)$ values in nM, colored by standard error. Performance metrics are reported in Supplementary Table 5. Red dashed line indicates the identity line.

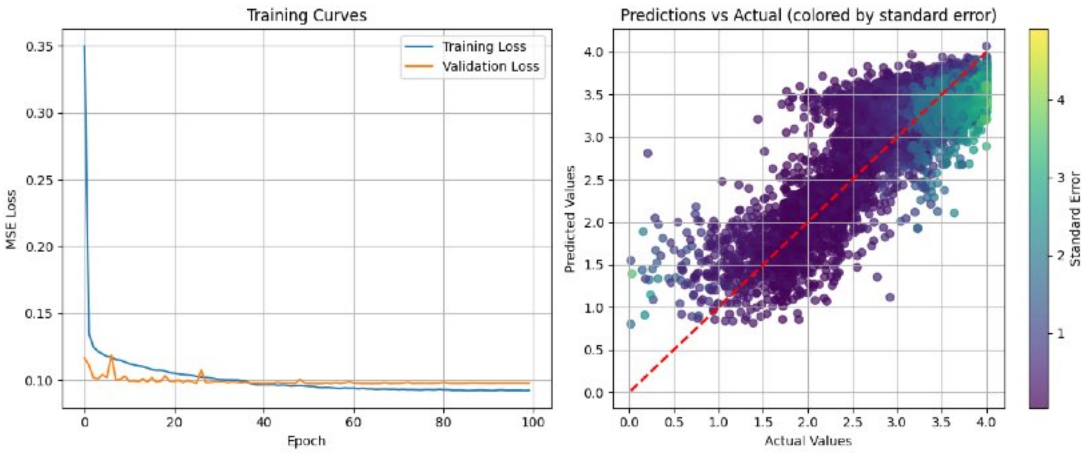


**Supplementary Figure 73 — Performance of five-position library CNN during Fold 2 of 5-fold cross-validation.** Left: Training (blue) and validation (orange) MSE loss as a function of training epoch. Right: Correlation between predicted and actual (experimentally-measured) $\log_{10}(K_d)$ values in nM, colored by standard error. Performance metrics are reported in Supplementary Table 5. Red dashed line indicates the identity line.

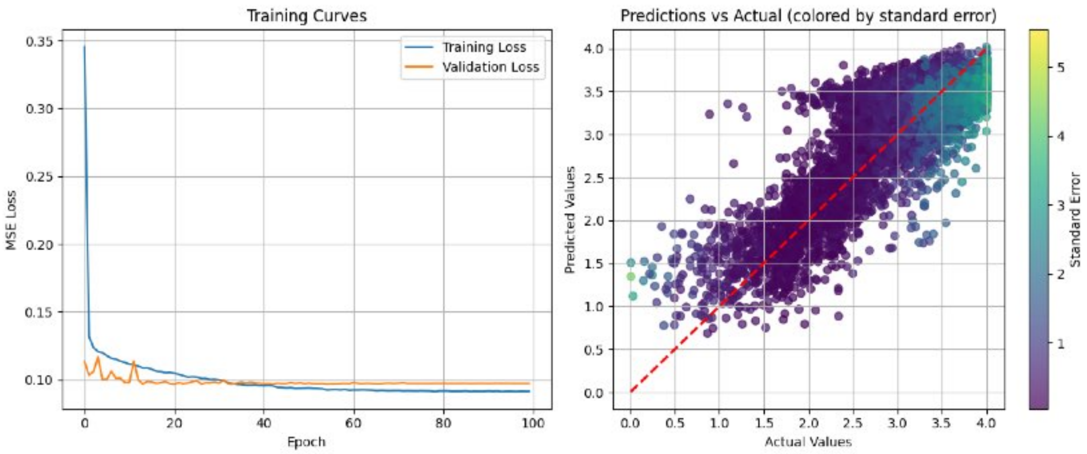


**Supplementary Figure 74 — Performance of five-position library CNN during Fold 3 of 5-fold cross-validation.** Left: Training (blue) and validation (orange) MSE loss as a function of training epoch. Right: Correlation between predicted and actual (experimentally-measured) $\log_{10}(K_d)$ values in nM, colored by standard error. Performance metrics are reported in Supplementary Table 5. Red dashed line indicates the identity line.

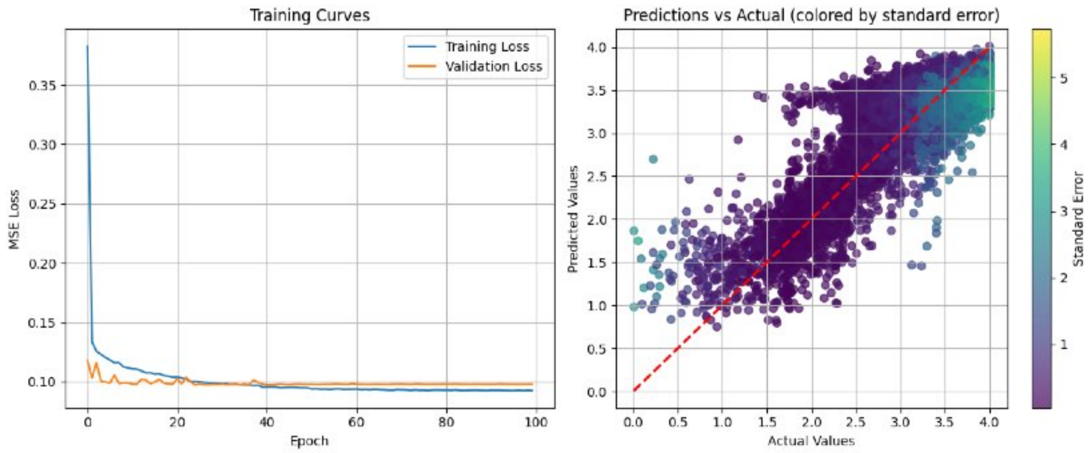


**Supplementary Figure 75 — Performance of five-position library CNN during Fold 4 of 5-fold cross-validation.** Left: Training (blue) and validation (orange) MSE loss as a function of training epoch. Right: Correlation between predicted and actual (experimentally-measured) $\log_{10}(K_d)$ values in nM, colored by standard error. Performance metrics are reported in Supplementary Table 5. Red dashed line indicates the identity line.

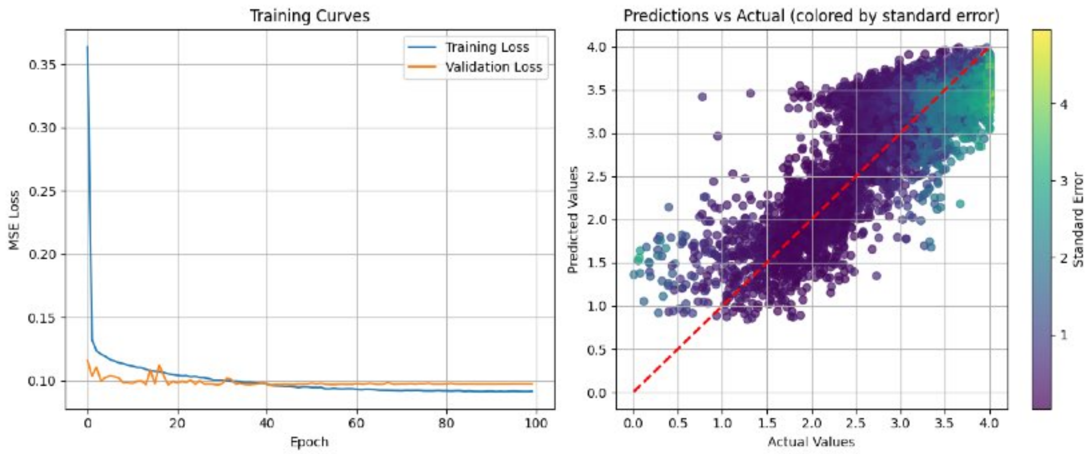


**Supplementary Figure 76 — Performance of five-position library CNN during Fold 5 of 5-fold cross-validation.** Left: Training (blue) and validation (orange) MSE loss as a function of training epoch. Right: Correlation between predicted and actual (experimentally-measured) $\log_{10}(K_d)$ values in nM, colored by standard error. Performance metrics are reported in Supplementary Table 5. Red dashed line indicates the identity line.

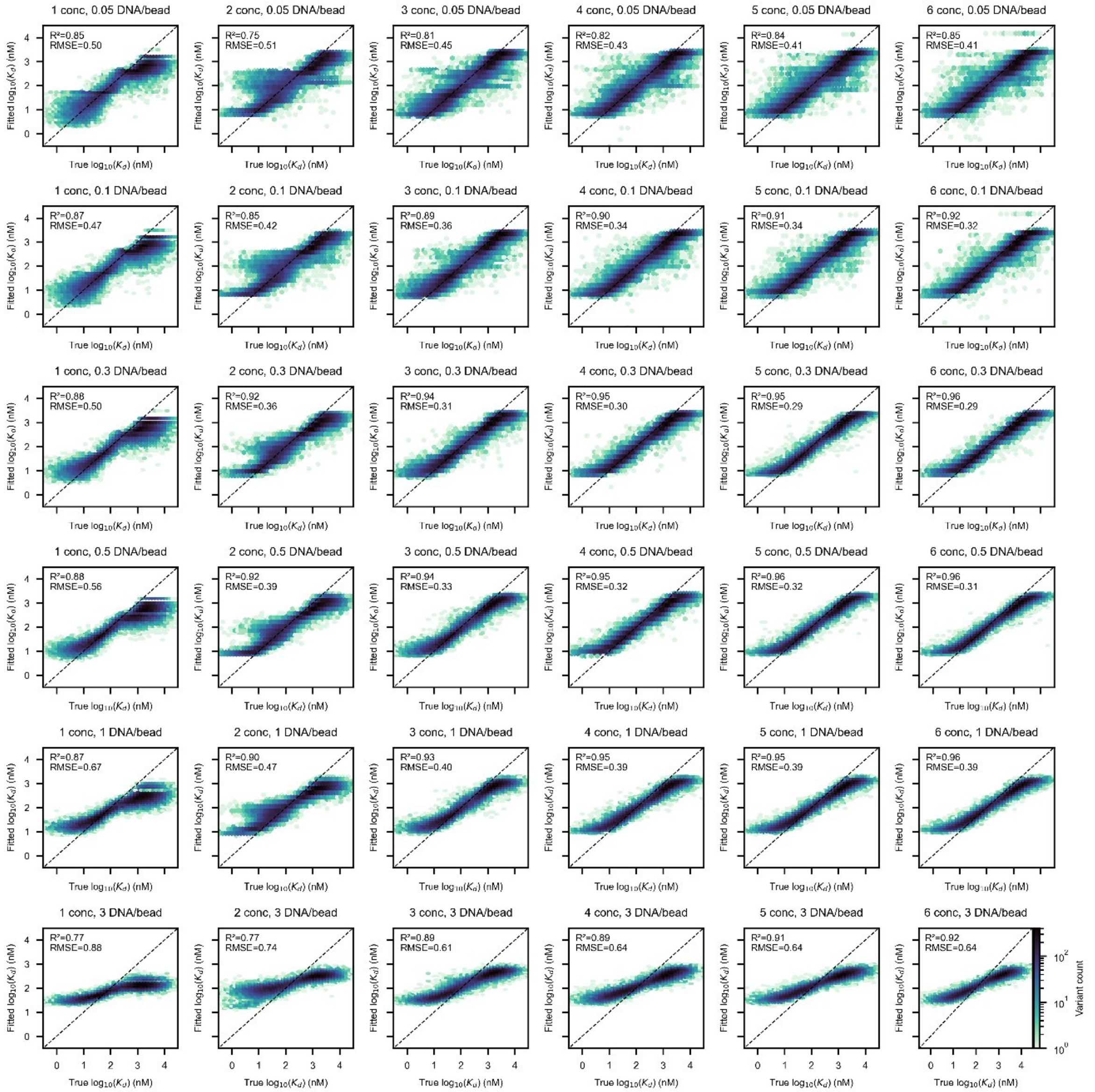


**Supplementary Figure 77 — Correlation between returned fitted and "true" input $\log_{10}(K_d)$ values from APB-TiteSeq binding simulations as a function of the number of ligand concentrations and mean DNA/bead parameters for a representative library.**

Hexbin plots showing returned "fitted" $\log_{10}(K_d)$ values versus "true" input $\log_{10}(K_d)$ values for a simulated representative deep mutational scanning library as a function of number of ligand concentration and mean DNA/bead parameters. Hexbins are colored by the number of variants with a given affinity range (log scale).

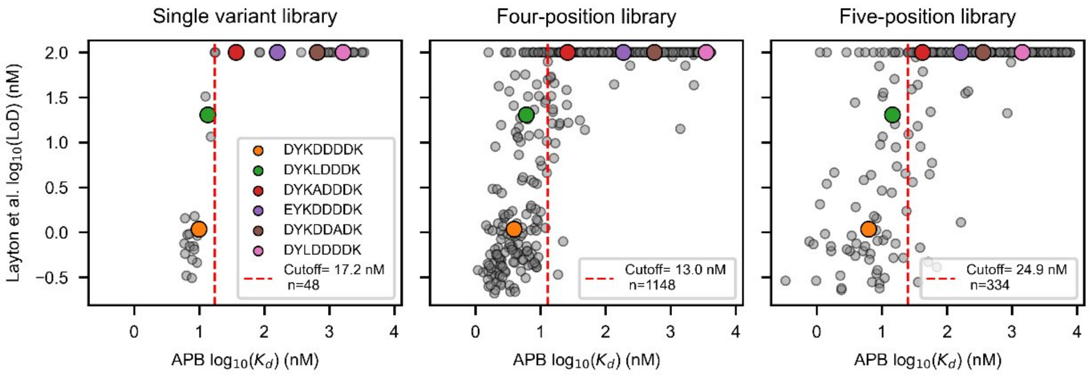


**Supplementary Figure 78 — Comparison between Prot-MaP Limit of Detection (LoD) values APB-TiteSeq-measured $K_d$ values.**

Prot-Map [50] $\log_{10}$(LoD) versus APB-TiteSeq $\log_{10}(K_d)$ values for FLAG variants shared between each APB library and the Prot-MaP dataset (single-variant, n = 48; four-position, n = 1,148; five-position, n = 334). Calibration variants with orthogonal FP measurements are shown as colored points; all other shared variants are shown in gray. Variants reported as "above detection limit" for Prot-MaP (LoD > 100 nM) were assigned LoD = 100 nM. The red dashed vertical line marks the Prot-MaP dynamic range boundary, defined as the APB-TiteSeq $K_d$ at which ≥ 50% of variants within a rolling window (maximum of 20 variants or 5% of the overlap population, centered) have LoD = 100 nM; this represents the APB-TiteSeq $K_d$ above which Prot-MaP no longer resolves differences in binding affinity.

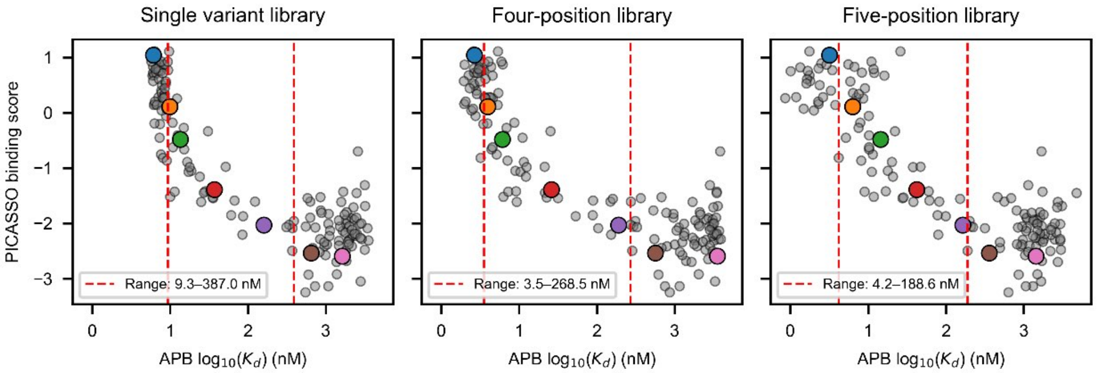


**Supplementary Figure 79 — Comparison of APB-TiteSeq-measured $K_d$ values with PICASSO Cas9-Display binding scores.**

PICASSO binding scores versus APB-TiteSeq $\log_{10}(K_d)$ values for FLAG variants shared between each APB library and the single-variant PICASSO [52] dataset (single-variant, n = 152; four-position, n = 153; five-position, n = 153). Variants that were also measured via FP are shown as colored points (blue: DYKDDDDW, orange: DYKDDDDK, green: DYKLDDDK, red: DYKADDDK, purple: EYKDDDDK, brown: DYKDDADK, pink: DYLDDDDK); all other shared variants are shown in gray. Red dashed vertical lines mark the dynamic range of the PICASSO assay, defined as the APB-TiteSeq $K_d$ interval over which the absolute Spearman correlation between APB-TiteSeq $\log_{10}(K_d)$ and PICASSO score remains ≥ 0.5 within a rolling window (maximum of 10 variants or 15% of the overlap population, centered). Within this range, PICASSO scores meaningfully track binding affinity as measured by APB-TiteSeq.

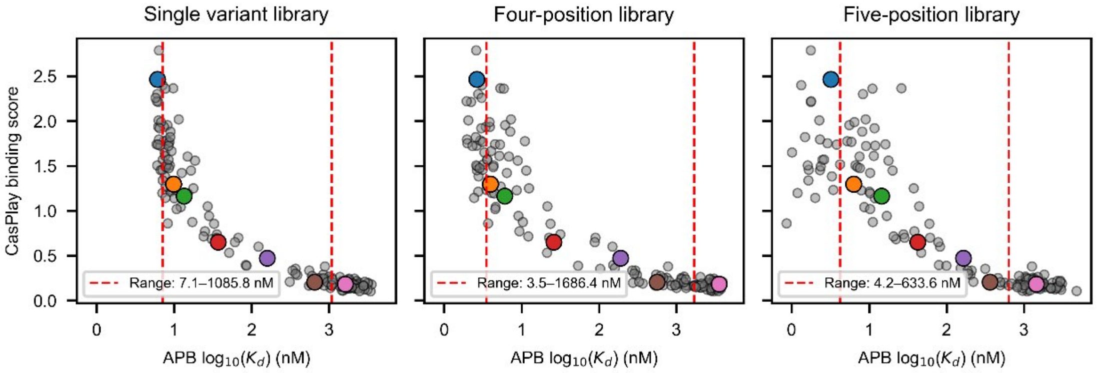


**Supplementary Figure 80 — Comparison of APB-TiteSeq-measured $K_d$ values with CasPlay Cas9-Display binding scores.**

CasPlay binding scores versus APB-TiteSeq $\log_{10}(K_d)$ values for FLAG variants shared between each APB library and the single-variant CasPlay [49] dataset (single-variant, n = 152; four-position, n = 153; five-position, n = 153). Variants that were also measured via FP are shown as colored points (blue: DYKDDDDW, orange: DYKDDDDK, green: DYKLDDDK, red: DYKADDDK, purple: EYKDDDDK, brown: DYKDDADK, pink: DYLDDDDK); all other shared variants are shown in gray. Red dashed vertical lines indicate the lower and upper clipping bounds of each APB measurement. Red dashed vertical lines mark the dynamic range of the CasPlay assay, defined as the APB $K_d$ interval over which the absolute Spearman correlation between APB $\log_{10}(K_d)$ and CasPlay score remains $\geq 0.5$ within a rolling window (maximum of 10 variants or 15% of the overlap population, centered). Within this range, CasPlay scores meaningfully track binding affinity as measured by APB-TiteSeq.

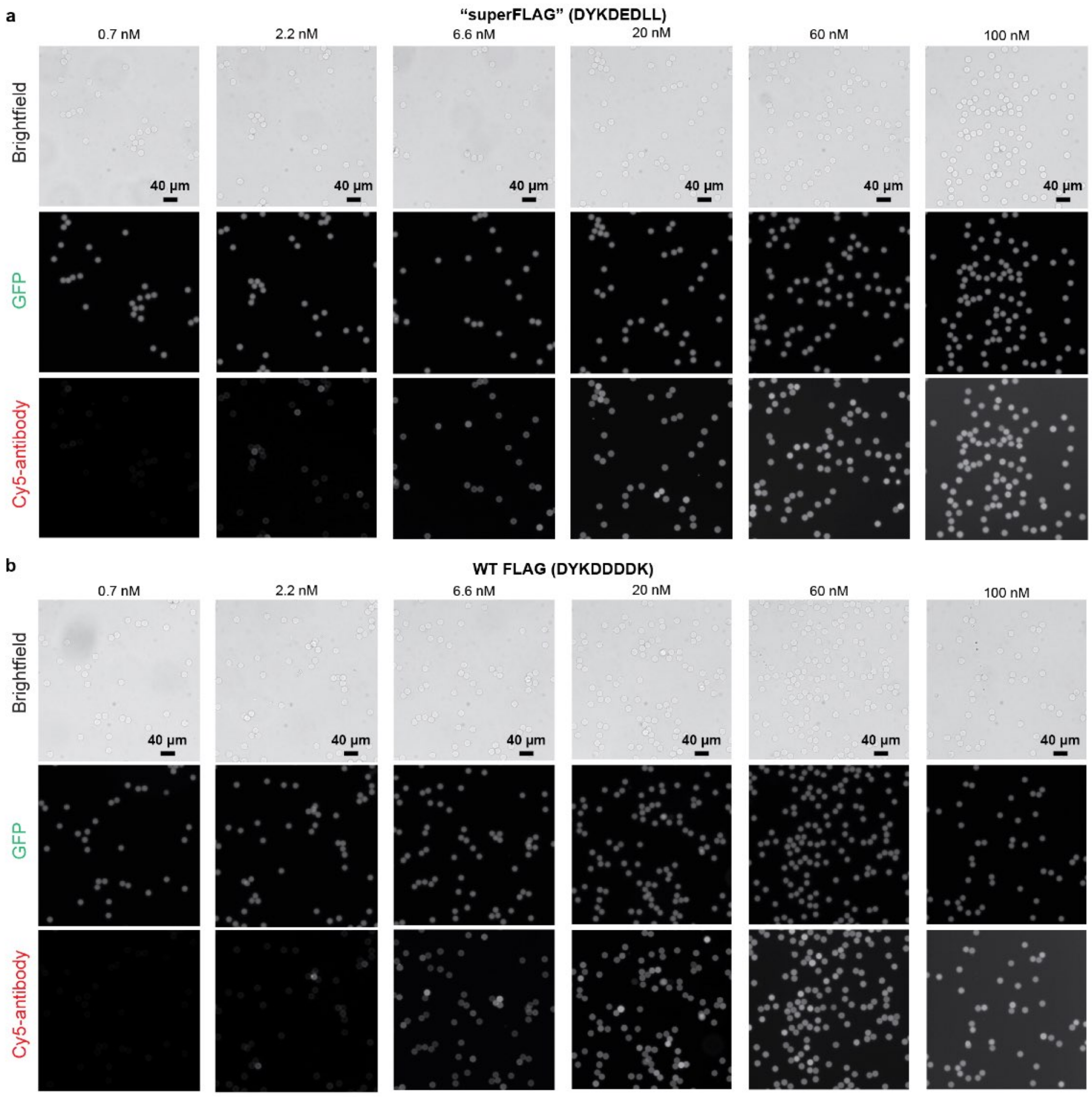


**Supplementary Figure 81 — Microscopy images of concentration-dependent antibody binding for “superFLAG” and WT FLAG variants.**

(a) Microscopy images of SNAP-DYKDEDLL-eGFP (“superFLAG”) immobilized on BG-coated beads after overnight incubation with increasing concentrations of M2 primary and Cy5 secondary antibodies. Beads were diluted 10x in buffer prior to imaging. Top row: bright field, middle row: GFP (100 ms), middle row: Cy5 (100 ms) fluorescence channels. Scale bar = 40 µm. (b) Microscopy images of SNAP-DYKDDDDK-eGFP (“WT FLAG”) immobilized on BG-coated beads after overnight incubation with increasing concentrations of M2 primary and Cy5 secondary antibodies. Beads were diluted 10x in buffer prior to imaging. Top row: bright field, middle row: GFP (100 ms), middle row: Cy5 (100 ms) fluorescence channels. Scale bar = 40 µm.

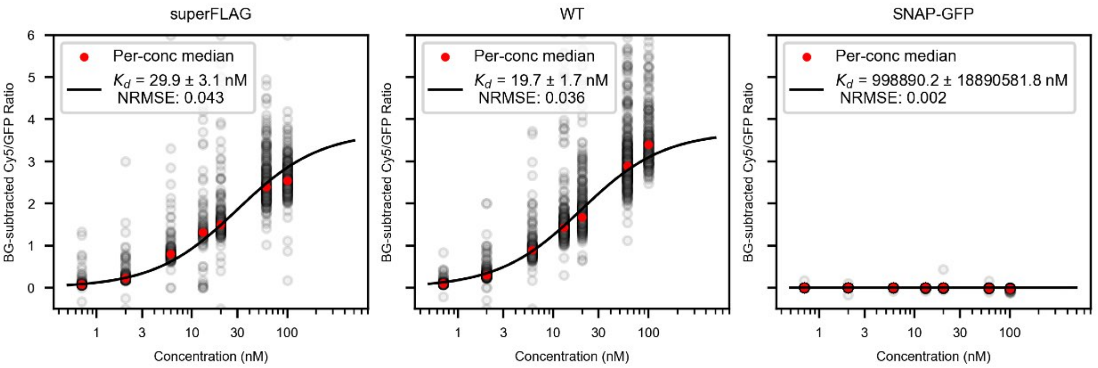


**Supplementary Figure 82 — Microscopy-generated binding curves for "superFLAG", WT FLAG, and no FLAG variants.**

Cy5/GFP intensities from microscopy images as a function of M2 antibody concentration (shared global $R_{\text{max}}$ = 3.7) for SNAP-DYKDEDLL-eGFP ("superFLAG"; left), SNAP-DYKDDDDK-eGFP ("WT FLAG"; center), and SNAP-eGFP ("SNAP-GFP"; left). Gray markers indicate background-subtracted Cy5/GFP intensities for individual beads (subset shown in Supplementary Figure 81); red markers indicate per-concentration median intensities; black line indicates the Langmuir isotherm fit yielding the annotated fitted $K_d$ value. No variants reached saturation at the highest staining concentration used for this experiment (100 nM). As a result, the global $R_{\text{max}}$ is under-determined and estimated affinities could be biased from true values; however, the relative difference between "superFLAG" and WT would be unaffected.

### Supplementary Tables

**Supplementary Table 1 — Sequencing run summary for APB experiments.**

Summary of sequencing metrics across five APB experiments, including library size, amplicon length, sequencing cost, read counts at each processing stage, and final filtered read counts for each library of interest.

| | **DRP215** | **DRP232** | **DRP236** | **DRP236** | **DRP221** |
|---|---|---|---|---|---|
| *Experiment parameters* | | | | | |
| Assay type | APB-Display | APB-TiteSeq | APB-TiteSeq | APB-TiteSeq | APB-SortSeq |
| Library size (expected variants) | 25 | 153 | 20,000 | 20,000 | 150,000 |
| Amplicon length (bp) | 262 | 262 | 262 | 262 | 262 |
| Sequencing requested (Gb) | 1 | 3 | 12 | 12 | 20 |
| Sequencing cost ($) | 500 | 600 | 1,050 | 1,050 | 1,450 |
| *Read counts* | | | | | |
| Total reads | 4,174,509 | 11,230,568 | 44,233,747 | 43,420,933 | 75,093,033 |
| Reads with variant region | 2,336,630 | 5,913,142 | 26,177,424 | 24,950,148 | 38,997,223 |
| Barcode-demultiplexed reads | 267,903 | 4,241,696 | 17,361,017 | 18,637,668 | 23,006,422 |
| *Library of interest* | | | | | |
| Fraction allotted | 0.75 | 0.50 | 0.86 | 0.86 | 0.95 |
| Expected reads | 200,927 | 2,120,848 | 14,930,475 | 16,028,394 | 21,856,101 |
| Actual reads | 206,597 | 1,486,610 | 14,795,957 | 16,023,892 | 21,568,417 |
| Filtered reads (e.g. expected variants) | 180,895 | 1,452,407 | 14,414,229 | 15,614,017 | 21,426,183 |

**Supplementary Table 2 — Simulation input parameters for each experimental dataset.**

Input parameters for APB-TiteSeq and APB-SortSeq simulations calibrated to each experimental dataset. *Assay type*: APB-TiteSeq measures binding across multiple antibody concentrations; APB-SortSeq uses a single concentration with orthogonally-validated calibration variants. $n_{\text{variants}}$: number of unique variants in the simulated library. $n_{\text{beads}}$: total beads loaded into emulsion PCR. $\lambda_{\text{DNA}}$: mean Poisson loading of DNA templates per bead during emulsion PCR. $p_{\text{double}}$: probability of co-encapsulating two beads in emulsion IVTT. $\sigma_{\text{Ab}}$: standard deviation of log-normal noise applied to the simulated antibody signal (Cy5/GFP ratio). Reads/bead: ratio of sequencing reads to occupied beads. $R_{\text{max}}$ strategy and value: method and fixed value for determining the saturation signal in $K_d$ fitting (*fixed* sets $R_{\text{max}} = 1$ in normalized signal units; not applicable for APB-SortSeq). Fitting mode: *calibrated* fits $K_d$ from a single concentration using calibration variants (APB-SortSeq only). $n_{\text{cal}}$: number of calibration variants. $K_d$ range: range of true $K_d$ values (nM) log-normally sampled for simulated library variants. Freq. skew: ratio of the 90th and 10th most frequent variants in the input library. Concentrations: antibody concentrations (nM) used in FACS simulation. Fitted $R_{\text{max}}$: experimental saturation signal (Cy5/GFP ratio units) derived from global Langmuir isotherm fits used for signal normalization. Replicates: number of independent simulation runs.

| | **DRP232** | **DRP236** | **DRP237** | **DRP221** |
|---|---|---|---|---|
| Assay type | APB-TiteSeq | APB-TiteSeq | APB-TiteSeq | APB-SortSeq |
| $n_{\text{variants}}$ | 153 | 20,000 | 20,000 | 145,000 |
| $n_{\text{beads}}$ | 300,000 | 3,000,000 | 3,100,000 | 50,000,000 |
| $\lambda_{\text{DNA}}$ | 0.30 | 0.10 | 0.10 | 0.03 |
| $p_{\text{double}}$ | 0.10 | 0.10 | 0.10 | 0.10 |
| $\sigma_{\text{Ab}}$ | 0.35 | 0.35 | 0.35 | 0.35 |
| Reads/bead | 2.5 | 5.0 | 5.0 | 6.0 |
| $R_{\text{max}}$ strategy | fixed | fixed | fixed | — |
| $R_{\text{max}}$ value | 1.0 | 1.0 | 1.0 | — |
| Fitting mode | — | — | — | calibrated |
| $n_{\text{cal}}$ | — | — | — | 7 |
| $K_d$ range (nM) | [0.5, 50,000] | [0.5, 50,000] | [0.5, 50,000] | [0.5, 50,000] |
| Freq. skew | 1.6 | 10.2 | 10.2 | 5.6 |
| Concentrations (nM) | 0.6, 2, 6, 20, 60, 200, 400 | 0.6, 6, 20, 60, 200, 400 | 0.6, 6, 20, 60, 200, 400 | 60 |
| Fitted $R_{\text{max}}$ | 17.92 | 10.78 | 6.49 | 20.92 |
| Replicates | 10 | 3 | 3 | 1 |

**Supplementary Table 3 — Per-concentration simulation parameters for each experimental dataset.**

Per-concentration FACS gate boundaries and blank-bead background binding parameters used in simulations for each experimental dataset. Gate upper bounds ($g_1$–$g_4$) are the background-subtracted Cy5/GFP signal thresholds separating the four sort bins, normalized by the fitted $R_{\max}$ of each experiment; values >1 indicate gates extending above the expected saturation signal. Background binding parameters describe the residual non-specific Cy5/GFP signal on blank beads after median background subtraction, modeled as a log-normal distribution: $\tilde{r}_{\text{bg}}$, median background ratio used for background subtraction; $\sigma_{\text{bg}}$, log-normal shape parameter; $s_{\text{bg}}$, log-normal scale parameter ($e^{\mu}$, where $\mu$ is the log-scale mean).

| | Conc. (nM) | $g_1$ | $g_2$ | $g_3$ | $g_4$ | $\tilde{r}_{\text{bg}}$ | $\sigma_{\text{bg}}$ | $s_{\text{bg}}$ |
|---|---|---|---|---|---|---|---|---|
| DRP232 | 0.6 | 0.012 | 0.054 | 0.110 | 0.557 | 0.028 | 3.074 | 0.008 |
| | 2.0 | 0.023 | 0.107 | 0.330 | 1.112 | 0.076 | 2.639 | 0.044 |
| | 6.0 | 0.034 | 0.160 | 0.495 | 1.388 | 0.140 | 2.106 | 0.117 |
| | 20.0 | 0.037 | 0.316 | 0.818 | 1.935 | 0.339 | 1.185 | 0.391 |
| | 60.0 | 0.022 | 0.245 | 0.915 | 2.199 | 0.606 | 0.929 | 0.709 |
| | 200.0 | 0.147 | 0.426 | 1.152 | 2.714 | 1.369 | 0.803 | 1.568 |
| | 400.0 | 0.109 | 0.444 | 1.003 | 2.677 | 2.040 | 0.842 | 2.339 |
| DRP236 | 0.6 | 0.012 | 0.045 | 0.184 | 0.926 | 0.018 | 2.336 | 0.008 |
| | 6.0 | 0.040 | 0.133 | 0.550 | 1.385 | 0.070 | 1.207 | 0.069 |
| | 20.0 | 0.038 | 0.131 | 0.734 | 1.383 | 0.086 | 1.046 | 0.091 |
| | 60.0 | 0.074 | 0.260 | 0.909 | 1.837 | 0.204 | 0.890 | 0.228 |
| | 200.0 | 0.152 | 0.338 | 1.080 | 2.749 | 0.358 | 0.854 | 0.417 |
| | 400.0 | 0.140 | 0.326 | 1.068 | 2.737 | 0.493 | 0.855 | 0.570 |
| DRP237 | 0.6 | 0.022 | 0.062 | 0.231 | 4.008 | 0.001 | 2.407 | 0.001 |
| | 6.0 | 0.020 | 0.091 | 0.461 | 4.007 | 0.007 | 2.461 | 0.003 |
| | 20.0 | 0.020 | 0.091 | 0.538 | 4.007 | 0.014 | 2.318 | 0.008 |
| | 60.0 | 0.036 | 0.150 | 0.612 | 4.003 | 0.034 | 1.615 | 0.032 |
| | 200.0 | 0.094 | 0.233 | 0.911 | 3.994 | 0.095 | 1.225 | 0.107 |
| | 400.0 | 0.134 | 0.288 | 0.905 | 3.988 | 0.130 | 1.199 | 0.147 |
| DRP221 | 60.0 | 0.176 | 0.415 | 0.845 | 2.853 | 0.313 | 0.986 | 0.303 |

**Supplementary Table 4 — Four-position CNN cross-validation results.**

Summary of 5-fold cross-validation performance metrics for the trained four-position CNN, including mean squared error (MSE), mean absolute error (MAE), and coefficient of determination ($R^2$) for each fold. Ensemble $R^2$ reflects the performance of the full model trained on all data.

| | **Fold 1** | **Fold 2** | **Fold 3** | **Fold 4** | **Fold 5** | **Average** | **Ensemble** |
|---|---|---|---|---|---|---|---|
| MSE | 0.0344 | 0.0353 | 0.0358 | 0.0357 | 0.0350 | | |
| MAE | 0.1338 | 0.1378 | 0.1347 | 0.1372 | 0.1337 | | |
| $R^2$ | 0.9381 | 0.9377 | 0.9413 | 0.9389 | 0.9392 | 0.9390 | 0.962 |

**Supplementary Table 5 — Five-position CNN cross-validation results.**

Summary of 5-fold cross-validation performance metrics for the trained five-position CNN, including mean squared error (MSE), mean absolute error (MAE), and coefficient of determination ($R^2$) for each fold. Ensemble $R^2$ reflects the performance of the full model trained on all data.

| | **Fold 1** | **Fold 2** | **Fold 3** | **Fold 4** | **Fold 5** | **Average** | **Ensemble** |
|---|---|---|---|---|---|---|---|
| MSE | 0.100 | 0.098 | 0.097 | 0.098 | 0.097 | | |
| MAE | 0.236 | 0.236 | 0.233 | 0.234 | 0.234 | | |
| $R^2$ | 0.707 | 0.698 | 0.703 | 0.706 | 0.704 | 0.704 | 0.739 |

**Supplementary Table 6 — Cost analysis for APB experiments.**

Itemized reagent costs across five APB experiments, including unit costs and per-experiment totals. Costs are summarized by experimental stage. Cost-per-variant metrics are reported for both expected and measured library sizes, and normalized by number of ligand concentrations where applicable.

| **Reagent** | **Unit cost** | **Units** | **DRP215** | **DRP232** | **DRP236** | **DRP237** | **DRP221** |
|---|---|---|---|---|---|---|---|
| *Experiment parameters* | | | | | | | |
| Assay type | | | APB-Display | APB-TiteSeq | APB-TiteSeq | APB-TiteSeq | APB-SortSeq |
| Library size (expected variants) | | | 25 | 153 | 20,000 | 20,000 | 150,000 |
| Library size (measured variants) | | | 25 | 152 | 18,000 | 18,000 | 88,000 |
| No. ligand concentrations | | | 1 | 7 | 6 | 6 | 1 |
| *Cloning (~2 days, intermittent)* | | | | | | | |
| DNA synthesis | | | $121.00 | $682.00 | $191.00 | $— | $70.00 |
| Golden Gate assembly (BsaI) | $7.12 | $/µL enzyme | $7.12 | $7.12 | $7.12 | $— | $7.12 |
| PCR mix | $1.28 | $/µL enzyme | $0.64 | $0.64 | $0.64 | $— | $0.64 |
| Gel purification columns | $2.32 | $/column | $2.32 | $2.32 | $2.32 | $— | $2.32 |
| *Template dilution check (2 hours)* | | | | | | | |
| PCR mix | $1.28 | $/µL enzyme | $2.82 | $1.69 | $2.12 | $— | $2.82 |
| BioRad oil | $0.01 | $/µL | $1.20 | $0.45 | $2.40 | $— | $1.20 |
| *APB-Display: emulsion PCR (3 hours)* | | | | | | | |
| Oligo/BG functionalized beads | $0.16 | $/µL beads | $3.26 | $21.19 | $78.24 | $78.24 | $163.00 |
| PCR mix | $1.28 | $/µL enzyme | $3.95 | $10.13 | $18.46 | $18.46 | $98.71 |
| BioRad oil | $0.01 | $/µL | $0.30 | $1.80 | $7.20 | $7.20 | $15.00 |
| Perfluorooctanol | $0.01 | $/µL | $0.22 | $3.30 | $4.40 | $4.40 | $27.50 |
| *APB-Display: emulsion IVTT (3 hours)* | | | | | | | |
| PURExpress (A+B, no discount) | $1.59 | $/µL | $33.29 | $190.20 | $665.70 | $665.70 | $1,664.25 |
| RNase Inhibitor | $1.04 | $/µL | $1.25 | $7.27 | $24.94 | $24.94 | $51.95 |
| BioRad oil | $0.01 | $/µL | $0.35 | $1.95 | $7.80 | $7.80 | $16.25 |
| Perfluorooctanol | $0.01 | $/µL | $0.33 | $2.75 | $5.50 | $5.50 | $27.50 |
| SNAP-SurfaceBlock (4 mM) | $2.79 | $/µL | $16.72 | $34.83 | $34.83 | $34.83 | $41.79 |
| *Staining (overnight)* | | | | | | | |
| M2 anti-FLAG (1 mg/mL) | $2.48 | $/µL | $1.24 | $44.55 | $44.55 | $44.55 | $74.25 |
| Cy5 secondary (2 mg/mL) | $0.59 | $/µL | $0.30 | $5.35 | $5.35 | $5.35 | $14.85 |
| *FACS (3-9 hours)* | | | | | | | |
| FACS time | $60.00 | $/hour | $180.00 | $240.00 | $540.00 | $540.00 | $540.00 |
| *Sequencing (4 hour prep)* | | | | | | | |
| PCR mix | $1.28 | $/µL enzyme | $108.33 | $15.77 | $14.10 | $14.10 | $2.26 |
| KAPA beads | $0.02 | $/µL | $— | $25.20 | $21.24 | $21.24 | $3.24 |
| Qubit reagents | $0.00 | $/µL | $3.20 | $25.60 | $22.00 | $22.00 | $4.00 |
| Gel purification columns | $2.32 | $/column | $6.96 | $— | $— | $— | $— |
| Plasmidsaurus sequencing | | | $500.00 | $600.00 | $1,050.00 | $1,050.00 | $1,450.00 |
| **Total** | | | **$994.78** | **$1,924.10** | **$2,749.90** | **$2,544.30** | **$4,278.65** |
| *Cost per variant* | | | | | | | |
| Per variant (expected library) | | | $39.79 | $12.58 | $0.12 | $0.11 | $0.03 |
| Per variant (measured library) | | | $39.79 | $12.66 | $0.15 | $0.14 | $0.05 |
| Per variant, per conc. (expected) | | | $39.79 | $1.80 | $0.02 | $0.02 | $0.03 |
| Per variant, per conc. (measured) | | | $39.79 | $1.81 | $0.03 | $0.02 | $0.05 |

**Supplementary Table 7 — FLAG library sequences used in APB experiments.**

Sequences of all FLAG variant libraries used in APB experiments. Sequences are listed in the 5′ → 3′ direction and written as space-separated codons. The WT FLAG sequence is `gac tac aag gac gac gac gac aag` (positions 1–8). For single-substitution libraries, each row describes all 8 variants for a given amino acid substitution; the mutated codon replaces the corresponding WT codon at positions 1–8 while all other positions remain as WT. NNK denotes a degenerate codon (N = any base; K = G or T). RVK denotes a restricted degenerate codon (R = A or G; V = A, C, or G; K = G or T). NNC denotes a restricted degenerate codon (N = any base; C = C only).

| **Variants** | **Sequence pattern (5′ → 3′)** |
|---|---|
| *FLAG 25-variant library* | |
| flag_wt | gac tac aag gac gac gac gac aag |
| flag_A1–A8 | GCG substituted at positions 1–8 |
| flag_L1–L8 | CTG substituted at positions 1–8 |
| flag_E1–E8 | GAA substituted at positions 1–8 |
| *FLAG single-variant library* | |
| flag_A1–A8 | GCG substituted at positions 1–8 |
| flag_C1–C8 | TGC substituted at positions 1–8 |
| flag_D1–D8 | GAT substituted at positions 1–8 |
| flag_E1–E8 | GAA substituted at positions 1–8 |
| flag_F1–F8 | TTC substituted at positions 1–8 |
| flag_G1–G8 | GGT substituted at positions 1–8 |
| flag_H1–H8 | CAT substituted at positions 1–8 |
| flag_I1–I8 | ATT substituted at positions 1–8 |
| flag_K1–K8 | AAA substituted at positions 1–8 |
| flag_L1–L8 | CTG substituted at positions 1–8 |
| flag_M1–M8 | ATG substituted at positions 1–8 |
| flag_N1–N8 | AAC substituted at positions 1–8 |
| flag_P1–P8 | CCG substituted at positions 1–8 |
| flag_Q1–Q8 | CAG substituted at positions 1–8 |
| flag_R1–R8 | CGT substituted at positions 1–8 |
| flag_S1–S8 | AGC substituted at positions 1–8 |
| flag_T1–T8 | ACC substituted at positions 1–8 |
| flag_V1–V8 | GTT substituted at positions 1–8 |
| flag_W1–W8 | TGG substituted at positions 1–8 |
| flag_Y1–Y8 | TAT substituted at positions 1–8 |
| *FLAG double mutant library* | |
| flag_2pos_$i$_$j$ ($i < j$, positions 1–8) | NNK substituted at positions $i$ and $j$; all 28 pairwise combinations included |
| *FLAG four position library* | |
| flag_4pos (oligo: gDRP-016) | RVK tac aag NNC RVK gac RVK aag |
| *FLAG five position library* | |
| flag_5pos (oligo: gDRP-017) | RVK tac aag NNC RVK gac RVK NNC |

**Supplementary Table 8 — Primers used in APB experiments.**

Sequences of oligonucleotide primers used for emulsion PCR and evSeq barcoding. Sequences are listed in the $5' \rightarrow 3'$ direction. The acrydite modification (/5Acryd/) enables covalent attachment of the reverse primer to polyacrylamide beads during bead generation.

| **Name** | **Purpose** | **Sequence ($5' \rightarrow 3'$)** |
|---|---|---|
| primDRP-039_PCRfwd | Forward primer for emulsion PCR | CCACCTGACGTCTAAAGAAACC |
| primDRP-047_Acryd_IVTTrev | Acrydite reverse primer for bead attachment during emulsion PCR | /5Acryd/ACCGTATTACCGCCTTTGAGTGAG |
| primDRP-075_evSeq_SMT213fwd | Forward primer for extracting DNA from beads for evSeq barcoding | CACCCAAGACCACTCTCCGG<br>CATGAAGGTCATCGTCTGGGTAAAC |
| primDRP-076_evSeq_SMT213rev | Reverse primer for extracting DNA from beads for evSeq barcoding | CGGTGTGCGAAGTAGGTGC<br>TGAACAGCTCCTCGCCCTTG |

**Supplementary Table 9 — Amplicon sequences used in APB experiments.**

Full sequences of amplicons used in APB experiments, listed in the $5' \rightarrow 3'$ direction.

| **Name** | **Purpose** | **Sequence ($5' \rightarrow 3'$)** |
|---|---|---|
| SMT213-FLAGwt full amplicon | Amplicon containing T7 promoter, SNAP, GS linker, WT FLAG, GS linker, eGFP, and terminator | CCACCTGACGTCTAAAGAAACCGAATAATACGACTCACTATAGGGCTTAAGTA<br>TAAGGAGATATACATATGGACAAAGATTGCGAAATGAAACGTACCACCCTGGA<br>TAGCCCGCTGGGCAAACTGGAACTGAGCGGCTGCGAACAGGGCCTGCATGAAA<br>TTAAACTGCTGGGTAAAGGCACCAGCGCGGCCGATGCGGTTGAAGTTCCGGCC<br>CCGGCCGCCGTGCTGGGTGGTCCGGAACCGCTGATGCAGGCGACCGCGTGGCTG<br>AACGCGTATTTTCATCAGCCGGAAGCGATTGAAGAATTTCCGGTTCCGGCGCTG<br>CATCATCCGGTGTTTCAGCAGGAGAGCTTTACCCGTCAGGTGCTGTGGAAACTG<br>CTGAAAGTGGTTAAATTTGGCGAAGTGATTAGCTATCAGCAGCTGGCGGCCCTG<br>GCGGGTAATCCGGCGGCCACCGCCGCCGTTAAAACCGCGCTGAGCGGTAACCCG<br>GTGCCGATTCTGATTCCGTGCCATCGTGTGGTTAGCTCTAGCGGTGCGGTTGGC<br>GGTTATGAAGGTGGTCTGGCGGTGAAAGAGTGGCTGCTGGCCCATGAAGGTCAT<br>CGTCTGGGTAAACCGGGTCTGGGAGGTGGAGGGTCTGGGGGAGGAGGCTCAGGC<br>GACTACAAGGACGACGACGACAAGGGTGGCGGTTCTGGCGGTGGCGGTAGTGGC<br>AGCAAGGGCGAGGAGCTGTTCACCGGGGTGGTGCCCATCCTGGTCGAGCTGGAC<br>GGCGACGTAAACGGCCACAAGTTCAGCGTGTCCGGCGAGGGCGAGGGCGATGCC<br>ACCTACGGCAAGCTGACCCTGAAGTTCATCTGCACCACCGGCAAGCTGCCCGTG<br>CCCTGGCCCACCCTCGTGACCACCCTGACCTACGGCGTGCAGTGCTTCAGCCGC<br>TACCCCGACCACATGAAGCAGCACGACTTCTTCAAGTCCGCCATGCCCGAAGGCT<br>ACGTCCAGGAGCGCACCATCTTCTTCAAGGACGACGGCAACTACAAGACCCGCG<br>CCGAGGTGAAGTTCGAGGGCGACACCCTGGTGAACCGCATCGAGCTGAAGGGCA<br>TCGACTTCAAGGAGGACGGCAACATCCTGGGGCACAAGCTGGAGTACAACTACA<br>ACAGCCACAACGTCTATATCATGGCCGACAAGCAGAAGAACGGCATCAAGGTGA<br>ACTTCAAGATCCGCCACAACATCGAGGACGGCAGCGTGCAGCTCGCCGACCACT<br>ACCAGCAGAACACCCCCATCGGCGACGGCCCCGTGCTGCTGCCCGACAACCACT<br>ACCTGAGCACCCAGTCCGCCCTGAGCAAAGACCCCAACGAGAAGCGCGATCACA<br>TGGTCCTGCTGGAGTTCGTGACCGCCGCCGGGATCACTCTCGGCATGGACGAGC<br>TGTACAAATAATGATTCCCGCTGATAGTGCTAGTGTAGATCGCTACTAGTAGCGG<br>CCGCTGCAGTCCGGCAAAAAAGGGCAAGGTGTCACCACCCTGCCCTTTTTCTTT<br>AAAACCGAAAAGATTACTTCGCGTTATGCAGGCTTCCTCGCTCACTGACTCGCT<br>GCGCTCGGTCGTTCGGCTGCGGCGAGCGGTATCAGCTCACTCAAAGGCGGTAAT<br>ACGGT |
| SMT213-FLAGwt evSeq amplicon | evSeq amplicon containing WT FLAG and representative forward and reverse evSeq barcodes | TCGTCGGCAGCGTCAGATGTGTATAAGAGACAGGATCATGCACCCAAGACC<br>ACTCTCCGGCATGAAGGTCATCGTCTGGGTAAACCGGGTCTGGGAGGTGGA<br>GGGTCTGGGGGAGGAGGCTCAGGCGACTACAAGGACGACGACGACAAGGGT<br>GGCGGTTCTGGCGGTGGCGGTAGTGGCAGCAAGGGCGAGGAGCTGTTCAGC<br>ACCTACTTCGCACACCGGCAGTTCCTGTCTCTTATACACATCTCCGAGCC<br>CACGAGAC |

### Supplementary Notes

#### *Supplementary Note 1: Simulation of APB-TiteSeq $K_d$ Estimation*

To characterize the accuracy and dynamic range of APB-TiteSeq as a function of experimental parameters, we implemented a computational simulation that recapitulates each physical step of the workflow, propagating stochastic noise at every stage.

**Library construction.** The simulation begins by constructing a virtual protein variant library of $N$ variants, each assigned a true dissociation constant $K_d$ drawn from a user-specified distribution (log-uniform or log-normal across a defined range, or from a mixture of components to model realistic deep mutational scanning libraries with bimodal binder/non-binder subpopulations). Each variant is also assigned a relative library frequency, which may be uniform or drawn from a log-normal distribution parameterized by a fold-skew factor; this captures the natural abundance variation present in real combinatorial libraries.

**Emulsion PCR loading.** DNA templates are loaded onto beads under Poisson statistics with a mean of $\lambda$ templates per bead. The total number of templates is drawn as Poisson($N_{\text{beads}} \times \lambda$); each template is then assigned independently to a random bead and a variant identity drawn in proportion to library frequencies. This produces a data structure with one entry per (bead, variant) pair, correctly representing multi-variant beads as multiple entries sharing a bead identifier. At the default $\lambda = 0.1$, approximately 9.5% of beads are singly occupied and ~0.5% carry two or more variants, closely matching experimental Poisson loading conditions.

**DNA amplification.** Following emulsion PCR, each (bead, variant) entry undergoes in-droplet PCR amplification. The amplified copy number is drawn from a log-normal distribution centered on template_count×amplification_factor, with multiplicative noise controlled by a log-space standard deviation $\sigma_{\text{amp}}$. This captures bead-to-bead variability in PCR efficiency, which propagates through all downstream steps as heterogeneity in the protein display level.

**Emulsion PCR breaking and DNA smearing.** During emulsion breaking, a low-probability cross-contamination event transfers a small fraction of amplified DNA from a donor bead to a randomly chosen recipient bead. This introduces rare off-identity DNA molecules that can generate protein signal from the wrong variant, creating a background that increases with sequencing depth.

**Protein expression via IVTT.** Protein copy number for each (bead, variant) entry is drawn from a log-normal distribution centered on amplified_dna_count × expression_factor, with log-space noise $\sigma_{\text{exp}}$. The aggregate GFP signal of a bead is the sum of protein copy counts across all its variants, reflecting the contribution of each expressed protein through the APB display scaffold.

**IVTT emulsion loading and double encapsulation.** Beads are re-encapsulated into IVTT droplets. A fraction of droplets receive two beads. When co-encapsulated beads carry different variants, expressed protein diffuses bidirectionally: each bead retains a fraction $(1 - f_{\text{smear}})$ of its own protein and acquires $f_{\text{smear}}$ of its partner's protein. Critically, migrated protein is assigned zero amplified DNA count, so it contributes Cy5 signal via antibody binding without contributing DNA to sequencing. This decouples the fluorescence readout from the sequencing identity and introduces a systematic source of $K_d$ estimation error for beads involved in double-encapsulation events.

**IVTT emulsion breaking and protein smearing.** Analogous to the ePCR smearing step, a low-probability event transfers a small fraction of expressed protein between beads during IVTT emulsion breaking.

**Antibody binding.** For each concentration in the titration series, bound antibody (Cy5) count is computed for each (bead, variant) entry as

$$\text{antibody\_count} = \text{prot\_count} \times \frac{[L]}{K_d + [L]}, \tag{9}$$

where $[L]$ is the free ligand concentration in nM and $K_d$ is the variant's true dissociation constant. A multiplicative log-normal noise term $\sigma_{\text{ab}}$ is applied per entry to model bead-to-bead variability in antibody labeling efficiency and fluorescence detection. The per-bead aggregate Cy5 signal is the sum of antibody counts across all variants on that bead, including any contributions from co-encapsulation cross-contamination.

**Non-specific antibody binding (optional).** To replicate the systematic background present at high staining concentrations in experimental data, the simulation optionally models non-specific antibody adsorption at the sorting step. In real experiments, blank beads (displaying no antigen) produce a concentration-dependent Cy5/GFP signal because antibody adsorbs non-specifically to the bead surface; this background is log-normally distributed across beads. Standard practice subtracts the per-concentration median blank-bead Cy5/GFP ratio from all samples before analysis, but because the log-normal distribution is right-skewed, median subtraction leaves a positive mean residual that inflates the apparent signal of weak binders and biases fitted $K_d$ values upward.

When this option is activated, the user provides a dataframe of pre-computed log-normal fit parameters (shape, location, and scale) and the empirical median of the blank-bead Cy5/GFP distribution at each antibody concentration, derived by fitting their experimental blank-bead data with a log-normal distribution (`scipy.stats.lognorm`) prior to simulation. During simulation, after computing each bead's specific Cy5/GFP ratio from antibody binding, an independent residual is drawn for each bead: a non-specific Cy5/GFP value is sampled from the fitted log-normal distribution for the matching concentration, the empirical median is subtracted (mirroring experimental background subtraction), and the result is divided by the experimental $R_{\text{max}}$ to convert to the simulation's normalized Cy5/GFP scale. This residual is added to the bead's specific ratio and clipped at zero before bin assignment. This step is used only when the simulation is matched to a specific real experiment.

**FACS sorting.** Beads are sorted into $n$ bins (default $n = 4$) based on their Cy5/GFP ratio. Beads below a minimum GFP threshold are excluded as non-expressing. Bin edges may be specified as (i) experimental gate boundaries imported directly from flow cytometry data, (ii) fixed values applied uniformly across concentrations, or (iii) adaptive per-concentration edges computed from the empirical distribution of Cy5/GFP ratios at that concentration. The adaptive strategy places equal fractions of beads into each bin by default, ensuring all bins are populated at every concentration; a top-heavy variant assigns a smaller fraction (default 10%) to the highest-signal bin to provide a tighter upper constraint on the binding isotherm, mirroring experimental gating practice.

**Amplicon recovery.** Within each sort bin, the pool of DNA available for sequencing is the sum of amplified DNA copy counts across all beads sorted into that bin, aggregated by variant. A log-normal amplification bias ($\sigma_{\mathrm{rec}}$) is applied independently per (variant, bin) pair to model unequal PCR efficiency during post-sort amplicon recovery.

**Sequencing simulation.** Sequencing reads for each (concentration, bin) pair are drawn by multinomial sampling from the amplicon pool proportions, after applying a per-variant log-normal sequencing bias ($\sigma_{\mathrm{seq}}$). The total read depth can be specified as a fixed number of reads per bin, a fixed total per concentration distributed proportionally to bin sizes, or a ratio of reads per sorted bead.

**Sequencing analysis and mean ratio calculation.** The simulated count data passes through the identical analysis pipeline used for experimental data. Read counts are normalized by the number of sorted beads per bin, and a mean Cy5/GFP ratio per variant per concentration is computed as the read-count-weighted average of the per-bin median Cy5/GFP ratios, with a pseudocount added to stabilize estimates for low-count variants.

**$K_d$ fitting.** For each variant, a non-cooperative 1:1 binding isotherm

$$\overline{R}([L]) = R_{\mathrm{max}} \frac{[L]}{K_d + [L]} \tag{10}$$

is fit to the mean Cy5/GFP ratio as a function of ligand concentration using non-linear least squares. $R_{\mathrm{max}}$ is estimated by a global two-stage procedure: $K_d$ and $R_{\mathrm{max}}$ are first fit jointly for all variants, and the 98th percentile of the resulting $R_{\mathrm{max}}$ estimates is adopted as a global $R_{\mathrm{max}}$; $K_d$ is then re-fit with $R_{\mathrm{max}}$ held fixed. This strategy is robust to outliers and avoids overfitting that arises from per-variant $R_{\mathrm{max}}$ estimation when the concentration range does not fully saturate tight binders.
